# scRIPT: a rapid, high-coverage single-cell lineage-tracing method applied to the hematoendothelial differentiation of embryonic stem cells

**DOI:** 10.64898/2026.10.14.759042

**Authors:** Ridvan Cetin, Eric Bindels, Jente van Staalduinen, Lucas Kuijpers, Giulia Picco, Mathijs A. Sanders, Remco M. Hoogenboezem, Gregory van Beek, Joost Gribnau, Eskeatnaf Mulugeta, Danny Huylebroeck, Frank Grosveld

## Abstract

Single-cell lineage tracing, which often involves CRISPR-Cas-based barcode insertion followed by novel barcode creation, links a living cell’s transcriptome to its clonal history. It is often less suitable for short cell differentiation protocols: the barcoding time window does not always align well with that of differentiation, barcode dropout rates can be too high, library preparation remains laborious, and many studies lack appropriate lineage-specific quality control. We developed scRIPT (<u>s</u>ingle-<u>c</u>ell <u>R</u>apid Integrated <u>P</u>latform for <u>T</u>racing), which combines rapid ribonucleoprotein (RNP)-based barcoding with fluorescent reporter coding sequence insertion, reporter-based enrichment of informative cells by cell sorting, streamlined library preparation, and dual short-read sequencing, coupled to dedicated quality-control metrics for lineage data. scRIPT achieves a low cell dropout rate (<5% of cells presenting without a detectable lineage barcode) and a reliable assignment of barcodes (of >90% of cells). We applied scRIPT to document mouse embryonic stem cell (ESC) differentiation into early hematoendothelial cells and revealed clonal relationships and fate coupling among the emerging cell (sub)types. A subset of cell clones was biased towards single fates, and bifurcating progenitors already express mRNA of canonical regulators of their eventual fate. The results also suggest a heritable, fate-independent clonal memory enriched with imprinted genes. Taken together, scRIPT provides a convenient, rapid, reliable, high-coverage, and quality-controlled platform that combines cell lineage and transcriptome readouts.

## Introduction

Understanding how early embryonic cells are allocated to the germ layers, how cell fate is subsequently determined, and how they acquire lineage-specific functions and further differentiate into tissue-specific or organ-resident cells is essential to understanding normal development and congenital and adult disease. Live-imaging microscopy (more recently often in combination with genetic labeling approaches) has helped track single cells here (Sulston et al., 1983); however, it yields limited-depth, poorly quantified precise lineage relationships. GESTALT (McKenna et al., 2016), scGESTALT (Raj et al., 2018), and ScarTrace (Alemany et al., 2018) methods use highly diverse barcodes in one go while still allowing cell lineage and (sub)type mapping, without limiting the number of traced cells. These single-cell lineage tracing (scLT) methods rely on heritable CRISPR-Cas9-based random edits (cumulative insertions and deletions, also called scars) to the barcodes; these scars can be recorded, thereby mapping evolving cell branches and trees. A key innovation in such lineage tracing transgenic methods was the CRISPR-Cas9 Target Array (TA) (Jinek et al., 2012) to target multiple genes simultaneously (i.e., “multiplex”), thus avoiding the design and testing of high numbers of single guides. Lineage information is then captured from the cell’s transcriptome, yielding both cell (sub)type identity (via the transcriptome) and lineage barcode (as scars).

Many other scLT methods, including MEMOIR, LINNAEUS, Cas9-L1 Barcode, scGESTALT.2, CARLIN, DARLIN, and Spatio-DARLIN (Bowling et al., 2020; Frieda et al., 2017; Gao et al., 2026; Hwang et al., 2019; L. Li et al., 2023; Raj et al., 2020; Spanjaard et al., 2018) have meanwhile been developed and adapted for various experimental setups, thereby markedly improving cell lineage and differentiation studies. However, several challenges remain. For example, barcoding can be too slow and inefficient because it takes time to obtain and maintain sufficient active (i.e., scar-creating) Cas9. Further, preparing lineage libraries remains complicated and costly because it sits at the nexus of unpredictable genome editing and the technical limits of single-cell sequencing. In addition, the field itself and many scLT approaches lack widely accepted quality-control (QC) metrics for lineage data in many of the studies. A scLT variant that addresses these hurdles would noticeably improve various applications, including in cell differentiation, drug response and resistance, tissue repair, immune cell dynamics, and clonal oncogenesis.

Instructed differentiation of cultured mouse embryonic stem cells (mESCs), e.g., into early hematoendothelial lineages (eHELs), is a valuable model for studying the genetic and molecular regulation of cell fate. This model mimics *in vivo* developmental processes; cell differentiation is reproducible, and its duration is short (Ng et al., 2025; Theeuwes et al., 2025). It enables the investigation of the progressive formation of primitive erythrocytes, megakaryocyte progenitors, endothelial cells, and other cell types. Here, scLT would reveal lineage relationships at three hierarchical levels: (i) between cell types; (ii) within a single cell type (including in multi-or bipotent progenitors); and (iii) between clones, through the ability of these to inherit specific traits, behaviors, or developmental bias from a common ancestor cell across multiple cell mitoses (often referred to as clonal memory).

We wanted to apply scLT to the differentiation into eHELs (see also our studies (Cetin et al., 2025; Picco et al., 2026)) and considered several scLT methods available at the start. Our first scLT experiments, however, prompted us to design scRIPT as a new scLT method. It generated about 5,000 unique barcodes with 40,000 starting cells, with cell dropout (the fraction of cells without detectable lineage barcode) below 5%. Importantly, we added QC metrics for lineage data to help remove low-quality cells and problematic barcodes. Also, doublets could be removed using “scar doublet scores”, a useful lineage barcode-based doublet detection score.

The development and application of scRIPT fit into a six-step workflow that (1) defines the developmental trajectory of eHELs; (2) explores, for example, CARLIN (Bowling et al., 2020) briefly, but (3) quickly made us decide to develop scRIPT; and (4) apply scRIPT across eHEL differentiation and annotate the cell (sub)types; (5) merge lineage and transcriptome data with new QC metrics; and (6) answer several questions that emerged during this work. Questions would be whether the different cell types’ developmental trajectories are coupled to each other, whether the cells are biased within the progenitor state concerning their differentiated cell type, and whether clones with different origins share a transcriptomic signature that belongs to their origin and can still be detected even in the differentiated state(s). By applying scRIPT in this cell system, fate couplings between cell (sub)types were mapped, and the expected within-trajectory relationships alongside cross-compartment couplings were recovered. We also found that a subset of clones is biased towards single fates, and that bifurcating progenitor cells already express mRNAs of canonical regulatory genes of their eventual fate.

## Results

### Defining the developmental trajectory of eHELs by developing scRIPT first

Following on from our previous work that used ESC-derived eHELs (Cetin et al., 2025; Picco et al., 2026) we asked (1) what are the lineage relationships of eHELs, and which differentiating cell types share a clonal origin?; (2) do individual ESC clones and derived progenitor cells show non-random fate bias, and would such biased progenitors already express mRNA of canonical regulators of the eventual cell fate?; and (3) do clones carry a heritable, fate-independent transcriptional memory that marks a clone irrespective of which cell type(s) its cells adopt? (Fig. 1a, Q1-Q3 panels). One of our initial approaches employed CARLIN and its cell line during a 4-day differentiation of mESCs into eHELs, followed by library preparation using the CARLIN protocol (Bowling et al., 2020). Regrettably, and at least in our hands, this resulted in relatively low lineage-barcode capture rates (Fig. S1; Table S1), presumably for the reasons outlined and effectively resolved by these authors. Building upon their recommendations (Bowling et al., 2020) and leveraging our expertise in ESC→eHEL differentiation, we designed scRIPT. In an effort to enhance lineage barcode capture efficiency, we introduced major adaptations to the scLT methodologies noted above. They are fivefold (see Table 1): using an appropriate scRIPT-mESC cell line; rapid and efficient lineage barcoding by ribonucleoprotein (RNP)-based transfection; pre-selection of cells with capturable barcodes; simplified lineage library preparation; and cost-efficient, dual short-read sequencing, respectively.

**Fig. 1.**
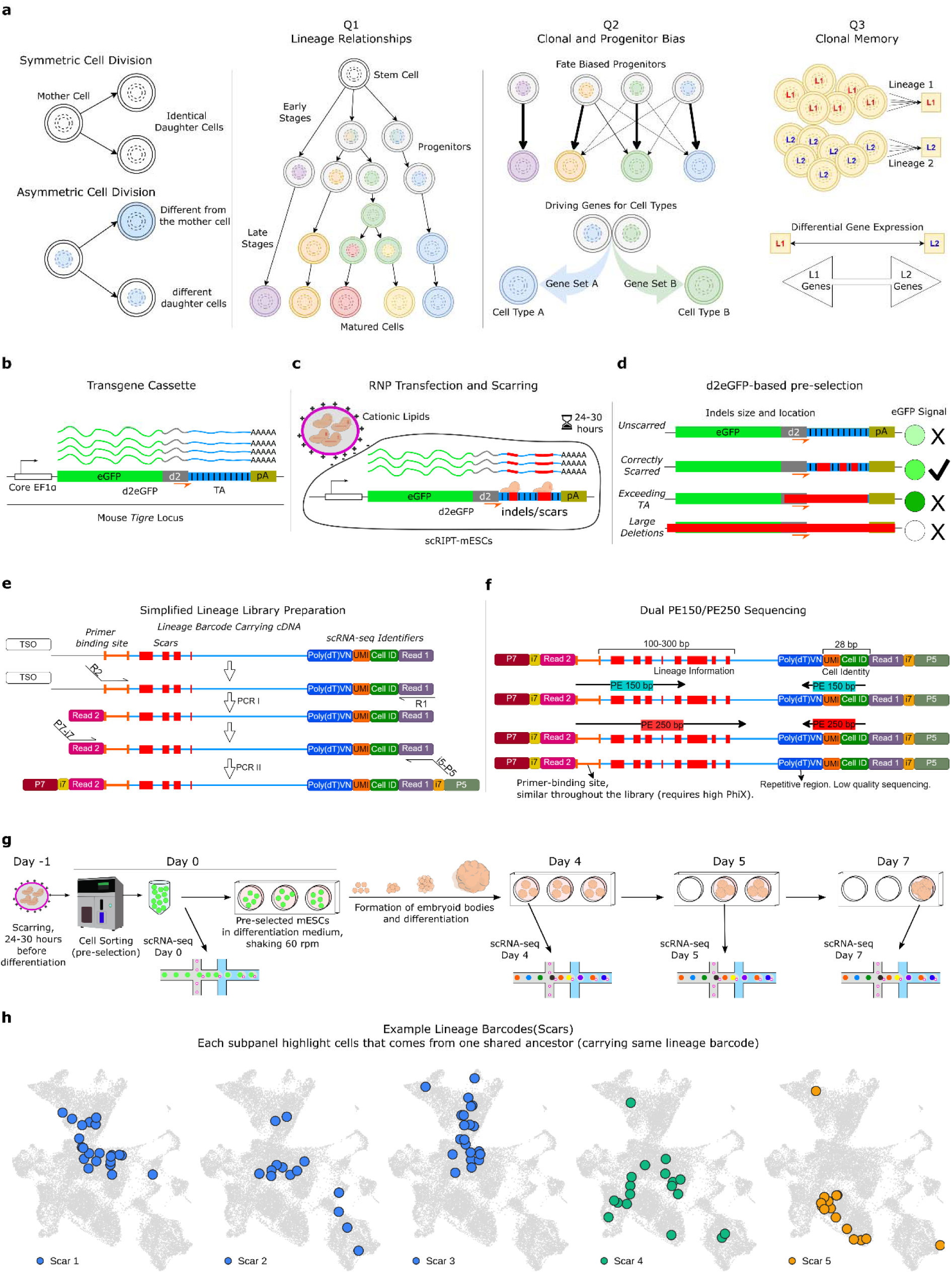
Overview of the scRIPT platform and workflow. The design rationale and implementation for each optimization step are given in Table 1. **a**, (left to right) Symmetric versus asymmetric cell division enables the generation of identical and different daughter cells, respectively, and co-determines hierarchical lineage relationships. Question (Q)-1 (see main text) is graphically represented by a hypothetical lineage tree in which various, identifiable early or later progenitors give rise to one or more distinct (including mature) cell types (with the colored circles representing identifiable, transcriptionally distinct states). Q2: By integrating lineage barcodes with single-cell transcriptomes (see main text), early progenitor populations and their gene expression profiles can be identified, and their possible bias investigated, before (histo)morphologically detectable cell differentiation. The comparisons reveal differential gene expression (shown here as Gene Set A vs B) that drives commitment toward specific cell types (here A and B, respectively). Q3: Assessment of the existence of clonal memory by comparing cells of the same differentiated type but with distinct lineage trajectories (here Lineage L1 vs L2), which reveal persistent signatures inherited from progenitor states. **b**, The scRIPT-mESC line, generated by stable integration of a lineage-recording cassette (*EF1*α core promoter, d2eGFP reporter, TA, and poly(A) signal sequences) at the *Tigre* safe-harbor locus. **c**, Barcoding by high-efficiency transfection of Cas9-RNP, using cationic lipids, achieving scarring within 24 hours. **d**, Possible outcomes of FACS enrichment for cells with usable barcodes, defined as indels that retain intact primer-binding sites while showing changes (from white to darker green) in the level of the d2eGFP signal. **e**, Streamlined, simplified library preparation using codon-optimized primer-binding sites, enabling a two-step PCR, but still compatible with 10X Genomics Chromium Single Cell 3′ v3 chemistry. (P5/P7: flow-cell binding sequences, i5/i7: sample index barcodes, Read1/Read2: paired-end sequencing reads, UMI: Unique Molecular Identifier, TSO: template switch oligo) **f**, Dual sequencing combining PE150 sequencing for routine analysis with longer PE250 sequencing for comprehensive barcode recovery (for abbreviations, see panel e). **g**, The scRIPT experiment: scarring was performed 24-30 hours before the start of differentiation and, after pre-selection for increased d2eGFP at 30 h post-transfection, a cell portion was taken for D0 scRNA-seq, and the remainder of the cells was transferred to differentiation medium (Cetin et al., 2025), with scRNA-seq applied at D4, D5, and D7. Because each time point derives from independent mESC-derived embryoid bodies (indicated in the panel between D0 and D4/5/7), the lineage analysis is restricted to within each time point, and barcodes are not traced across time points. **h**, Example of scarred lineage barcodes on the differentiation map: a UMAP with cells sharing a representative barcode (scar, numbered 1 to 5) highlighted, with one barcode per panel; all other cells are in light grey.

**Table 1.**
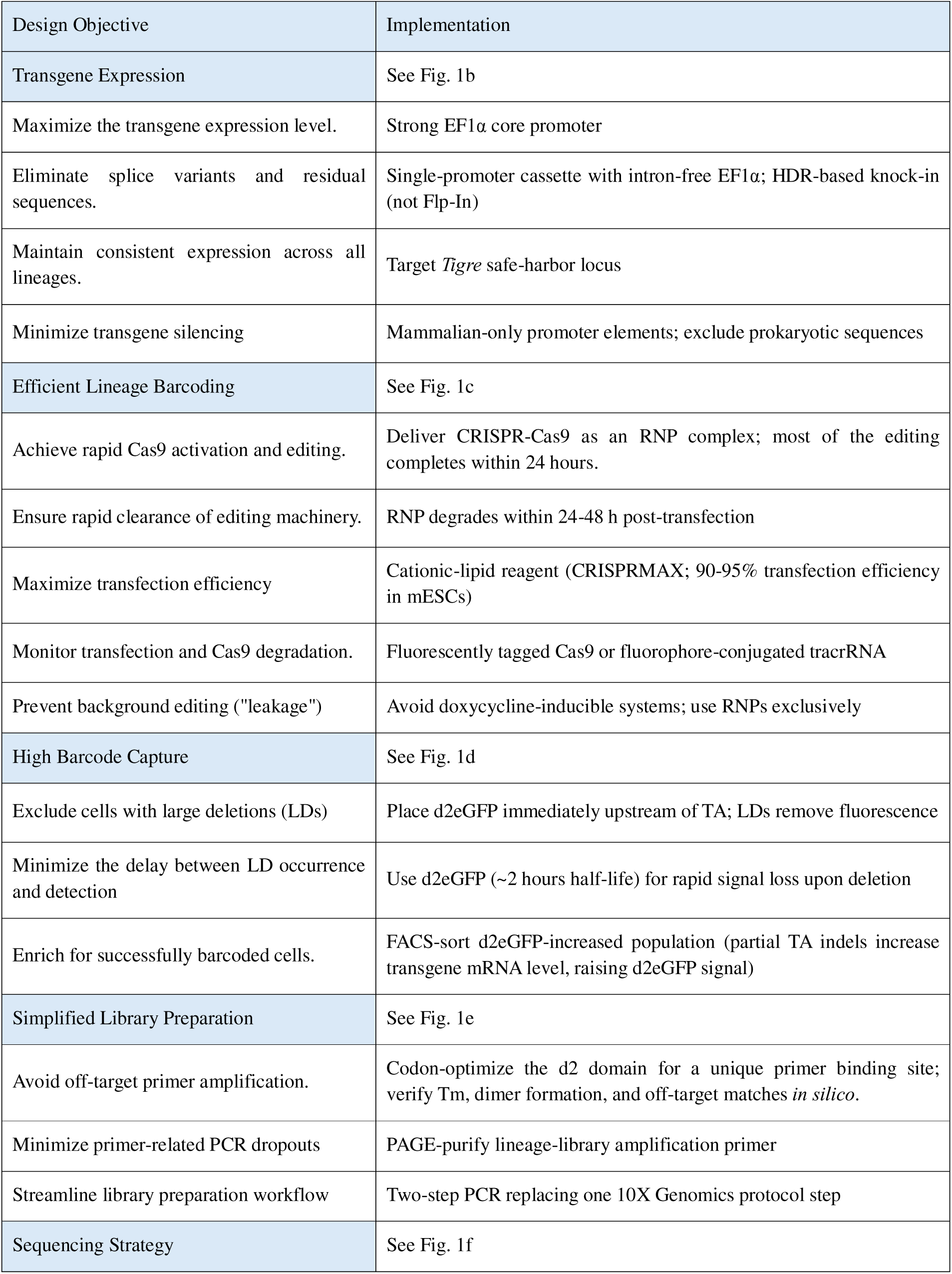
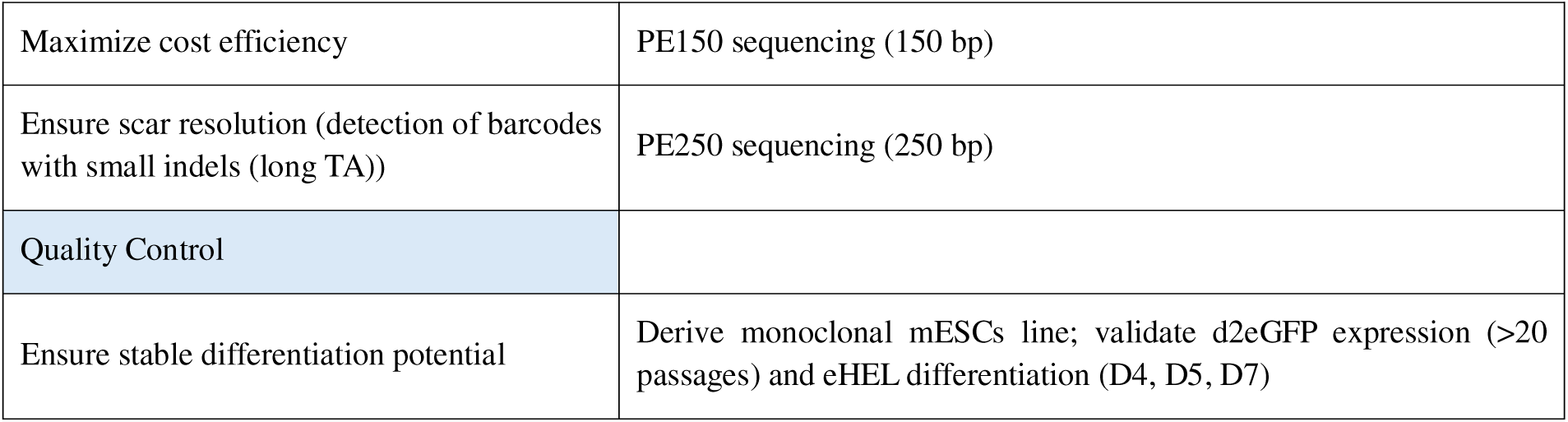
Design objectives and implementations for scRIPT’s wet-lab optimization.

| Design Objective | Implementation |
| --- | --- |
| Transgene Expression | See Fig. 1b |
| Maximize the transgene expression level. | Strong EF1α core promoter |
| Eliminate splice variants and residual sequences. | Single-promoter cassette with intron-free EF1α; HDR-based knock-in (not Flp-In) |
| Maintain consistent expression across all lineages. | Target *Tigre* safe-harbor locus |
| Minimize transgene silencing | Mammalian-only promoter elements; exclude prokaryotic sequences |
| Efficient Lineage Barcoding | See Fig. 1c |
| Achieve rapid Cas9 activation and editing. | Deliver CRISPR-Cas9 as an RNP complex; most of the editing completes within 24 hours. |
| Ensure rapid clearance of editing machinery. | RNP degrades within 24-48 h post-transfection |
| Maximize transfection efficiency | Cationic-lipid reagent (CRISPRMAX; 90-95% transfection efficiency in mESCs) |
| Monitor transfection and Cas9 degradation. | Fluorescently tagged Cas9 or fluorophore-conjugated tracrRNA |
| Prevent background editing ("leakage") | Avoid doxycycline-inducible systems; use RNPs exclusively |
| High Barcode Capture | See Fig. 1d |
| Exclude cells with large deletions (LDs) | Place d2eGFP immediately upstream of TA; LDs remove fluorescence |
| Minimize the delay between LD occurrence and detection | Use d2eGFP (~2 hours half-life) for rapid signal loss upon deletion |
| Enrich for successfully barcoded cells. | FACS-sort d2eGFP-increased population (partial TA indels increase transgene mRNA level, raising d2eGFP signal) |
| Simplified Library Preparation | See Fig. 1e |
| Avoid off-target primer amplification. | Codon-optimize the d2 domain for a unique primer binding site; verify T<sub>m</sub>, dimer formation, and off-target matches *in silico*. |
| Minimize primer-related PCR dropouts | PAGE-purify lineage-library amplification primer |
| Streamline library preparation workflow | Two-step PCR replacing one 10X Genomics protocol step |
| Sequencing Strategy | See Fig. 1f |
| Maximize cost efficiency | PE150 sequencing (150 bp) |
| Ensure scar resolution (detection of barcodes with small indels (long TA)) | PE250 sequencing (250 bp) |
| Quality Control |  |
| Ensure stable differentiation potential | Derive monoclonal mESCs line; validate d2eGFP expression (>20 passages) and eHEL differentiation (D4, D5, D7) |

Our scRIPT-mESC line carries a (single-copy) transgene cassette integrated at the safe harbor *Tigre* locus on one of its two mouse chr9, and in which a CARLIN TA and destabilized (d) d2eGFP (X. Li et al., 1998; Warren et al., 2010) reporter share a single transcript (see Methods; Fig. 1b; Table S2). Barcoding is by transient delivery of CRISPR-Cas9 RNP by cationic-lipid transfection, which is rapid and efficient (editing up to 95% of the cells) and thus avoids the longer time windows of other (including inducible system) alternatives (Fig. 1c). Large indels (that delete the TA) are the major source of dropouts (e.g., P7 in Fig. S2b-d). d2eGFP-based pre-selection (P5, population with increased eGFP signal) outcomes are summarized in Fig. 1b-d. scRIPT thus pre-selects cells retaining a capturable barcode through the use of short-lived d2eGFP (≈2 h half-life) as a TA-deletion sensor. Cells that lose the TA due to a large deletion also lose d2eGFP-coding parts and can be sorted away (by FACS) at 24-30 h post-transfection (P7 in Fig. S2b-d). Cells with informative partial indels that increase transgene mRNA abundance and show increased d2eGFP levels can be collected by FACS (P5). Based on the eGFP intensities, we assigned signal groups (P4-P6) at 30 h post-transfection (Fig. S2; see also Fig. 1d). We observed that the number of mRNA molecules originating from arrays carrying the d2eGFP transgene and belonging to cells with scars, which result in a shorter 3′UTR of the transgene, is significantly higher in cells that carry the WT/unscarred transgene (Fig. S3).

For subsequently generating the lineage library, scRIPT replaces one step of the standard 10X Genomics 3′ scRNA-seq pipeline (see Methods) with a simple two-step PCR (PCR I and PCR II, Fig. 1e). For the latter, it uses a gel-purified primer designed for a codon-adjusted (for more information, see Methods) primer binding site next to the TA to minimize bias and dropout (see Methods; Fig. 1e; Fig. S4). Then, the libraries are sequenced using a dual short-read strategy; it involves an initial, poolable 150 bp paired-end (PE150) run that gauges barcode coverage and then guides a deeper, targeted 250 bp paired-end (PE250) run (Fig. 1f). Together, these maximize recovery while reasonably limiting the cost and turnaround time (Fig. 1f; Figs. S5, S6).

### Applying scRIPT across ESC→eHELs differentiation and annotating cell types

After lineage barcoding of the TA-containing undifferentiated mESCs, we enriched the d2eGFP-increased population by FACS (see above). An aliquot was taken immediately after FACS to generate the day-0 (D0) scLT sample, and the remaining cells were subjected to differentiation in cell culture (as applied previously, (Cetin et al., 2025)). Differentiated cells were then collected on D4, D5, and D7, respectively. Lineage-library preparation and sequencing were carried out as described (see Methods) (Fig. 1g,h; Table S3).

scRNA-seq (see Methods) analysis showed that the mESCs gave rise to eHELs, as well as other mesodermal derivatives and endodermal and ectodermal cells, as found previously (Cetin et al., 2025; Picco et al., 2026). scLT was performed at the four time points by selecting different embryoid bodies (EBs) from the cell pool, yielding D0, D4, D5 (divided into three libraries), and D7 (three libraries) samples. Each time point allows us to create lineage “relationships within” since cells form EBs and multiply within these, while different EBs originate from different initial cells. After the QC of each sample’s transcriptome (see Methods), we retained 35,891 cells.

Next, the scRNA-seq datasets were used for annotating cell clusters based on known marker genes (Fig. 2a-c; Figs. S7, S8). The intended eHELs comprised 5,543 cells, representing 21.7% of the 25,562 D4-D7 cells. These eHELs included hematoendothelial progenitors (HEPs), but also erythro-myeloid progenitors (EMPs), primitive erythrocytes, megakaryocyte progenitors, endothelium, and endocardium (Fig. 2b), as documented previously (Cetin et al., 2025; Picco et al., 2026). Other mesodermal cells were also annotated (Fig. 2b), consistent with our prior work (Cetin et al., 2025; Picco et al., 2026). These are cardiopharyngeal mesoderm (CardPhPr), (pre)somitic tissue cells (SomiticTis), immature cardiomyocytes (CardioM), epicardium, and mesenchyme/mesothelium (MchyMthel), respectively. Bridging these (see Fig. 2a) were early mesodermal populations, defined as nascent mesoderm (EarlyMesod) and mixed mesoderm (MixMesod) progenitor cells. The other clusters correspond to remaining naïve pluripotent mESCs (Naïve_PSCs), and epiblast-like cells (indicated here briefly (see also (Imaz-Rosshandler et al., 2023)), as Epiblast), mesendodermal cells (endodermal and notochord/node-like), and ectodermal cells, whose transcriptome resembled non-neural cells of the floor plate (of the hindbrain) and dorsal cerebral neurons (Fig. 2a,b,d; Fig. S9).

**Fig. 2.**
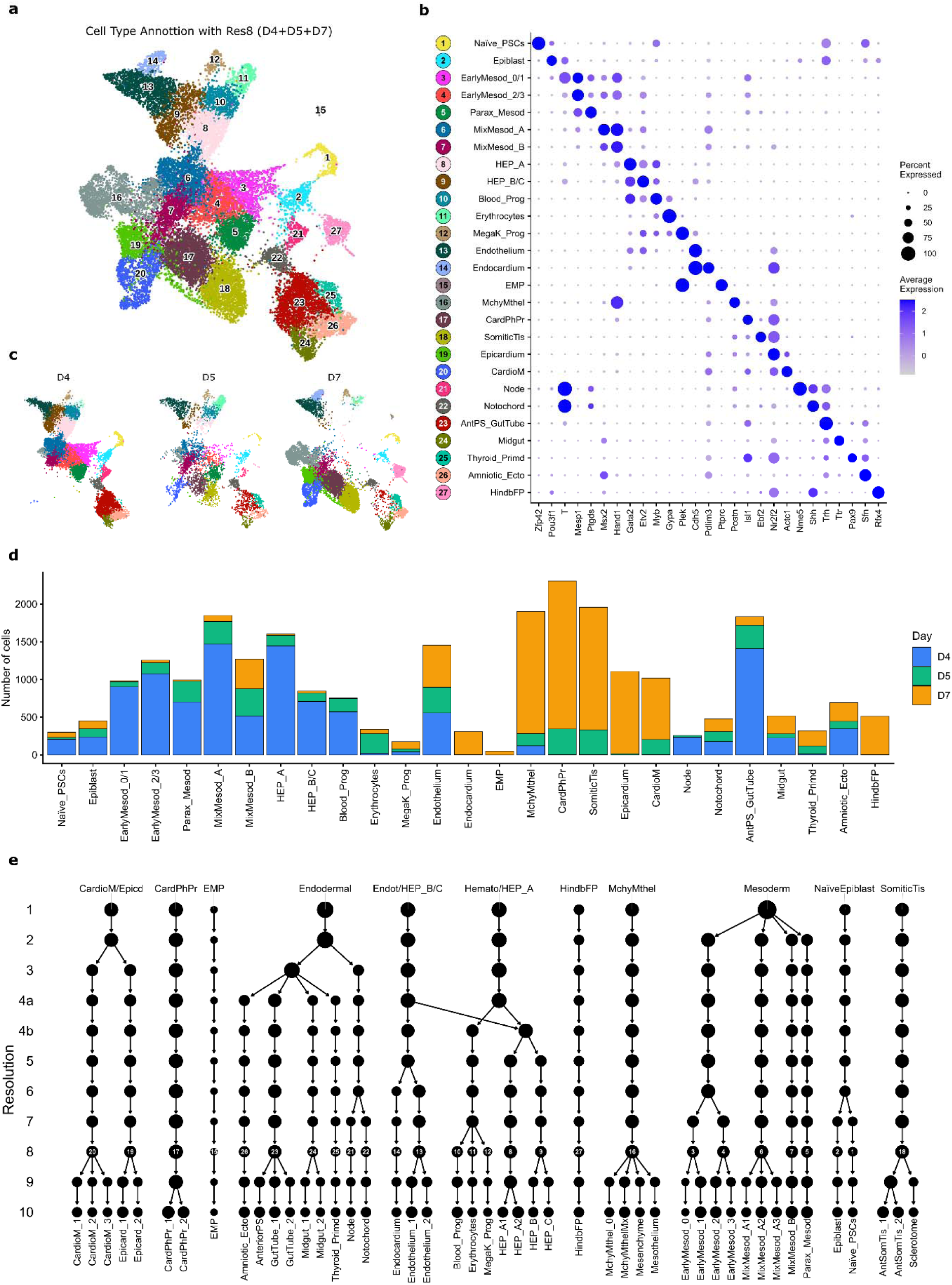
Harmony-integrated single-cell transcriptomes resolve marker-annotated cell types across early differentiation (D4-D7) of mESCs; nested annotations of increasing resolution define the cell-type schemes used for all downstream lineage-tracing analyses. **a**, Uniform Manifold Approximation and Projection (UMAP) embedding of 25,562 Harmony-integrated cells from D4/D5/D7 (11,016; 4,123; and 10,423 cells, respectively), colored by cell-type identity. Each cell type is labelled with a number; the number-to-identity key is shared with the y-axis of b. D4 was prepared as a single sequencing library; D5 and D7 were prepared as three libraries each (see main text and Methods; one sample split for capacity, not for serving as biological replicates). **b,** Dot plot of canonical marker genes (x-axis) across cell types (y-axis), used for cell type assignment. Dot size, percentage of cells in each cell type expressing the gene; color, mean expression z-scored across cell types per gene. Colored numbered badges on the y-axis match the cell types and colors in panel a. **c,** The embedding in panel a, split by differentiation time point, colored by cell type (as in panel a). **d,** Stacked bar plot of cell number per cell type (x-axis as in panels a and b), colored by differentiation time point (D4, blue; D5, green; D7, orange, and unlike in panels a, c, and e, where color denotes cell type). **e,** Cluster tree (clustree) showing how cells are partitioned under the eleven curated annotations of increasing resolution (res1-10, with alternative splits 4a and 4b, at level 4). Rows are annotation levels, from coarsest (res1, top) to finest (res10, bottom). Each node is one cell type at that level, with node size proportional to the number of cells it contains. Edges link clusters that share cells between adjacent levels. Numbered badges on the res8 row identify cell types using the same numbering as in a. Cell type names are shown for the top (res1) and bottom (res10) rows.

We annotated cell (sub)types at several hierarchical resolutions, from coarse to fine, because differentiation proceeds continuously. We realize that most subsequent analyses based on this annotation remain sensitive to the clustering resolution, batch effects, and possibly misannotations. To minimize these problems, we carried out multiple rounds of clustering and annotation (Fig. 2e; Fig S7-S9; Table S4). We can select different parameters, particularly for groups of transitioning cells where “cell type” and “cell state” are blurred; therefore, results that depend on such complex cell-type and cell-state assignments should always be interpreted carefully in these types of datasets, regardless of the methodology used.

### Merging lineage and transcriptome data with new QC metrics

After sequencing the lineage libraries, we aligned reads to the reference TA, assigned lineage barcodes to single-cell barcodes using the cell barcodes detected in the scRNA-seq, and merged the lineage and transcriptome data after removing overrepresented scars (see Methods). To identify doublets from lineage data alone, we devised a barcode-based scar-doublet score, defined as “more than one lineage barcode in a single cell” (see Methods). It agrees with the transcription-simulation-based scDblFinder (Germain et al., 2021) for 93% of cells; it also flagged a further 1% that scDblFinder would have missed, while scDblFinder called an additional 6% of cells that the scar-doublet score would not support (Fig. 3a). Cells were then filtered for barcode quality, removing those with short or unresolvable fragments, scar-doublets, missing barcodes (called 0NA), and unedited wild-type barcodes (1WT).

**Fig. 3.**
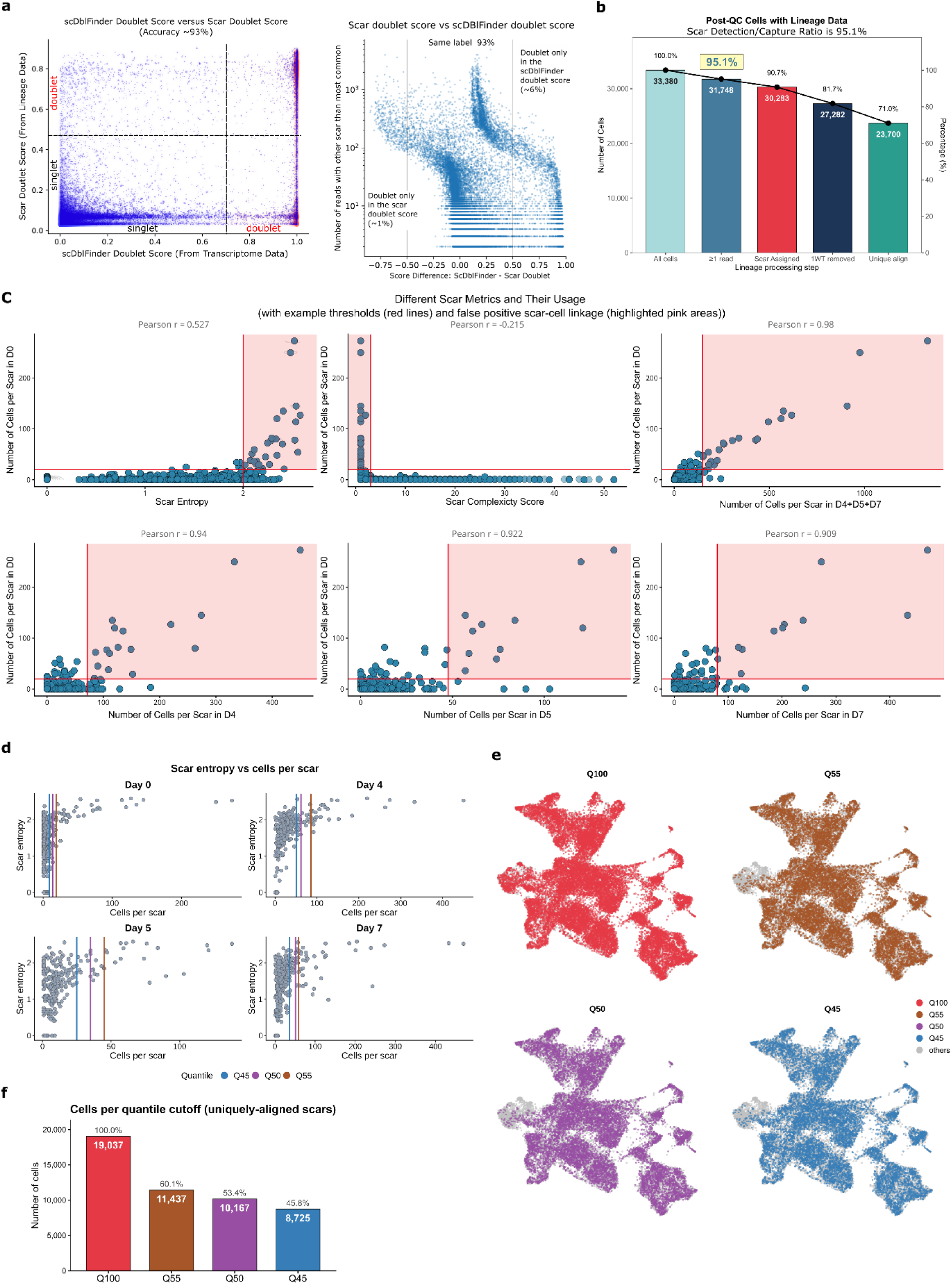
Cross-validated doublet removal and scar-level QC yield high-confidence lineage assignments; cells-per-scar quantiles define the Q55/Q50/Q45 stringencies. **a**, Lineage (scar doublet score) vs transcriptome-based (scDblFinder) doublet scores. Left panel, scDblFinder (x-axis) vs scar doublet (y-axis). Right panel, score difference (scDblFinder - scar doublet, x-axis) vs lineage reads assigned to a scar other than the cell’s most abundant (y-axis, in log scale): concordant (93%), scDblFinder-only doublets (6%), and scar-only doublets (1%). **b,** Cell numbers through successive lineage-data QC steps, pooled over D0-D7 (y-axis on the left, cell number; right, % of all cells). All cells, total in transcriptome data; ≥1 read, ≥1 read in lineage library; scar assigned, enough reads for confident assignment; 1WT removed, single unedited (1WT), with TA excluded; unique alignment, reads aligning to a single full-length TA (retained downstream). **c**, Scar-level metrics flagging over-represented, uninformative scars, using D0 (earliest, pre-differentiation time point, before clonal expansion inflates counts) as reference. Each dot is one scar; D0 cells per scar (y-axis in all graphs) vs scar entropy (x-axis in top left panel; Pearson r = 0.527). Same for scar complexity score (see Methods; top middle; r = −0.215), total cells per scar over all time points (top right panel; r = 0.98), and cells per scar at D4 (bottom left panel; r = 0.94), D5 (bottom middle panel; r = 0.922), and D7 (bottom right panel; r = 0.909). Higher entropy and lower complexity tend to accompany a larger D0 clone size, which tracks later and overall clone size. **d,** Per-day of differentiation quantile thresholds. Scar entropy (y-axis) vs cells per scar (x-axis) for D0, D4, D5, D7; the three vertical lines per graph mark the Q55, Q50, Q45 cutoffs (colors in key). **e,** UMAP of D4-D7 cells (as in Fig. 2), highlighting cells retained at each stringency, from least to most stringent (Q100, Q55, Q50, Q45); non-retained cells are grey. Colors as in panels d, f. **f,** Cells retained at each cutoff among uniquely aligned-scar cells passing full transcriptome QC (i.e., their % vs Q100). Cutoffs were computed before doublet removal and further filtering, so realized fractions exceed nominal quantiles. Colors as in panels d, e.

scRIPT achieved high capture rates (95.1% of cells with a scar/lineage transcript) and assignment (90.7%) (cells with a confidently assigned scar/lineage barcode). After filtering, 23,700 cells (out of 33,380; 71.0%) were analyzed. Relaxing the filters (yielding a higher risk of including barcodes with barcode homoplasy) would increase the usable set to 27,282 cells (81.7% of the cells) or more, but the stricter criteria were applied to maximize the confidence in lineage assignments (Fig. 3b). Some scars recur far more often than expected and can generate false lineage relationships. This is why four complementary QC metrics were introduced (for details, see Methods): a scar’s abundance at D0 (at most 8 cells, after an estimate of three cell divisions; higher counts would indicate independent recurrence (barcode homoplasy) or failure to cell-sort); its cumulative count across samples; ‘scar entropy’ (higher entropy ∼ high scar homoplasy); and ‘scar complexity’ (multi-fragment scars and indels give higher complexity scores), respectively (for operational definitions, see Methods; Fig. 3c, Fig. S10). Applying a quantile cutoff on the cells-per-scar distribution at four stringencies (i.e., Q100 (no cut-off), Q55, Q50, and Q45, named for their upper-quantile cutoff) produced the high-confidence lineage datasets used in all subsequent scLT analyses (Fig. 3d-f; Table S5).

### Clonal coupling maps the lineage relationships in eHEL differentiation

To determine which differentiated cell types share a clonal origin, we used the lineage barcodes recovered by scRIPT to quantify fate coupling, defined as the degree to which two cell types co-occur within the same clones more than expected by chance. For each used differentiation day and annotation resolution, we used the CoSpar package (Wang et al., 2022) to score every cell-type pair by its normalized clonal co-occurrence. This involves permuting clone labels separately for each time point while preserving clone sizes, and doing this for 10,000 permutations (see Methods). We then call a pair coupled when its co-occurrence exceeded the permutation null (p < 0.05) and was supported by ≥3 distinct clones spanning the two types (Fig. 4a-f, Figs. S11-S17, for details, including in the legends; Table S6).

**Fig. 4.**
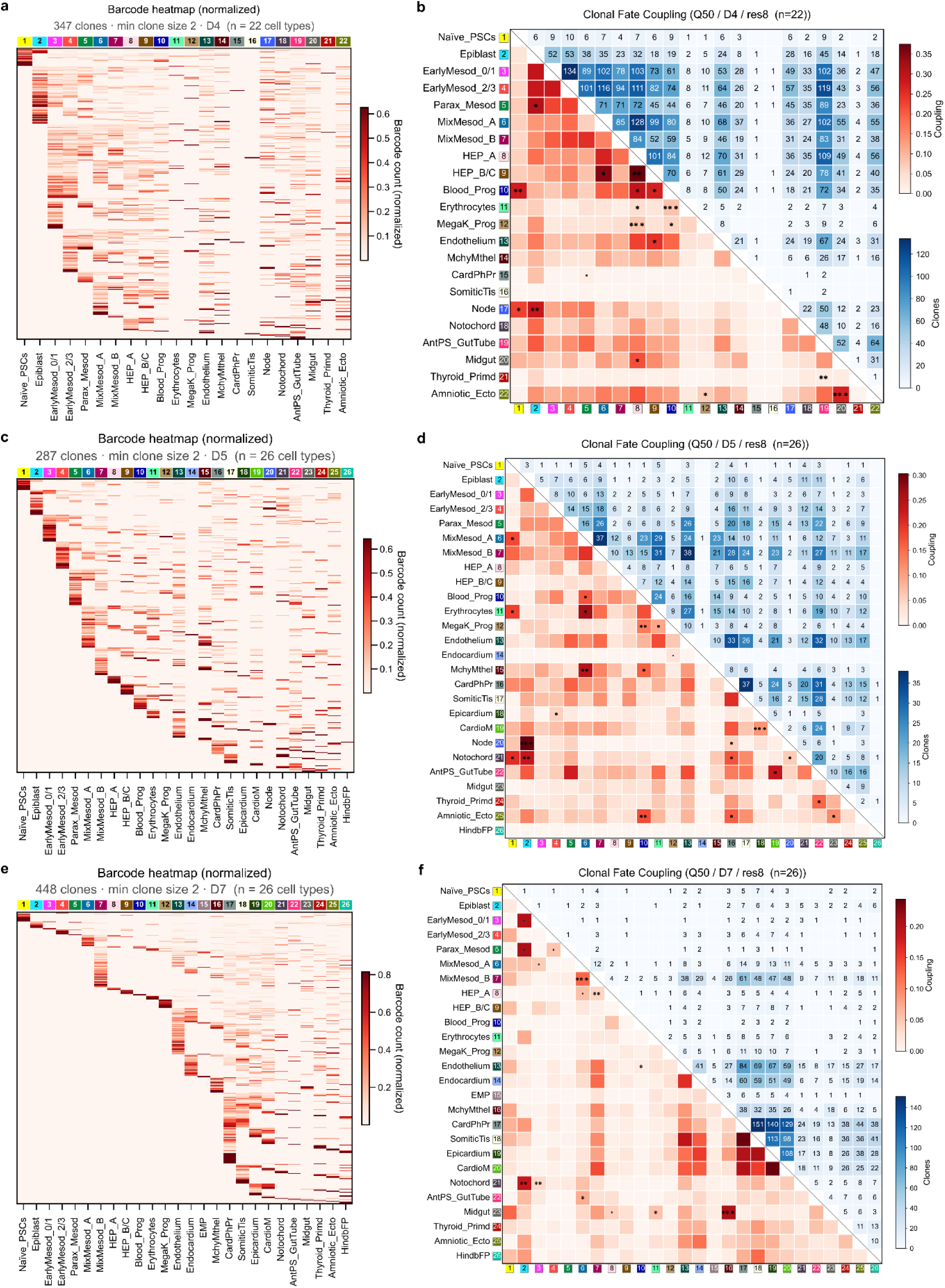
Clone-resolved cell-type composition and pairwise fate-coupling across differentiation time points (at Q50, for res8). **a**, Clonal profile of the normalized proportion of each annotated cell type (column) within each clone (row). Barcode heatmap, D4 (min clone size 2; n=347 clones encompassing 22 annotated res8 cell types). Rows = individual clones; columns = res8 cell types (top color strip = cell-type palette also used in Fig. 2). Cell value = per-clone-normalized barcode count (white → dark red). **b**, Heatmap of clonal coupling scores across cell types at D4 (shown in panel a, for n=22 cell types). Lower-left triangle = coupling score (white → red); upper-right triangle = number of supporting clones per pair (white → blue; with the count printed in each cell). Significance marks in lower-left: ‘*’ / ‘**’ / ‘***’ = permutation p < 0.05 / 0.01 / 0.001 AND ‘clone_support ≥ 3’; ‘.’ = p < 0.05 with ‘0 < clone_support < 3’ (weak support); blank = p ≥ 0.05, or ‘clone_support = 0’. **c**, As panel a, for time point D5 (n=287 clones; 26 res8 cell types). **d**, As panel b, for Q50, D5, res8 (n=26 res8 fates). **e**, As panels a and c, for D7 (n=448 clones; 26 res8 fates). **f**, As panels b and d, for Q50, D7, res8 (n=26 fates).

A key challenge is determining which scar/clone barcodes reflect genuine clonal lineages and which are artifacts of similar repair patterns happening across different individual clones. We addressed this earlier by applying Q100 (every clone, including the largest dominant clone), Q55, Q50, and (the most stringent) Q45 filters on the cell-per-scar distribution. Applying the fate-coupling pipeline across all four stringencies, we compared Q100 with the other three. We observed that Q100 produced false-positive clonal couplings and also added noise, causing the loss of authentic couplings (Fig. 5a). When we compared the combined fate couplings across stringencies, Q100 was significantly different from every other stringency on each studied differentiation day. By contrast, Q50 and Q55 showed no significant differences on any such day, and Q45 showed significant differences in D5 and D7, but not in D4 cells. Relative to Q50 (368 significant fate couplings), the total number of fate couplings was as follows: Q45 −14%, Q55 +3%, Q100 +44% (Fig. 5b). Taken together, these results show that Q100 drastically inflates the fate couplings and is not reliable for further fate-coupling analysis. The other three (Q45, Q50, and Q55) gave more reproducible results, with small differences among them (Fig. 5b). It should be noted that Q45 may be too strict in some, but few, cases, particularly for cell types with too few cells, and this for some clusters at D5 and D7 (*data not shown*). The couplings described below will draw on these three stringencies (i.e., without Q100), the multi-resolution annotation, and datasets from the different days of differentiation.

**Fig. 5.**
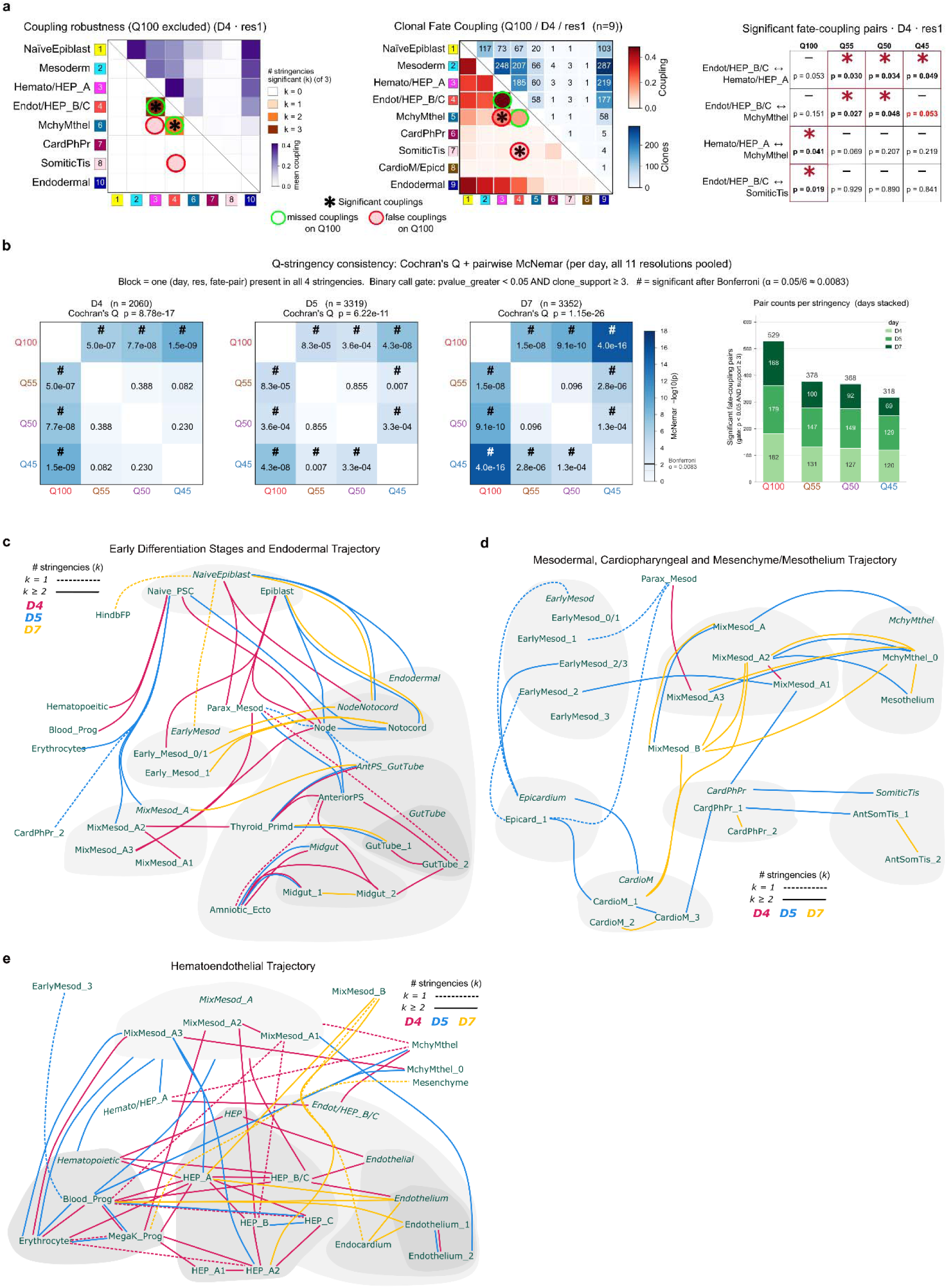
Q100 inflates the lineage-coupling network, whereas Q45 / Q50 / Q55 retain a stable core, spanning three early-differentiation trajectories. **a**, Coupling robustness at D4 · res1. Left panel, robustness heatmap (Q100 excluded): lower-left = # of Q45 / Q50 / Q55 significant (white to dark orange, 0 → 3); upper-right = mean coupling (white to darker purple). Middle panel, Q100-only heatmap: coupling (white to darker red); clone count (white to darker blue). Overlay: green=pairs significant at all of Q45 / Q50 / Q55 but missed at Q100 (false negatives); red = pairs significant only at Q100 (false positives). Right panel, pair-level table for the 4 fate-pairs significant at ≥ 1 stringency: rows = pairs, columns = stringencies; cells show coupling, p-value, and ‘clone_support’. * + red box = passes ‘p_greater < 0.05 AND clone_support ≥ 3’; not-significant = n.s. **b**, Stringency-consistency tests pooled across all (day × resolution × fate-pair) combinations testable at all 4 stringencies (same gate as panel a). Three 4 × 4 heatmaps (per day) show pairwise exact McNemar p-values; Cochran’s Q omnibus p-value is given above each; asterisks = Bonferroni-significant (α = 8.33 × 10 ³). Right: Q100 calls 40-65 % more pairs than the project-grade stringencies. McNemar localizes disagreement to Q100 vs each of Q45/Q50/Q55. **c**, Scheme of robust lineage couplings. Edges = pairs significant at ≥ 1 of Q45 / Q50 / Q55, pooled across all 11 resolutions (res1-res10). Edge color = day (D4 / D5 / D7; palette as in panel b). Edge style: solid = k > 1 (≥ 2 scar stringencies (Q55, Q50 and Q45)); dashed = k = 1. **d**, As in panel c, now for the cardiopharyngeal / somitic/mesenchymal trajectory (CardPhPr, CardioM/Epicd, SomiticTis, MchyMthel + coarser-resolution parents). **e**, As in panel c, for the hematoendothelial trajectory (HEP_A / B / C, Hematopoietic, Endothelial, Blood progenitors + coarser-resolution parents).

A challenge in scRNA-seq studies of continuously differentiating systems like ours is deciding where to draw the boundaries between cell clusters (see Fig. 2). Rather than committing to a single resolution and treating the outcome as ground truth, we generated cell-type annotations (11 in total) at increasing resolution, from res1 (which are broad cell types, low resolution) to res10 (fine cell types, high resolution), with res4 notably provided as two variants (res4a, res4b). Collectively, this allows selection of a resolution that meets a particular biological question. Higher resolutions accordingly yielded higher significant fate couplings, allowing relationships between subclusters to be resolved. Importantly, many couplings fell within single trajectories and recapitulated known developmental relationships of cells. Within the hematoendothelial trajectory, for example, the eHELs were coupled with HEPs and with each other (Figs. 4, 5). The strongest couplings were observed among the more differentiated blood and endothelial subtypes (Figs. 4, 5). Comparable within-trajectory couplings were recovered from the mesenchyme/mesothelium, cardiopharyngeal, endodermal, and naïve pluripotency/epiblast-like trajectories. Recovering these expected, albeit complex relationships from lineage information alone indicates that scRIPT can detect true clonal signals (Fig. 5c-e; Fig. S18).

Importantly, beyond these expected relationships, we observed several couplings that span trajectories. Most notably, eHELs and their subclusters were coupled with the mesenchyme/mesothelium trajectory and its subclusters (Fig. 5e). This is consistent with (Biben et al., 2023): their *in vivo* barcoding of mouse yolk sac cells identified three dual-outcome precursors linked within the hemangioblast (blood and endothelium), mesenchyme-angioblast (mesenchyme and endothelium), and hematomesoblast (blood and mesenchyme). We further observed couplings between endothelial cell subclusters and subclusters of cardiomyocytes-1 and somitic tissues, and cross-germ-layer couplings between endodermal subclusters and both HEP and hematopoietic subclusters.

These couplings should be interpreted with caution, particularly those involving cross-germ-layer coupling. For example, scars are introduced before the onset of cell differentiation, when Naïve_PSCs and Epiblast-like cells are still proliferating, and their daughter cells can separately give rise to presumptive primitive endodermal and presumptive mesodermal cells. Because barcoding precedes germ-layer specification, any subsequently diverging lineages can co-occur within a clone by virtue of their shared early ancestry. This possibility may not be unique to cross-germ-layer pairs. The filters available in this setting (the coupling score, the permutation test, and the requirement for ≥ 3 distinct clones spanning a pair) do not remove this contribution, and no other established method currently removes it either. Additionally, permuting the clone labels separately for each D4, D5, and D7 time point preserves the clone sizes but not the correlated multi-lineage output of individual large early clones, which can raise specific couplings above the null. Consequently, statistical significance reflects a genuine clonal connection but, on its own, not a specific lineage relationship. We accordingly interpret the couplings also on biological grounds, i.e., those matching already acknowledged developmental relationships or known, independent lineage data (e.g., as reported by (Biben et al., 2023)). These aforementioned cross-germ-layer couplings, but without developmental precedent, are in any case interpreted conservatively as shared early ancestry rather than specific (later) progenitor. For targeted lineage relationships, we recommend trying to induce lineage barcoding at a later stage (e.g., comparable to during gastrulation or post-gastrulation, if efficient enough) to narrow the shared-ancestry window.

During differentiation, the respective cell types emerge at distinct time points, make up varying fractions of the differentiation pool, and likely persist for different durations. So, not every fate coupling will be detectable on every time point or day, while some will appear over multiple days. For focused questions about a specific lineage relationship, we recommend adapting the sampling window by collecting samples at multiple, closely spaced time points and combining this with the planned, timed introduction of lineage barcodes -if feasible - as mentioned above. Even so, scRIPT recovered the couplings expected from established hematoendothelial differentiation, confirming that the method reports genuine clonal relationships.

### Clonal fate bias and transcriptional priming

#### The lineage barcodes reveal non-random fate bias in individual clones

Fate coupling describes the relationships between cell types. We asked whether individual clones are, as it were, primed, thus contributing their cells disproportionately to one fate. For each clone (those with ≥2 cells), we tested enrichment per cell type against a hypergeometric null (here, the observed overlap occurs by chance, not by association) that accounts for clone size and fate abundance. This serves as a control for the false-discovery rate within clone-size strata (CoSpar, (Wang et al., 2022); see Methods). A substantial minority of clones were indeed significantly biased toward a single fate (Fig. 6a). This amounts to 16-32% across the applied days, at the three intermediate annotations, increasing only modestly with finer annotation. However, the chance of calling a clone primed rises steeply with clone size, from 4-12% for the smallest clones to 64-79% for the largest. This may resemble a “community effect” (the need to provide enough target cells prone to differentiation cues), proposed to influence the mesoderm induction response (i.e., muscle cell formation) in amphibian embryo animal cap explants (Standley et al., 2002; Standley & Gurdon, 2002). So, many unprimed clones may be too cell-poor to reach significance, while primed clones may be skewed towards large clones (median of about 15 cells). Priming may therefore be under-detected altogether (Fig. 6a).

**Fig. 6.**
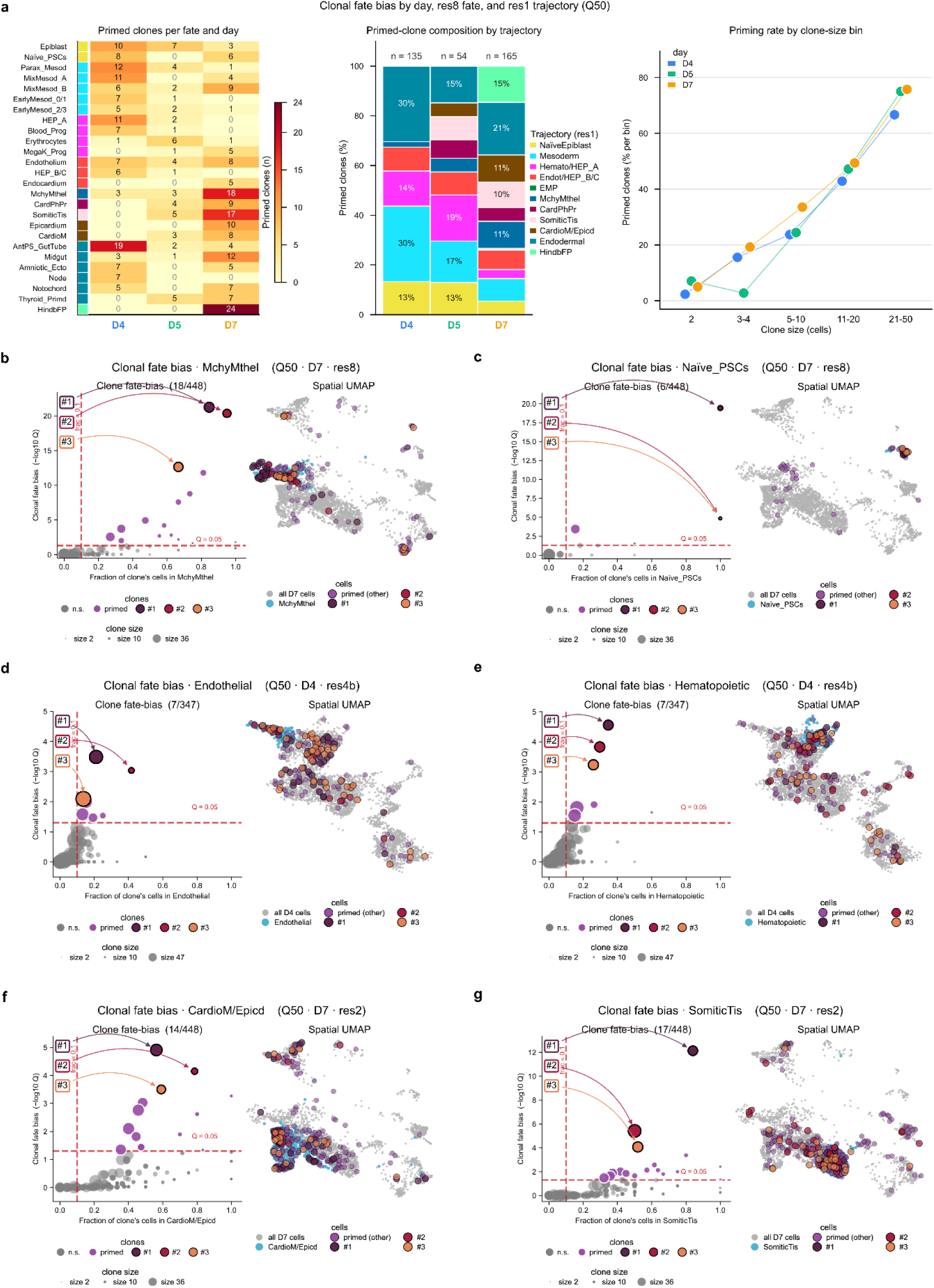
Clones primed toward individual cell fates emerge throughout the mESC differentiation. **a**, Clonal fate priming at res8 (Q50, min clone size 2). A clone is primed when CoSpar clonal_fate_bias returns BH-q < 0.05 (one-sided hypergeometric, size-stratified BH) AND fate fraction > 0.10. Left panel, primed-clone counts per res8 fate × day; rows = fates with ≥ 3 primed clones on any day, grouped by res1 parent trajectory (left strip), descending max-count within group. Middle panel, % res1-parent composition per day; totals above bars (events / clones): D4 = 135/119; D5 = 54/50; D7 = 165/139. Right panel, % primed for ≥ 1 fate vs clone size, per day (n= 347/287/448; bins with < 3 clones omitted). The monotonic rise reflects the test’s size-dependent power floor, not greater fate restriction in large clones. **b**, Clonal fate-biased clones at MchyMthel (Q50 · D7 · res8; n=448 clones, min size 2). Left panel, MchyMthel fraction vs -log10 q (BH-adjusted upper-tail hypergeometric, size-stratified BH; CoSpar clonal_fate_bias). Primed (dashed red lines): 18/448 (purple); top three highlighted as #1/#2/#3. Right, D7 UMAP, 3,636 partition cells (grey); 251 MchyMthel cells (turquoise); the 18 primed clones overlaid in purple (top three colored as in the left panel). **c-g**, as in panel b, but for additional fates: for panel c, Naïve_PSCs; d, Endothelial; e, Hematopoietic; f, CardioM/Epicd; g, SomiticTis.

The most prevalent primed fates (clones and their cell distribution that are significantly biased towards a given fate) shifted across stages (using a cross-sectional comparison, since clones do not span days); further, each time point (D4, D5, and D7) creates a separate dataset within, and has no connection, lineage-wise, between different time points): blood and endothelial programs (Hematopoietic/HEP_A, Endothelium) and Naïve/Epiblast-like were more prevalent at D4, and, by D7, the leading fates were Mesenchyme/Mesothelium, Somitic Tissues, Cardiopharyngeal Progenitors, Epicardium, and Hindbrain Floor Plate, with Naïve/Epiblast still present. Blood and endothelial trajectory priming did not disappear but shifted away from Hematopoietic/HEP_A toward mature subtypes (from D4 to D7) (Fig. 6a).

Some primed clones were strongly biased towards a single fate, the most extreme clones ending up with 85-100% of their cells in a single fate (for example, a 26-cell clone with 85% of the cells being Mesenchyme/Mesothelium, q=5×10□²²) (Fig. 6b). Clones restricted to the naïve-pluripotency state persisted at every time point (D4, D5, D7), including even a D7-clone that was 100% naïve; this provides a clone-level entry to the naïve-persistent clones examined under heritable clonal memory (see below) (Fig. 6c). In addition, we detected biased clones at both D4 and D7 time points: Endothelial- and Hematopoietic-biased clones at D4, and Cardiomyocyte/Epicardium and Somitic-biased clones at D7 (Fig. 6d-g). This analysis allows us to infer, from the clonal composition, that non-random fate priming exists in a subset of clones. Whether this is already encoded and detectable in the immediate progenitor transcriptome is investigated next.

#### Fate-biased progenitors already express the genes encoding the regulators of their eventual fate

The previous work showed, using clonal composition as the parameter, that a subset of clones is primed toward single fates. We next asked whether this priming is already detectable in the progenitor transcriptome. This could mean that a progenitor’s eventual fate can be predicted from its transcriptome, and the presence of such predictive marker transcripts may then correspond to the known, acknowledged regulators of that fate. To address this for each of the two lineage decisions detailed below, we built a clonal transition map from progenitor (t0) to (defined here as) terminal (t1) cell types, again using CoSpar (following the DARLIN framework; (L. Li et al., 2023, 2025)). This approach can predict finite-time transition maps and cellular development potential by integrating single-cell transcriptomics with lineage tracing data.

Here, we used it to assign a fate bias for the differentiated state(s) to each progenitor. For this, an analysis of differentially expressed genes (DEGs) was conducted for the most strongly committed progenitors of each branch (for details, see Methods). Committed progenitors are partitioned using a transition map that integrates clonal coupling with transcriptomic state, and CoSpar smooths the clonal signal over the cell-state graph. The differential expression is therefore not an independent test of cell state that predicts fate; rather, the lineage data supervise the direction of the map. The substantive result is that, in both documented examples (see below), the genes separating the two committed groups are known canonical, lineage-specific master regulators of the given choice, concordant with independent developmental genetics-based approaches. This indicates that progenitors can carry genuine, fate-aligned priming signatures in their transcriptome. The differential expression was applied to acknowledged (i.e., known marker gene profiles) committed progenitors (roughly 30-40% of the progenitors).

##### Example 1: HEPs do prime toward hematopoietic or endothelial fate

At D4 and D5, HEPs split into hematopoietic-biased and endothelial-biased progenitors (Fig. 7a), as observed previously (Cetin et al., 2025; Picco et al., 2026). The former exhibits the characteristic expression of key transcription factor-encoding genes of the core hemogenic program; this program (ranked by significance and fold-change) features the expression of *Runx1*, *Myb*, *Cbfa2t3*, *Nfe2l2*, *Zfpm1*, *Stat5b*, *Ikzf2*, *Ikzf1*, *Runx1t1*, *Zfp704*, *Prkar2b*, *F2r*, *Fth1,* and *Gata1*. The endothelial-biased progenitors displayed characteristic expression of *Peg3*, *Hmga2*, *Sox11*, *Etv2*, *Id3*, *Id1*, *Lef1*, *Lmo2*, *Phlda2*, *Gata6*, *Rbms1*, *Kdr*, *Amot*, and *Meg3* (Fig. 7b,c; Table S7). Importantly, these results show that fate-biased progenitors at the HEP stage already express the canonical regulatory genes that rank at the top of the acknowledged gene regulatory networks of their respective differentiated fates.

**Fig. 7.**
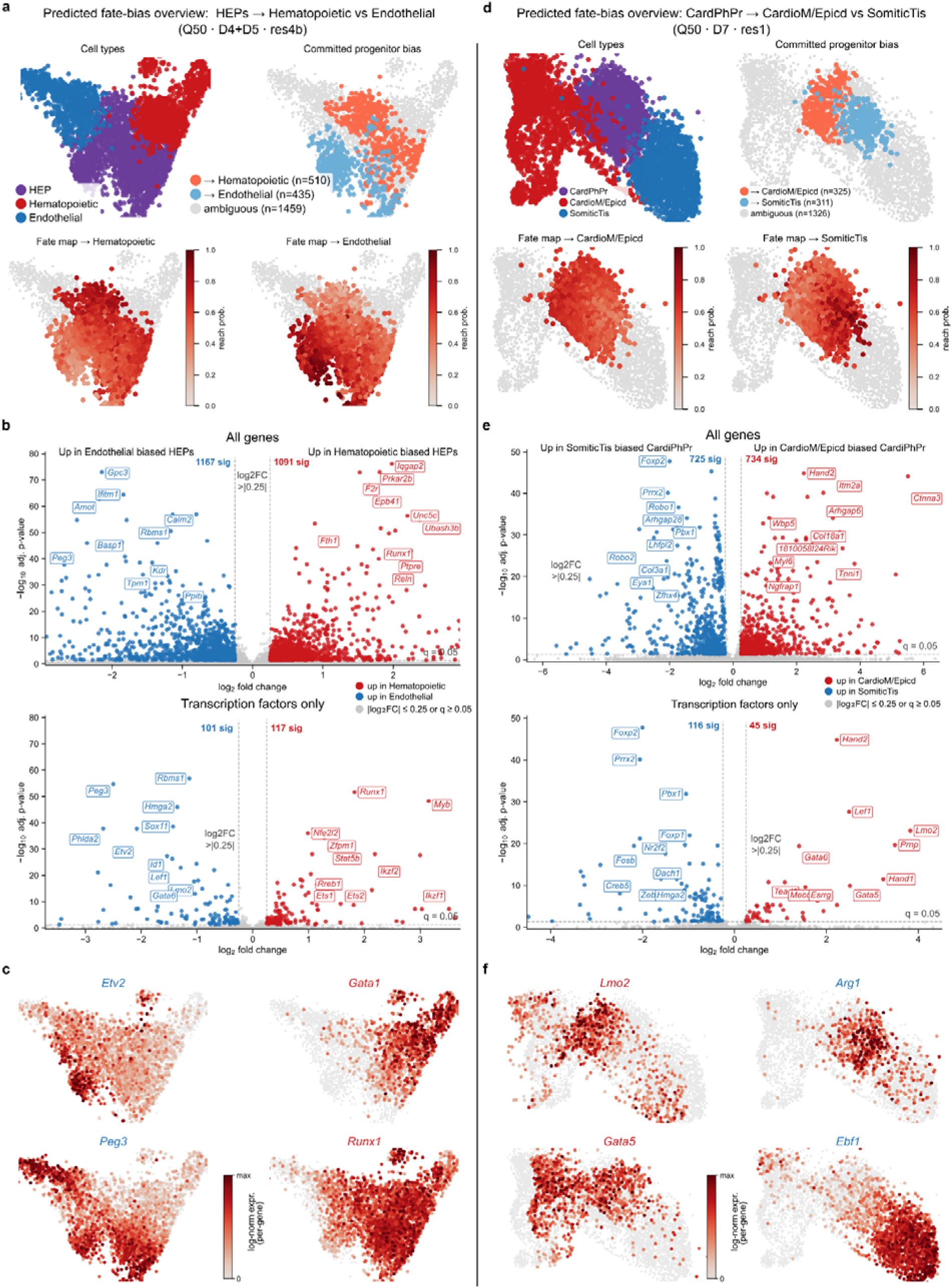
Predicted fate bias analyses identify committed progenitors and fate-discriminating transcriptional programs in two examples of mesodermal bifurcation. **a**, Overview UMAPs of HEP progenitors → Hematopoietic versus Endothelial presumptive differentiation (Q50; D4+D5; res4b). (i) Cell types: HEP progenitors (purple, t0); Hematopoietic (red, t1); Endothelial (blue, t1). (ii) Committed-progenitor bias: classification of t0 HEP cells from the within-day, clonally derived transition map: → Hematopoietic (orange, n=510); → Endothelial (light blue, n=435); ambiguous (grey, n=1,459; total t0 HEP cells = 2,404). A progenitor is committed when its predicted fate bias toward one terminal exceeds 60% (with sufficient reach probability); intermediate cells are ambiguous. (iii-iv, bottom of panel a) Per-cell fate-map probability for each terminal cell type (dark red colormap, 0 → 1). **b**, Volcano plot of Wilcoxon differential expression between committed-Hematopoietic and committed-Endothelial progenitors (ambiguous excluded; n=945 of 2,404 cells). Significance: |log fold change| > 0.25 AND BH-adjusted p < 0.05. Right, all 27,998 genes (1,091 up in Hematopoietic, red; 1,167 up in Endothelial, blue). Left, the 1,797 mouse transcription factors tested (117 up in Hematopoietic, red; 101 up in Endothelial, blue). The top-10 up- and top-10 down-regulated genes per side are labeled; ribosomal/mitochondrial gene labels are not shown, but their points are retained. **c**, Per-cell expression UMAPs for discriminating indicative genes: *Etv2* and *Peg3* (up in Endothelial), and *Gata1* and *Runx1* (up in Hematopoietic), respectively. Coordinates as in panel a. **d**, As in panel a, for CardioPharyngeal progenitors (CardPhPr) → Cardiac/Epicardial (CardioM/Epicd) versus Somitic Tissue (SomiticTis) (Q50; D7; res1): CardPhPr (purple, t0); CardioM/Epicd (red, t1); SomiticTis (blue, t1); → CardioM/Epicd n=325; → SomiticTis n=311; ambiguous n=1,326; total t0 CardPhPr = 1,962. **e**, As in panel b, for the CardPhPr bifurcation (n=636 of 1,962). Left, all genes (734 up in CardioM/Epicd; 725 up in SomiticTis). Right, TF-encoding genes (45 up in CardioM/Epicd; 116 up in SomiticTis). **f**, As in panel c, for the CardPhPr bifurcation: *Lmo2* and *Gata5* (up in CardioM/Epicd), and *Arg1* and *Ebf1* (up in SomiticTis), respectively.

##### Example 2: Cardiopharyngeal progenitors prime toward cardiac/epicardial or somitic fate

The D7 cardiopharyngeal (CardPhPr) progenitors were biased toward either the cardiac/ epicardiac (CardioM/Epicd) or the anterior somitic terminal (SomiticTis) (Fig. 7d). The cardiac-biased progenitors showed higher expression of *Hand2*, *Wbp5*, *Myl6*, *Tnni1*, *Tpm1*, *Lef1*, *Lmo2*, *Gata6*, *Tead1*, *Mecom*, *Gata5*, and *Esrrg*, while the somitic-biased progenitors do this for *Foxp2*, *Prrx2*, *Pbx1*, *Foxp1*, *Foxc1*, *Foxc2*, *Fosb*, *Nr2f2*, *Creb5*, *Dach1*, *Zeb2*, *Six1*, *Six2*, *Eya1*, *Zfhx4*, *Tle1, Arg1* and *Ebf1* (Fig. 7e,f; Table S8).

Thus, scRIPT can additionally detect progenitor fate bias. In our two examples of bifurcating lineages, DEG analysis showed that progenitor populations are heterogeneous, and their transcriptomes already express regulatory genes that also mark the differentiated end-point cell types. Because these regulatory genes are already expressed at the progenitor stage, the earlier and/or upstream master regulatory mechanisms, possibly exerted by the listed transcription factors, can now be (re)examined before such cell-type marker genes are active later in more mature cells and thereby would confound the comparison.

#### Heritable clonal memory

##### Clones carry a heritable, fate-independent transcriptional memory

In the previous section, we described fate-linked clonal differences and progenitor priming toward a fate. Next, we asked whether clones also carry fate-independent, heritable transcriptional memory, whether they have expression differences that mark a clone irrespective of which cell type(s) its cells will adopt. We sought to identify such candidate clonal memory genes using GEMLI (Eisele et al., 2024), which flags genes whose between-clone variance exceeds their within-clone variance.

We applied GEMLI systematically and extensively, running it independently within each cell type and sweeping across the three sampled time points (D4, D5, D7), and applying all annotation resolutions and all clone-calling stringencies (see above). Taking the Q50 stringency as our primary setting, we built a tier system (Tier-0 to Tier-4) to grade the robustness of each clonal memory gene, with higher tiers indicating higher confidence. Tier-0 then comprises genes present in the GEMLI memory list at least once. For each time point, we then intersected the memory-gene lists across all resolutions (Fig. 8a). This step matters most early in differentiation: at D4 and D5, many memory genes were called at every resolution except the coarsest (res1; indicated in red in Fig. 8a) and would have been missed at a single low resolution. In contrast, the effect was minimal at D7, consistent with early cell identities being less distinct and harder to separate at coarse resolution.

**Fig. 8.**
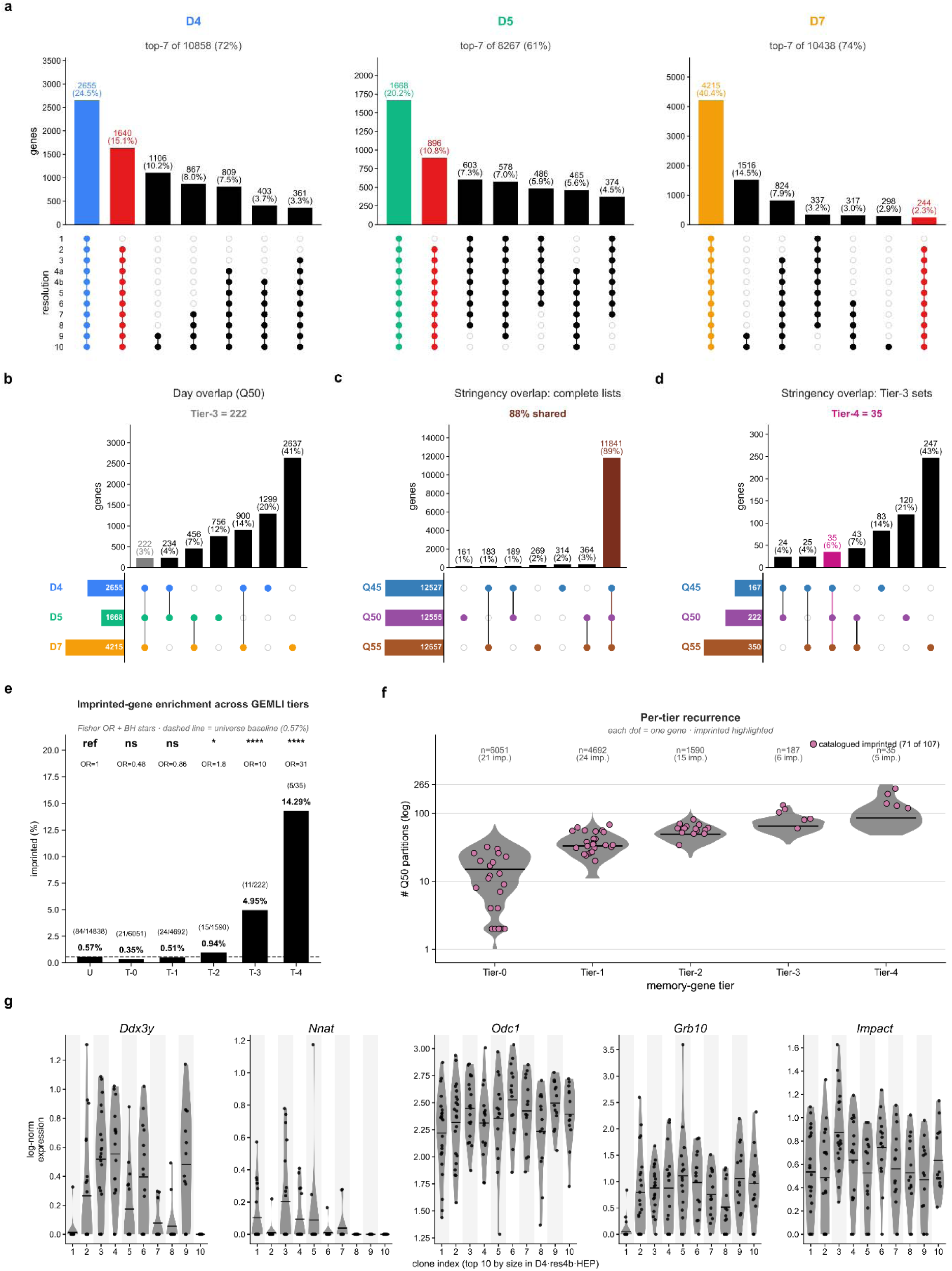
A resolution- and stringency-robust GEMLI memory-gene catalogue identifies a 35-gene cross-stringency core of genes, which is enriched for imprinted loci. **a**, UpSet plot of per-day resolution-membership patterns at Q50 (top seven patterns per day). Blue/Green/Orange, the all-resolution (resolution-robust) set; red, the pattern excluding only the coarsest resolution, res1. **b**, UpSet plot of the per-day resolution-robust sets. Genes robust on one, two, or three days define Tier-1 (4,692 genes), Tier-2 (1,590), and Tier-3 (222; grey). **c**, UpSet plot of the per-stringency Tier-3 sets; their intersection is the cross-stringency core, Tier-4 (35 genes; brown; a subset of Tier-3). **d**, UpSet plot of the per-stringency memory unions (Q45, Q50, Q55). The three-way core (dark-pink) is 11,841 genes, 88.9% of the union, indicating a stringency-robust callable pool. Because these unions pool all tiers, the overlap is dominated by background (Tier-0) calls; panel c restricts the comparison to Tier-3. **e**, Imprinted-gene enrichment per tier against the Q50-tested universe (one-sided Fisher exact; dashed line, 0.566% background). Odds ratio (OR), raw p, and significance (n.s., not significant) per tier: Tier-0, OR 0.48 (n.s., depleted); Tier-1, OR 0.86 (n.s.); Tier-2, OR 1.8 (p = 3.2 × 10□²); Tier-3 (inclusive of Tier-4), OR 10.4 (p = 4.7 × 10□□); Tier-4, OR 31.1 (p = 1.5 × 10□□). Under Bonferroni correction for five tests (α = 0.010), only Tier-3 and Tier-4 pass; Tier-2 is nominally significant only. Tier-0 depletion indicates that resolution-robustness, not memory-call membership per se, drives the signal. Asterisks, raw-p bins (*p < 5 × 10□²; **p < 1 × 10□²; ***p < 1 × 10□³; ****p < 1 × 10□□). **f**, Per-tier recurrence: n_partitions, the number of Q50 partitions (of 374) in which a gene is called memory, under the exclusive convention (Tier-3 excludes Tier-4). Reddish-purple points, the 71 catalogued imprinted genes in the Q50 memory union. Between-tier significance is not tested, as recurrence rises with tier by construction. **g**, Per-clone single-cell expression of the five most recurrent Tier-4 genes (*Ddx3y*, *Nnat*, *Odc1*, *Grb10*, *Impact*) across the ten largest clones of the D4, res4b, HEP partition. Descriptive only; illustrates within-versus between-clone variation.

Combining these per-time point lists, we grouped genes by the number of time points (as days) on which they were recovered: one day (Tier-1), any two days (Tier-2, which is the union of the D4∩D5, D4∩D7, and D5∩D7 intersections), or all three days (Tier-3; in total 222 genes) (Fig. 8b; Table S9). In addition, we intersected the Tier-3 lists from each of the three stringencies. This yielded 35 genes, designated here as Tier-4 (Fig. 8c). Notably, and independently of the tier analysis, GEMLI calls were highly robust to scar-barcode stringency. Indeed, of the union of memory genes called across the Q45, Q50, and Q55 partitions, 88.9% (11,841 out of 13,321 genes) were shared across all three (Fig. 8d).

##### Stage-persistent clonal memory may be enriched for imprinted genes

Within the candidate clonal memory gene list (see above), we also noted imprinted genes. Furthermore, this imprinted component is reproducibly a cell-type-persistent part of the clonal signal: it survives the many cell divisions between D0 (barcoding) and D4 to D7 (readouts), by which we believe any time-labile transcriptional differences would have decayed. Such persistence may signify a specific genuinely heritable, fate-independent clonal memory. Imprinted genes were indeed found enriched in the higher tiers of the candidate memory list relative to all genes detected in the data sets (Bonferroni-corrected at Tier-3 and Tier-4; Fig. 8e,f). Frequently recovered candidate clonal memory genes (from the GEMLI analysis) include *Ddx3y* (encoding a helicase, Y-chromosome-linked), *Odc1* (ornithine decarboxylase 1), *Nnat* (an imprinted gene encoding Neuronatin), *Grb10* (imprinted; growth factor receptor-bound protein 10), *Igf2* (imprinted; insulin-like growth factor 2) and *Impact* (imprinted; RWD-domain protein) (Fig. 8g).

Taken together, the GEMLI analysis and imprinted-gene enrichment suggest that scRIPT can detect a reproducible transcriptional signal whose heritable, fate-independent core may be the expression of imprinted genes.

##### Two naïve-persistent clones show distinct transcriptional profiles

We next considered clones that showed a clonal fate bias toward naïve pluripotency (see above, clonal fate bias section). Two clones indeed had a state that we refer to as Naïve-persistent: a D7 clone with 10 out of 10 cells in the Naïve_PSC state, and a D4 clone with 9 out of 10 cells in this state. Within the Naïve_PSCs, we compared the transcriptome of cells belonging to Naïve-persistent clones with that of other cells originating from clones in which 80% or more of the cells had exited the Naïve_PSCs (referred to as non-persistent) (Fig. 9a). Doing so, 28 genes were ranked as differentially expressed, 15 of which were, remarkably, in Tier-1 and Tier-2 of the clonal memory list (see above) (Fig. 9b; Table S10).

**Fig. 9.**
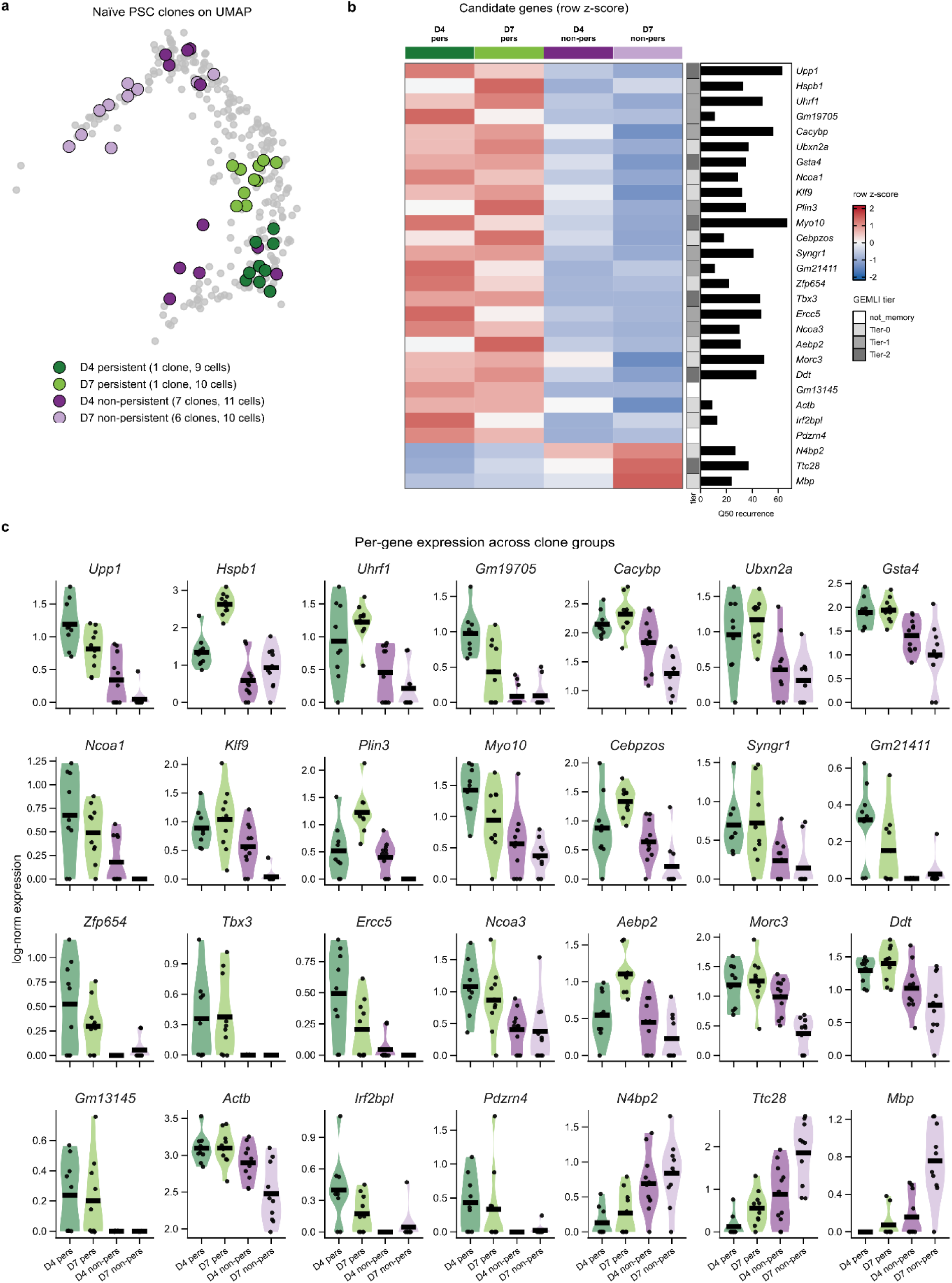
UMAP visualization and transcriptomic differences of Naïve-persistent versus non-persistent clones. **a**, UMAP location of the 2 persistent clones (d4_23M38D68M135D23M6D, D4, 9 cells, dark green; d7_43M25D56M10D17M150D, D7, 10 cells, light green) and 13 non-persistent reference clones (7 D4 clones, 11 cells, dark purple; 6 D7 clones, 10 cells, light purple), drawn from the same dataset as a within-day baseline. Highlighted points, the 40 cells from the 15 selected clones; grey, the remaining 260 Naïve_PSC cells. **b**, Heatmap of clone-mean log-normalized expression for the 28 persistence-associated genes (FDR < 0.05; Cohen’s-*d* cell-label permutation test, 10,000 permutations, Benjamini-Hochberg correction; *n=*19 cells across the 2 persistent clones vs 21 cells across the 13 non-persistent clones, pooled D4 + D7; Methods). Colour, row z-score of clone-group means (blue-white-red, clamped at ±2). Columns (left→right): D4 persistent, D7 persistent, D4 non-persistent, D7 non-persistent; the 2 non-persistent columns are the mean of clone means across the 7 D4 and 6 D7 reference clones, respectively. Rows are grouped by direction of effect: an upper block of 25 genes up-regulated in persistent clones and a lower block of 3 genes up-regulated in non-persistent clones. Per-gene right-side annotations: GEMLI tier from the Q50 memory-gene sweep (Fig. 8); Q50 recurrence, the number of Q50 partitions (of 374) in which the gene is called a memory gene; and direction of effect. GEMLI tiers: Tier-1 and Tier-2, resolution-robust at 1 and 2 of 3 days respectively, together comprising the 15 of 28 confident memory genes; Tier-0, present in the Q50 memory union but not resolution-robust at any day (background only, not counted); not_memory, absent from the Q50 sweep; no genes fall in Tier-3 or Tier-4. **c**, Per-gene single-cell distributions of log-normalized expression for all 28 candidates across the 4 clone groups. Each panel: violin plot with overlaid single-cell points (black) and the per-group mean (black crossbar). *y*-axes are independent per gene; absolute values are not comparable across panels.

Part of the 28 genes are, or might be, associated with pluripotency. These genes encompass *Tbx3* (encoding a naïve-pluripotency TF, (Niwa et al., 2009)), *Ncoa3* (Esrrb/OSN co-activator, (Percharde & Azuara, 2013)), *Uhrf1* (inner cell mass/naïve-state epigenetic regulator, (Graf et al., 2017)), *Morc3* (ERV-silencing chromatin remodeler, (Groh et al., 2021)), *Aebp2* (essential accessory subunit of the Polycomb Repressive Complex 2, (Grijzenhout et al., 2016)), *Actb* (core subunit of the esBAF/BRG1 chromatin remodeler, (Ho et al., 2009; Zhao et al., 1998)), *Ncoa1* (Esrrb coactivator, (Laan et al., 2014)), *Klf9* (transcription factor, differentiation-induced, (Yamane et al., 2018)), *Gm13145*/*Zfp978* and *Gm21411* (KRAB-ZFP proteins, (Bruno et al., 2025)), *Gm19705* (a lncRNA that, together with the lncRNA Platr22 may affect a super-enhancer of the *Zfp281* locus, (Yan et al., 2020)), and *Zfp654* (with hitherto unknown function) (Fig. 9c).

Thus, scRIPT can recover candidate clonal memory genes and, in some settings, be used to dissect the clonal differences seen in populations, such as the Naïve-persistent clones chosen here.

## Discussion

Here, we develop scRIPT as an adaptation of scLT methodology and apply it to mESC→eHEL differentiation that runs over a short period. It recovers an interpretable lineage barcode in the large majority of cells (95.1% capture, 90.7% assignment) and roughly 5,000 unique barcodes per 40,000 cells. We use the resulting clonal data to address which resulting cell types share a clonal origin, whether a progenitor’s eventual fate is already detectable in its transcriptome, and whether clones may carry heritable transcriptional memory independent of which fate(s) the cells will adopt. Barcoding occurs before the onset of differentiation rather than at a defined, progressed stage. The extensive analyses are meant to demonstrate what scRIPT makes measurable rather than to settle the underlying developmental biology mechanisms of this early hematoendothelial differentiation.

Methodologically, scRIPT’s gains come from decoupling the barcoding step from the cell differentiation and from removing a major source of barcode dropout before sequencing. Delivering Cas9 as an RNP collapses the days-long induction (and degradation) window of inducible systems into a brief, efficient pulse, concentrating the scarring at a narrower point rather than across the experiment. Because destabilized eGFP acts as a GFP-level-dependent sensor for the state of the TA, cells that have lost a capturable barcode can be removed by FACS before differentiation. A simplified, 10X-compatible two-step PCR and a hybrid PE150/PE250 sequencing strategy further reduce cost and turnaround time. Equally important, scRIPT introduces explicit QC metrics for lineage data (scar entropy and complexity, D0 and cumulative scar counts, and a transcriptome-independent scar-doublet score). It also goes together with a set of scar-calling stringencies that identify problematic barcodes and doublets from the lineage data themselves. Hereby, scRIPT addresses a gap that has made lineage results difficult to compare across studies.

We apply this in fate coupling and find that scRIPT recovers the expected within-trajectory relationships from lineage information alone (for example, the coupling of hematopoietic and endothelial cell types with their shared HEPs), indicating that the method captures a real clonal signal. It also recovers couplings between compartments, notably, as shown here, between the hematoendothelial and Mesenchyme/Mesothelium trajectories, consistent with the dual-outcome precursors (i.e., hemangioblast, mesenchymoangioblast, hematomesoblast) described *in vivo* (Biben et al., 2023). These couplings must, however, still be interpreted in light of when the barcodes are introduced, for scarring precedes, in our case, a stage comparable to germ-layer specification. Thus, lineages that diverge later can co-occur within a clone through shared early ancestry, which the available filters do not remove. A significant coupling, therefore, reflects a real clonal connection but not, on its own, a specific late progenitor. Accordingly, we read couplings that match known developmental relationships or independent lineage data as those relationships. Also, we treat couplings that cross germ layers without developmental precedent conservatively as shared early ancestry. Introducing barcodes at a defined later stage (likely by other means than RNP-mediated transfer used here), if efficient enough and not occurring too slowly, and sampling at closely spaced time points, would narrow this shared ancestry window.

A second question was then whether scRIPT could detect fate bias already encoded in the progenitor transcriptome. For this, we examined two examples of bifurcations in mesodermal cells: HEPs splitting toward hematopoietic versus endothelial fate, and Cardiopharyngeal Progenitors toward cardiac versus somitic fate. For each, the genes distinguishing the committed progenitors of each branch encompassed canonical, lineage-specific regulatory genes of that choice (for example, *Runx1* and *Myb* on the hematopoietic, and *Etv2* and *Kdr* on the endothelial side; *Hand2* and the *Gata* factors versus *Foxc1/2*, *Six1*, *Eya1*, and *Pbx1* for the cardiac versus somitic decision). This concordance with players previously established through developmental genetics is the substantive point. However, it carries a caveat: because the transition map (CoSpar) integrates clonal coupling with transcriptomic state, the differential expression is not an independent demonstration of state predicting fate. Rather, the lineage data supervise the direction of the map, and the recovered priming genes are the expected regulator-encoding genes, not arbitrary ones. Hence, scRIPT offers a practical handle on studying early regulation in an unprecedented way. Indeed, because expression of these regulatory genes is already detected at this resolution at the progenitor stage, the upstream programs that would bias fate can be examined early, prior to their presence and direct involvement in (more) mature cell types, which would confound the studies.

The clonal data also reveal two kinds of transcriptional memory. The most reproducible component is fate-independent, and possibly dominated by imprinted genes; these latter genes are significantly overrepresented among the stage-persistent memory genes, showing that scRIPT can recover such signals. And as presented, a second, state-linked signature distinguishes (the very few) clones that remain in the naïve-pluripotency state. The naïve-persistence genes, based on the aforementioned two-clone comparison, will need independent confirmation in further work. Finally, the fraction of clones we can call fate-biased is a detection floor set by clone-size-dependent power rather than a count of committed clones, so unprimed calls in small clones are uninformative. Several limitations contextualize these conclusions and delineate the optimal scope of the method. The system used here is *in vitro* mESC differentiation, and barcoding before cell differentiation produces a wide shared-ancestry window, which limits how finely couplings can be attributed to specific progenitors. Annotations of the continuously differentiating cells are sensitive to clustering resolution, which we mitigated by analyzing 11 resolutions but did not eliminate this issue. A minority of cells still carried barcodes that could not be fully resolved, a fraction that, despite our deep sequencing, might be reduced by deeper and/or long-read sequencing.

This detection floor is a consequence of a more general constraint, i.e., scRNA-seq lineage tracing recovers only a fraction of each clone’s progeny. Inefficient cell capture, dissociation-dependent bias against particular cell types, and barcode dropout each reduces the recovered fraction, and variable division rates across the state manifold further skew apparent clone sizes. In our experiment, the time window from the post-scarring/sort (D0) to capture (up to D7) allows up to about 10 cell divisions, considering a 16-hour mESC doubling time (which would result in far more progeny than we recovered). However, the true per-clone expansion is lower, as the cells slow their cell cycle on entering differentiation. Both simulations of CRISPR-based recorders (Salvador-Martínez et al., 2019) and recent large-scale *in vivo* lineage-tracing datasets (Koblan et al., 2025; L. Li et al., 2023) indicate that reconstruction accuracy and clonal coverage decline under realistic capture loss, placing our per-clone losses within current field norms. After restricting to the highest-confidence lineage barcodes (found here as Q50 stringency; see Methods), the cells recovered per clone were correspondingly low: a median of 2 (with ranges from 1 to 14) at the cardiopharyngeal-progenitor stage and a median of 5 (range 2-29) across progenitor and “terminal” compartments. We therefore treat assignments of fate as conditional on the cells observed: a clone scored as “uncommitted” may have produced “terminal” cells that went uncaptured.

With efficient barcoding at defined stages and denser sampling permitted, the same platform should be well suited to the *in vitro* applications that motivated it, including drug-response and resistance studies, and cell transition models for tissue regeneration. Taken together, in its present form, scRIPT provides an accessible, high-coverage, quality-controlled scLT platform optimized for *in vitro* cell differentiation for short-term differentiation (up to 7 days), and a clear example of how lineage and transcriptome can be combined to ask where in development a cell’s fate and its heritable identity are detectably set.

## Methods

### Generation of the knock-in mESC line

The donor template sequence and primer pairs are provided in Table S2. An sgRNA targeting the sequence GGTGTTATGGCAGTAACTTTAAT was designed and cloned into the previously described Cas9-sgRNA-containing plasmid. Transfection and detection of heterozygous knock-in clones were performed as previously described. mESC culture conditions and practices were as previously described. The selected clone, 2E12, was tested for differentiation into eHELs, and the results were assessed by flow cytometry for surface-marker expression; the antibody panels and detailed protocol have been described previously (Cetin et al., 2025; Picco et al., 2026; Staalduinen et al., 2026).

In the transgene cassette, between d2eGFP (X. Li et al., 1998) and the CARLIN TA, the primer-binding site is set for amplification of the lineage libraries after scRNA-seq capture. The primer-binding site largely overlaps the destabilization/degradation domain that originated from mouse ornithine decarboxylase and will create an alternative binding site in the mouse genome. For this reason, the coding sequence of the destabilization domain that the primer binds is altered, but the amino acid sequence is not.

### CARLIN barcoding and single-cell collection

Ten CRISPR RNAs (crRNAs) targeting ten sites in the CARLIN array were prepared as ribonucleoprotein (RNP) complexes and transfected using Lipofectamine CRISPRMAX Cas9 Transfection Reagent (CMAX00008). RNP preparation and cationic-lipid transfection followed the manufacturer’s instructions; details have been described previously (Cetin et al., 2025). Approximately 30 hours after transfection, the transfected pool was pre-selected for cells with increased GFP expression (Fig. S2). A sample was taken for the D0 scRNA-seq experiment, and the remaining cells were used for differentiation as previously described (Cetin et al., 2025). On days 4, 5, and 7 (D4, D5, D7), approximately 10 to 20 embryoid bodies were selected and dissociated.

### Transcriptome library preparation and sequencing

The dissociated cells were loaded onto the 10x Genomics Chromium X platform, and transcriptome libraries were generated using the Chromium Next GEM Single Cell 3’ Reagent Kit v3.1 (Dual Index) (see User Guide, CG000315 Rev F). Library preparation, sequencing, and raw-data processing followed the manufacturer’s instructions.

### Lineage barcode library preparation

Lineage barcode library preparation began after Protocol Step 2.2 (cDNA Amplification). An aliquot of the amplified cDNA was taken, and 100 ng was used as input for the first PCR, which amplified the TA. The primers are listed in Table S2C. Two polymerases were used: Phusion High-Fidelity DNA Polymerase (M0530L) and KAPA HiFi HotStart ReadyMix (07958927001). For Phusion, the cycling conditions were 98 °C for 1 min; 15 cycles of 98 °C for 20 s, 67 °C for 30 s, and 72 °C for 20 s; and a final extension at 72 °C for 1 min. For KAPA HiFi HotStart ReadyMix, the conditions were 98 °C for 35 s; 15 cycles of 98 °C for 20 s, 67 °C for 30 s, and 72 °C for 20 s; and a final extension at 72 °C for 1 min. The resulting PCR products were purified using 1.5x SPRIselect reagent (B23318). The protocol then continued with the indexing step (Protocol Step 3.5, Sample Index PCR) recommended for the Chromium Next GEM Single Cell 3’ Reagent Kit v3.1 (Dual Index). Indexed libraries were sequenced on a NovaSeq X (150 bp paired-end) and a NovaSeq SP (250 bp paired-end).

### Lineage Data Processing and Metrics

#### CARLIN

The raw reads of one 10xGenomics RNA-seq library trimmed and mapped with STAR solo (Dobin et al., 2012) with v3 10xGenomics settings to the constructed mm10 reference file including the supplied custom CARLIN construct. Next, molecules are assigned to genes, quality filtered, and deduplicated using SingleCellMultiOmics (https://github.com/BuysDB/SingleCellMultiOmics; bamtagmultiome.py). STAR is used for deduplicating reads and for generating the count table. The count table is loaded with the Python package scanpy (Wolf et al., 2018) for filtering and normalization. A gene needs to be present in a minimum of 3 cells, and a cell needs to have a minimum amount of 200 genes containing counts. Ribosomal, mitochondrial, and *Malat1* genes are removed. Counts are normalized to a million reads and log-transformed.

#### scRIPT preprocessing

Long and short reads of 10x Genomics RNA-seq libraries were processed and analyzed as follows: Reads were trimmed of their poly-A tail and demultiplexed using a custom script (*demux.py*) (https://github.com/BuysDB/scRIPTpreprocessing) (Barbanson, 2025). Only reads with a hamming distance of at most 2 between the primer sequence and the expected sequence are retained. Reads were trimmed of their adapters using FastP (Chen et al., 2018) with default settings. The trimmed reads were then mapped with Minimap2 (H. Li, 2018) to a custom reference genome, which includes the mm39 genome and the *Tigre* locus. To determine the final scar alignment, reads mapped to the *Tigre* locus were re-aligned using the Smith-Waterman alignment algorithm, which was adapted from swalign (https://github.com/mbreese/swalign). A biased scoring function was incorporated into the original code to prioritize alignment of the primer at the expected location without gaps. This is implemented by scoring the first 20 bases of both the reference sequence and the read using a −200 gap extension and a −200 gap penalty.

The mapped data from multiple short-and long-read libraries were merged by finding whether shorter alignments could be uniquely associated with a long alignment. Doublets were removed using a K-fold cross validation using a random forest classifier, with the following important features: total number of reads for the second most common scar, total number of unique sequences for the second most common scar, fraction of reads assigned to the second most common scar, fraction of reads assigned to the most common scar, number of components in the Hamming graph constructed from all sequences found in that cell. The transcriptome scDblFinder labels were used for fitting the random forest. Further details of the code can be found at (https://github.com/BuysDB/scRIPTpreprocessing) (Barbanson, 2025).

#### Scar-level QC metrics

A lineage-only doublet signal that flags a cell carrying more than one lineage barcode (scar); it agreed with the transcriptome-based scDblFinder call for 93% of cells (see Fig. 3a). The pipeline records per-cell diagnostic metrics (‘most_common_scar_n’, ‘matching_scar_n’, ‘distinct_scar_n’, ‘fraction_common’, ‘cardinality_distinct’) and a doublet posterior probability, ‘scar_posterior_doublet’. Cells were kept as singlets when ‘scar_posterior_doublet < 0.45’ and ‘scDblFinder_class == “singlet“’; cells failing either criterion were treated as doublets.

Assessed on unique scars (separately for PE150 and PE250) by parsing the CIGAR-like SC string: deletion and insertion events were counted and their lengths summed, giving for each scar a deletion count, an insertion count, a cumulative deletion length, a cumulative insertion length, and a mutation class (deletion-only, insertion-only, both, or no mutations); wild-type strings (‘WT’, ‘1WT’) were treated as zero-event scars. From these, each scar’s total mutation-event count (‘total_events’, deletions plus insertions) and cumulative indel length (‘total_mutation_length’, summed deletion and insertion lengths) were derived. The scar complexity score (‘scar_complexity_score_uniquè, see Fig. 3c) is the total mutation-event count; higher values reflect more extensive editing and flag scars unlikely to be partial.

Quantile cut-offs applied to the per-day cells-per-scar distribution to remove over-represented scars, each named for the percentile used as its cut-off. Q100 applies no cut-off (all scars retained); Q55, Q50, and Q45 progressively remove the highest cells-per-scar scars, so Q45 is the most stringent. Because cut-offs were computed before doublet removal and further filtering, realized retained fractions exceed the nominal quantiles (largest shift Q55: 60.1% vs 55%; see Fig. 3d,f).

Thresholds were computed on the combined D0-D7 data (doublets retained) after excluding wild-type scars: for each stringency (percentiles 0.45, 0.50, 0.55) and each day, the threshold was that percentile of the non-zero cells-per-scar values for that day. A scar was retained at a given stringency only if its cells-per-scar was strictly below the threshold (or zero) on every day (D0, D4, D5, D7); a scar at or above the threshold on any day was removed. The uniquely aligned variants (‘scar_q45_uniq’, etc.) additionally require ‘n_unique_alignments > 0’.

##### Scar Entropy

Scar entropy quantifies how widely a lineage barcode is distributed across transcriptionally distinct cell populations, and was used as a diagnostic for barcode homoplasy. For each sampling day (D0, D4, D5, D7) independently, cells called singlets from the lineage data (scar_posterior_doublet < 0.5) were cross-tabulated by their assigned scar and by that day’s preliminary transcriptome clustering, giving for each scar a vector of cell counts (n□ … n_K) over the K clusters of that day. The Shannon entropy of that vector was computed as H = −Σ□p□ ln p□, with p□ = n□ / Σ□n (natural logarithm, scipy.stats.entropy, which normalises the count vector internally; clusters containing none of the scar’s cells contribute zero). A scar confined to a single cluster therefore has H = 0, and H rises towards ln K as its cells spread evenly over more clusters, so that exp(H) can be read as the effective number of clusters a scar occupies. Each scar was assigned the maximum of its four per-day entropies, i.e. the value for the day on which that scar was most widely dispersed. Entropy was computed only for scars observed in more than five singlet cells pooled across days; scars below this threshold were left undefined. Because entropy is a property of the scar rather than of the cell, the value was joined back onto every cell carrying that scar (column scar_entropy, Tables S3 and S5).

A high entropy identifies a barcode whose cells are scattered across unrelated transcriptional states, the expected signature of an identical Cas9 repair outcome arising independently in several founder cells (barcode homoplasy) rather than of a single genuine clone. Scar entropy was used descriptively (Fig. 3c,d) and to select low-entropy scars for the per-scar visualisations (H < 1.6, i.e. fewer than about five effective clusters). It was not applied as a filter to the datasets used for the fate-coupling, fate-bias and memory-gene analyses, which were defined solely by the cells-per-scar quantile stringencies described above.

##### Scar Doublet Score

Scar doublet score. The scar doublet score (scar_posterior_doublet) estimates, from the lineage library alone, the probability that a droplet contained more than one cell. For each cell, the distinct target-array sequences recovered from the lineage library were summarised into read and sequence tallies: the reads supporting the most abundant scar, the reads and the number of distinct sequences either matching or differing from it, the fraction of reads carrying the most abundant scar, the long- and short-read counts, and the uniquely and ambiguously aligned read counts. These were combined with features of a per-cell sequence-similarity graph, in which nodes are the cell’s distinct sequences (at most 150) and an edge joins any sequence lying within a Hamming distance of 2 of a sequence carrying at least 5% of the cell’s reads, so that sequencing-error variants collapse onto their parent sequence. The total read count and the number of sequences of each of the three largest connected components were retained, together with the share of graphed reads held by the second-largest component. A cell derived from a single nucleus is expected to yield one dominant component, whereas a droplet containing two cells yields two substantial components.

A random-forest classifier (400 trees, maximum depth 5) was trained on these lineage-derived features together with a library identifier, using the transcriptome-based scDblFinder class as the target label. Each cell’s score is its out-of-fold predicted doublet probability under five-fold stratified cross-validation (cross_val_predict, scikit-learn), so that no cell contributes to the model that scores it. The model was fitted only on cells whose most abundant scar was supported by more than 10 reads; cells below this threshold were left unscored. Out-of-fold predictions agreed with the scDblFinder call for 93% of cells, with 6% called doublet by scDblFinder alone and 1% by the scar data alone (Fig. 3a).

Because every feature derives from the lineage library, the score provides a lineage-based doublet signal that can be recomputed without repeating transcriptome doublet detection. The classifier is, however, supervised on scDblFinder labels, so the score is calibrated to the transcriptome-based call rather than independent of it. In the per-library preprocessing, cells were retained when scDblFinder classified them as singlets and their scar doublet score did not exceed 0.45; cells that were left unscored were retained.

#### Transcriptome Data Processing

Eight 10x Genomics libraries were generated across four differentiation time points of the scRIPT lineage-tracing mESC system: one sample at D0, one at D4, three divided samples at D5 (D5a, D5b, D5c) and three divided samples at D7 (D7a, D7b, D7c). For each library, Cell Ranger (Zheng et al., 2017) filtered_feature_bc_matrix and raw_feature_bc_matrix outputs were imported into R 4.5.0 with Seurat::Read10X (Hao et al., 2024), and both matrices were converted into Seurat v5 objects using CreateSeuratObject with min.cells = 0 and min.features = 0, so that no additional gene-or cell-level filtering was applied at object construction. The trailing “-1” barcode suffix produced by Cell Ranger was replaced with a lowercase sample identifier (_d0, _d4, _d5a, …) on both the filtered and the raw matrices to ensure globally unique cell IDs when the libraries were later merged, while preserving droplet correspondence between the two matrices for downstream ambient-RNA correction. Sample and condition annotations were added as metadata columns. Lineage-doublet posterior probabilities inferred from the paired scRIPT lineage-tracing analysis (Table S3) were joined to the transcriptome metadata as scar_posterior_doublet. Standard QC metrics were then computed: total UMI count per cell (nCount_RNA), number of expressed genes per cell (nFeature_RNA), the percentage of mitochondrial reads (PercentageFeatureSet, pattern ^mt-) and the percentage of ribosomal-protein-gene reads (pattern ^Rpl/^Rps).

Because per-cell UMI and feature distributions differ markedly between the pluripotent D0 sample and the more differentiated D4-D7 samples, QC thresholds were tuned per library by visual inspection of violin and scatter plots. Filtering was applied in two passes: before doublet detection, cells were retained if they satisfied library-specific lower bounds on nCount_RNA and nFeature_RNA together with band filters on percent.mt and percent.ribo; after doublet removal, a second pass applied library-specific upper bounds on nCount_RNA and nFeature_RNA to eliminate residual high-UMI outliers. The retained ranges (nCount_RNA, nFeature_RNA, percent.mt, percent.ribo; “no upper bound” indicates that the corresponding second-pass filter was not applied) were as follows. D0: 4,500-23,000, 1,900-5,000, 0.1-6%, 15-35 %. D4: 10,000-70,000, 3,700-9,000, 1-6 %, 10-31 %. D5a: >30,000 (no upper bound), >6,300 (no upper bound), 1-7 %, 8-22 %. D5b: 27,000-120,000, >5,900 (no upper bound), 1-7 %, 7-23 %. D5c: >9,000 (no upper bound), >3,200 (no upper bound), 0.7-7 %, 14-32 %. D7a: 6,000-50,000, 2,700-7,000, 1-11 %, 8-28 %. D7b: 6,000-45,000, 2,600-7,500, 1.3-8 %, 6-26 %. D7c: 15,000-90,000, 4,700-9,500, 1.3-7 %, 6-25 %. The two orthogonal doublet filters, retention of only cells classified as singlets by scDblFinder, and removal of cells with a scRIPT posterior doublet probability > 0.45, were applied identically to all eight libraries.

To provide cluster-aware inputs for both doublet detection and ambient-RNA correction, filtered cells were normalized with NormalizeData, 4,000 highly variable genes were selected with FindVariableFeatures (vst method), and the object was scaled and PCA-reduced to 50 principal components. A k-nearest-neighbour graph was built with FindNeighbors (dims = 1:50, k.param = 10, n.trees = 1000) and a preliminary Leiden clustering (Traag et al., 2019) was performed at resolution 1.2 (FindClusters, method = “igraph”, algorithm = 4); this preliminary clustering was used as the cluster input for both the doublet-detection and the ambient-RNA-correction steps below. Transcriptomic doublets were identified with scDblFinder (Germain et al., 2021) v1.22.0 in cluster-aware mode (clusters = the preliminary Leiden clustering, iter = 10, k = 10, artificialDoublets = 5 × n_cells), and only cells labeled as singlets were retained. Cells whose scRIPT posterior doublet probability exceeded 0.45, corresponding to droplets in which two distinct lineage barcodes were co-detected, were additionally removed, providing an orthogonal, lineage-based doublet filter. The library-specific upper-bound pass on nCount_RNA and/or nFeature_RNA (Quality-control filtering, above) was then applied.

Ambient RNA was corrected with SoupX (Young & Behjati, 2020) v1.6.2. Filtered and raw droplet matrices were combined into a SoupChannel (calcSoupProfile = FALSE, keepDroplets = TRUE), the soup profile was estimated with estimateSoup, and the preliminary Leiden clusters were supplied via setClusters. The contamination fraction was inferred with autoEstCont, and integer-rounded corrected counts were generated with adjustCounts (roundToInt = TRUE). Original counts were retained in a dedicated layer, and the corrected counts replaced the default counts layer for all downstream analyses.

Cell-cycle scores were assigned with Seurat’s CellCycleScoring using mouse orthologues of the Tirosh S-and G2/M-phase gene lists (Tirosh et al., 2016) (as updated in Seurat’s cc.genes.updated.2019 object), obtained by mapping human symbols to *Mus musculus* orthologues with gprofiler2 (Kolberg et al., 2023) (gorth, source_organism = “hsapiens”, target_organism = “mmusculus”); S.Score, G2M.Score and Phase were stored in the metadata.

For dimensionality reduction and clustering, non-informative genes were excluded (mitochondrial ^mt-, ribosomal ^Rpl/^Rps and hemoglobin ^Hbb/^Hba genes) together with genes carrying fewer than 16 total UMIs across the library. On the resulting gene set, counts were log-normalized, 4,000 highly variable features were selected (vst), and data were scaled with regression of the technical and biological confounders nFeature_RNA, nCount_RNA, percent.mt, percent.ribo, S.Score and G2M.Score. Principal-component analysis was run with 50 components, a shared-nearest-neighbor graph was constructed (k.param = 10, n.trees = 1000, dims = 1:50), a final Leiden clustering was performed at resolution 1.2, and a UMAP embedding (McInnes et al., 2018) was computed on the same 50 principal components with n.neighbors = 10.

The seven post-QC differentiation-day Seurat v5 objects (D4, D5a-c, D7a-c; D0 was processed separately) were combined with merge() and DietSeurat() to strip inherited reductions. Mitochondrial (^mt-), ribosomal (^Rps/^Rpl) and hemoglobin (^Hba/^Hbb) genes, together with genes carrying fewer than 16 total UMIs across the merged object, were removed. Counts were log-normalized (NormalizeData), 4,000 highly variable genes were selected (FindVariableFeatures, vst), and expression was scaled with regression of nFeature_RNA, nCount_RNA, percent.mt, percent.ribo, S.Score and G2M.Score. RunPCA (npcs = 50) produced the merged_pca reduction.

Harmony (Korsunsky et al., 2019) v1.2.3 was applied to merged_pca through Seurat v5’s IntegrateLayers (method = HarmonyIntegration) at package defaults, producing the harmony reduction. For both merged_pca and harmony, shared-nearest-neighbour graphs were built with FindNeighbors (dims = 1:50, k.param = 10, n.trees = 1000) and cells were clustered with the Leiden algorithm (FindClusters, algorithm = 4) at a primary resolution of 0.8 together with a sweep of {0.5, 1, 1.5, 2, 2.5, 3, 3.5, 4, 4.5, 5, 10}. An additional Leiden clustering of the harmony embedding at resolution 2.75 was computed and retained as the working partition used to organize the cell-type annotation. The final harmony_umap was computed with RunUMAP (reduction = “harmony”, dims = 1:50, n.neighbors = 10, min.dist = 0.3, spread = 0.5, local.connectivity = 1, n.epochs = 1000); a diagnostic merged_umap from merged_pca, with matching dims and n.neighbors, supported before/after batch-removal comparison by sample of origin. Later, per-sample count layers were re-joined with JoinLayers.

##### Cell-type and developmental-stage annotation by projection onto the mouse gastrulation atlas

Query cells from the scRIPT dataset were annotated by projecting them onto a mouse gastrulation reference atlas using a mutual-nearest-neighbor (MNN)-based label-transfer approach (Imaz-Rosshandler et al., 2023). The reference was the extended single-cell atlas of mouse gastrulation and early organogenesis; cells spanning embryonic stages E6.5-E9.5 were used, and each reference cell carried two independent cell-type annotations that were transferred to the query in parallel: the original annotation from Pijuan-Sala et al.(Pijuan-Sala et al., 2019) and the refined, higher-resolution annotation introduced with the extended atlas (Imaz-Rosshandler et al., 2023). This yielded two mapped cell-type labels per query cell, each with its own confidence score, alongside a mapped developmental stage. Raw query counts, cell barcodes, gene identifiers and per-cell metadata were exported from the processed Seurat object and reloaded as a SingleCellExperiment (Amezquita et al., 2020); the reference was loaded as a SingleCellExperiment, log-normalized, and restricted to genes with a total count above 15 across the reference. Reference Ensembl gene identifiers were mapped to gene symbols using Ensembl release 87 (GRCm38) annotation.

Genes shared between query and reference were retained, from which likely uninformative or confounding features were removed prior to mapping: RIKEN cDNA and predicted (Gm) gene models, mitochondrially encoded genes, ribosomal-protein genes of the Rps and Rpl families, the hemoglobin genes Hba and Hbb, imprinted genes, the X-inactivation-associated transcripts *Xist* and *Tsix*, and all genes located on the Y chromosome. From the remaining genes, the 4,000 with the highest pseudobulk expression variance in the reference were selected as mapping features.

For mapping, reference and query counts were concatenated and jointly log-normalized across the two datasets with multiBatchNorm (without cosine normalization), and a shared 40-dimensional principal-component space was computed on the selected features with multiBatchPCA (batch = reference/query). The reference was first internally harmonized across its constituent samples by hierarchical MNN correction (reducedMNN), with samples merged in order of decreasing developmental stage (E9.5 → E6.5) and, within each stage, decreasing cell number. Query cells were then aligned to the batch-corrected reference by a second reducedMNN step. Each query sample was mapped independently. All batch-correction steps were performed with the batchelor package (Haghverdi et al., 2018), and random seeds were fixed for reproducibility (global seed 1234; batch-correction seed 42).

In the aligned principal-component space, each query cell was assigned the original cell type, extended cell type, and developmental stage of the majority of its k = 15 nearest reference neighbors (queryKNN, BiocNeighbors), with ties resolved in favor of the closest neighbor. For each of the two annotation systems, a per-cell mapping confidence score was defined as the fraction of the k neighbors supporting the assigned label. The single closest reference cell was also recorded for each query cell, and its atlas UMAP coordinates were transferred to the query cell for visualization on the reference embedding.

To visualize query and reference cells in a common embedding, the mapping was additionally run on the full query with per-dataset cosine normalization enabled, and a joint UMAP was computed from the combined batch-corrected principal-component space (n_neighbors = 25, min_dist = 0.3).

##### Cell-type annotation by Seurat anchor-based label transfer

As an alternative annotation strategy, the extended cell-type labels of the reference atlas were transferred to the scRIPT cells using the anchor-based label-transfer framework of Seurat. The reference was the extended single-cell atlas of mouse gastrulation and early organogenesis, loaded as a Seurat object; to reduce stage imbalance and computational cost, each developmental stage was randomly downsampled to 7,455 cells, matching the smallest stage (mixed_gastrulation). The query was the scRIPT dataset. For reference, the per-cell percentages of mitochondrial and ribosomal-protein transcripts were quantified. Both reference and query expressions were log-normalized (Seurat LogNormalize).

Genes common to query and reference were retained, from which mitochondrially encoded genes, ribosomal-protein genes of the Rps and Rpl families, the haemoglobin genes Hba and Hbb, a curated set of imprinted genes, the X-inactivation-associated transcripts *Xist* and *Tsix*, and all genes located on the Y chromosome (identified from Ensembl annotation, EnsDb.Mmusculus.v79, GRCm38) were removed; in contrast to the MNN-based mapping, RIKEN cDNA (Rik) and predicted (Gm) gene models were deliberately retained, as several displayed cell-type-specific expression. Genes with a total count below 16 in either the reference or the query were then excluded, and both objects were subset to the resulting shared gene set.

For each highly-variable-gene count, variable genes were selected in the reference (FindVariableFeatures, default variance-stabilizing-transformation method), and reference expression was scaled with regression of technical and biological covariates, total counts per cell, number of detected genes, percentage of mitochondrial and of ribosomal-protein transcripts, and previously computed S-and G2M-phase cell-cycle scores (ScaleData). Principal-component analysis was then computed over the variable genes (RunPCA). Transfer anchors between reference and query were identified by projecting the query into the reference principal-component space (FindTransferAnchors with reference.reduction set to the reference PCA; the Seurat “pcaproject” method, i.e., without joint integration of the two datasets), and the extended cell-type annotation was transferred with TransferData. For each query cell, this yielded a predicted cell-type label and an associated transfer confidence score (the maximum per-cell prediction score).

To assess the sensitivity of the annotation to feature-selection and dimensionality choices, the procedure was repeated across combinations of 4,000, 7,000, and 10,000 highly variable genes with 50, 75, and 100 principal components. Labels were transferred independently for each combination and retained as separate annotations, and the resulting distributions of maximum prediction scores were compared across settings. Transferred labels were visualized on the query’s Harmony-derived UMAP embedding.

##### Positive marker genes for cell types

Positive marker genes for each cell cluster at all eleven annotation resolutions (res1-res3, res4a, res4b, res5-res10). Markers were identified per resolution with Seurat’s FindAllMarkers (Wilcoxon rank-sum test; logfc.threshold = 0.25, min.pct = 0.1, only.pos = TRUE), using the res8 clustering shown for illustration. For each resolution, genes were filtered to those with a Benjamini-Hochberg-adjusted *P* < 0.01 and a positive average log fold-change, and the top 100 per cluster by average log fold-change were retained. Each resolution is provided as a separate sheet reporting, per marker, the cluster, gene symbol, average log fold-change, the fraction of cells expressing the gene within (pct.1) and outside (pct.2) the cluster, and the raw and adjusted *P* values.

#### Lineage Tracing Analysis with CoSpar

##### Clonal Fate Coupling

Clonal fate coupling was computed independently within each analysis partition, defined as a unique combination of barcode-calling stringency (Q100, Q55, Q50, Q45), sampling time point (D4, D5, and D7), and cell-type annotation resolution (eleven levels: res1-res3, res4a, res4b, and res5-res10), giving 132 partitions in total. Within each partition, a one-hot (binary) cell-by-clone assignment matrix was constructed and used to initialize a CoSpar object (Wang et al., 2022) (cs.pp.initialize_adata_object). Singleton clones, which cannot contribute to coupling, were removed (cs.tl.filter_clones, clone_size_threshold = 2, filter_larger_clones = False), following the DARLIN analysis workflow (L. Li et al., 2023).

Fate coupling was then computed between all pairs of cell types represented by surviving clones (cs.tl.fate_coupling, source = “X_clone”, normalize = True), using CoSpar’s default coupling estimator (the “SW” method). With normalize = True, counts were normalized first within each cell-type cluster and then within each clone per time point. Clones contributed in proportion to their per-cluster cell number rather than being binarised (CoSpar default, ignore_cell_number = False).

Statistical significance was assessed by permutation (cs.tl.pvalue_for_fate_coupling, normalize = True, max_N_simutation = 10,000), in which clone labels were shuffled within each time point so as to preserve the clone-size distribution, yielding one-sided p-values for enriched (pvalue_greater) and depleted (pvalue_less) co-occurrence; 10 permutations provide a p-value tail resolution of approximately 10 . A coupling was reported as significant when pvalue_greater < 0.05 and the cell-type pair was supported by at least three distinct clones spanning both cell types.

Pairs that were significant by p-value but supported by only one or two clones were considered weak, and pairs with no supporting clones, for which the permutation p-value is an artifact at a coupling of approximately zero, were not called. No multiple-testing correction was applied across partitions, as each partition is evaluated against its own permutation null and cell-type groupings are not preserved across annotation resolutions; this is consistent with common practice in clonal lineage-tracing analyses and with the DARLIN application (L. Li et al., 2023).

For partitions containing at least three cell types, the hierarchical organization of coupled fates was computed (cs.tl.fate_hierarchy, source = “X_clone”, normalize = True) and visualized as a dendrogram.

##### Fate bias

###### Observed clonal fate bias

Within each partition (barcode-calling stringency × sampling day × annotation resolution), clones of at least two cells were tested for statistical bias toward each cell type present, restricting the test to cell types labeled by at least three distinct clones. For each clone-fate combination, in-fate enrichment was assessed with CoSpar’s cs.tl.clonal_fate_bias (Wang et al., 2022), a per-clone hypergeometric (Fisher’s exact) test, with Benjamini-Hochberg q-values computed stratified by clone size (multiple_hypothesis_correction = True). The one-sided test was run with alternative = “less”, which under CoSpar’s inverted sign convention corresponds to the enrichment (survival-function) tail. A clone was called fate-biased toward a given cell type when its Q-value was below 0.05, and more than 10 % of its cells resided in that fate (clonal_fraction_in_target_fate > 0.1).

###### Predicted fate bias

Ten progenitor-to-terminal analyses were defined, each specified by its differentiation day(s), annotation resolution, and progenitor (t0) and terminal (t1) cell types, and each was run across the four barcode-calling stringencies. As scRIPT clones are captured within a single day and therefore lack an intra-clonal temporal split, the early-to-late axis required by CoSpar’s multi-time-point inference was defined by cell-type identity, progenitor cell types were treated as the earlier state (t0), and terminal cell types as the later state (t1), following the DARLIN application (L. Li et al., 2023).

For each analysis, a transition map was inferred with cs.tmap.infer_Tmap_from_multitime_clones (clonal_time_points = [“t0”], later_time_point = “t1”, smooth_array = [10, 10, 5], sparsity_threshold = 0.1, intraclone_threshold = 0.2, max_iter_N = 10, epsilon_converge = 0.01), requiring at least 20 t0→t1 bridging clones; analyses below this threshold were skipped. Backward fate maps to each terminal state were then computed in a single batched call (cs.tl.fate_map, map_backward = True), and progenitors were assigned to a committed fate. For analyses with two terminal states, a normalized fate bias was computed (cs.tl.fate_bias, pseudo_count = 0) and progenitors were committed with cs.tl.progenitor (bias_threshold_A = 0.6, bias_threshold_B = 0.4, sum_fate_prob_thresh = 0.2, avoid_target_states = True), retaining cells with a fate bias above 0.6 or below 0.4, excluding the ambiguous intermediate range, and excluding cells already residing in a target fate. For analyses with more than two terminal states, a progenitor was committed to its dominant terminal when that terminal’s normalized fate-map share exceeded that of the runner-up by a margin of 0.2 (equal to 2 × bias_threshold_A - 1) and its total reach probability exceeded 0.2.

Differential expression between committed progenitor groups was computed on the committed t0 progenitors only, using sc.tl.rank_genes_groups (Wolf et al., 2018) (method = “wilcoxon”, use_raw = False) so that the test operated on the log-normalized expression in.X (the.raw slot held raw counts), followed by sc.tl.filter_rank_genes_groups; this yields log fold-changes on the normalized scale. For analyses with two terminal fates, each committed progenitor group was compared directly against the other. Transcription-factor-focused figures were generated by restricting the results to a mouse transcription-factor reference list from the cisTarget resource collection (allTFs_mm.txt, https://resources.aertslab.org/cistarget/tf_lists/allTFs_mm.txt; 1,860 transcription factors) (González-Blas et al., 2023).

#### GEMLI, Gene-expression memory analysis

Memory genes were called per partition (barcode-calling stringency × sampling day × annotation resolution × cell type) with GEMLI’s memory_gene_calling() (Eisele et al., 2024) using per-clone means (use_median = FALSE), lineage sizes 3 to the largest clone (valid_lineage_sizes = 3:N), ground_truth = TRUE, and a fixed seed (42); a gene was called memory when its GEMLI p-value was ≤ 0.05, without per-gene multiple-testing correction (as in the GEMLI workflow). Partitions were analyzed only when ≥ 2 lineages passed a minimum-clone-size filter (size ≥ 3), ≥ 5 lineages had sufficient cells (min_clones = 5), ≥ 10 cells carried valid lineage IDs (min_cells = 10), and ≥ 50 genes passed expression filtering (detection fraction ≥ 0.10). The sweep spanned three stringencies (Q45, Q50, Q55), eleven resolutions (res1-res3, res4a, res4b, res5-res10; ∼5-60 cell types) and three days (D4, D5, D7), run in parallel (parallel::mclapply) (Bengtsson, 2021) with per-worker seeding.

Tiers were built from Q50 calls. For each day and resolution, a resolution-level set was the union of memory genes across cell types; the day-level resolution-robust set was their intersection across all eleven resolutions. Genes were assigned by the number of days robust (n_day): Tier-1, Tier-2, and Tier-3 for one, two, and three days; Tier-0 for genes called memory but robust on no day. Repeating this per stringency, Tier-4 was the intersection of the Q45, Q50, and Q55 Tier-3 sets.

Enrichment of each tier for 107 cataloged mouse imprinted genes (https://www.geneimprint.com/site/genes-by-species.Mus+musculus) was tested against the Q50 GEMLI-tested universe by one-sided Fisher exact test (scipy.stats.fisher_exact, alternative = “greater“; sample odds ratio ad/bc), with Bonferroni correction across the tier tests. Tier-3 robustness was assessed by relaxing the eleven-of-eleven requirement to K-of-eleven (K = 6-11); K = 11 was retained.

##### Differential expression between persistent and non-persistent Naïve-PSC clones

D4/D5/D7 single-cell counts (D457 dataset; 25,562 cells × 27,998 genes after QC) were log-normalized in Seurat v5 (NormalizeData, LogNormalize, scale.factor = 10□); all effect sizes and tests were computed on these natural-log values without reverting to raw counts. Cells were restricted to the naïve pluripotent (Naïve_PSC) compartment at celltype_res8 (300 cells), and clones were assigned from scRIPT scar barcodes at Q50 stringency. Two persistent clones (9 cells at D4, 10 at D7) were compared against thirteen non-persistent within-day reference clones (seven D4, six D7), giving 19 versus 21 cells.

Genes detected (non-zero) in ≥ 10 % of cells on either side were retained (OR-logic, MIN_PCT = 0.10). For each, per-group means, pooled SD (ddof = 1), Cohen’s d ((mean_B - mean_A)/pooled SD; negative d = higher in persistent) and AUROC (normalized Mann-Whitney U, mid-rank ties) were computed on log-normalized values. Significance was assessed by two-sided cell-label permutation of |Cohen’s d| (10,000 permutations, seed = 1; smoothed p = (exceed + 1)/(N + 1), floor ≈ 10□□), Benjamini-Hochberg-adjusted (Benjamini & Hochberg, 1995) across detected genes.

Candidates (BH-FDR < 0.05) were retained only if both persistent clones’ mean shifts, relative to the median of the thirteen non-persistent clone means, agreed in sign. Each was cross-referenced against the Q50 GEMLI memory-gene sweep (374 partitions; tier and recurrence recorded) and the 107-gene cataloged mouse-imprinted set (https://www.geneimprint.com/site/genes-by-species.Mus+musculus). Robustness was checked by leave-one-non-persistent-clone-out (direction stability, effect-size loss) and by an exhaustive clone-level permutation null over all 105 two-versus-thirteen splits (two-sided on |Δ|; floor ≈ 0.019; BH-adjusted). Analyses used R 4.5 and Seurat v5 with a fixed seed = 1.

## Data And Resource Availability

scRNA-seq data generated in this study have been deposited in the BioStudies database under the accession number E-MTAB-17351 (Sarkans et al., 2017). The code used is available at https://github.com/ridvan-cetin/scRIPT_Analysis_Notebooks. Our scRNA-seq can be explored with the Shiny app link at https://ridvan-cetin-scriptlineagetracing.share.connect.posit.cloud. All other relevant data and details of resources can be found within the article and its supplementary information.

## Declaration of generative AI in the writing process

During the preparation of this work, the author(s) used Claude.ai (Opus 4.7, 4.8, 5, and 5.5 models) and Grammarly to improve the clarity of writing. After using this tool/service, the author(s) reviewed and edited the content as needed and took full responsibility for the content of the publication.

## Acknowledgements

We dedicate this article to the memory of our dear colleague, mentor, and senior author, Frank Grosveld, who passed away on June 25, 2026, in the preparation phase of this manuscript. His passion for science and enduring spirit continue to inspire us. Frank was an outstanding, world-class scientist, mentor to 126 PhD students, inspiring colleague, and, most importantly, an amazing human being. R.C. is especially indebted to Frank Grosveld for personally supporting two years of this work from his own savings.

We are grateful to Alex Maas and Tsung Wai Kan for their invaluable assistance with running samples at the FACS facility, to Bart Theeuwes for his tremendous help during the cell-type annotation and his suggestions to improve the manuscript text, and to Buys de Barbanson for his fantastic help during the analysis of raw lineage data processing and cell-to-scar assignment. We thank Sixtine Aymen de Lageard for the help in improving figure clarity and visual harmony, and Fernando Camargo and colleagues for sharing the CARLIN mouse embryonic stem cell line.

## Author contributions

Conceptualization: R.C., J.v.S., E.M., F.G.; Methodology: R.C., E.B.; Formal analysis: R.C.; Investigation: R.C., E.B.; Resources: R.C., G.P., M.A.S., R.H., G.v.B., F.G.; Data curation: R.C.; Writing - original draft: R.C., D.H., J.G., F.G.; Writing - review & editing: R.C., J.v.S., L.K., G.P., M.A.S., R.H., G.v.B., E.M., D.H., J.G., F.G.; Visualization: R.C.; Supervision: F.G., D.H., J.G.; Project administration: F.G., D.H.

## Supplementary Figures

**Fig. S1.**
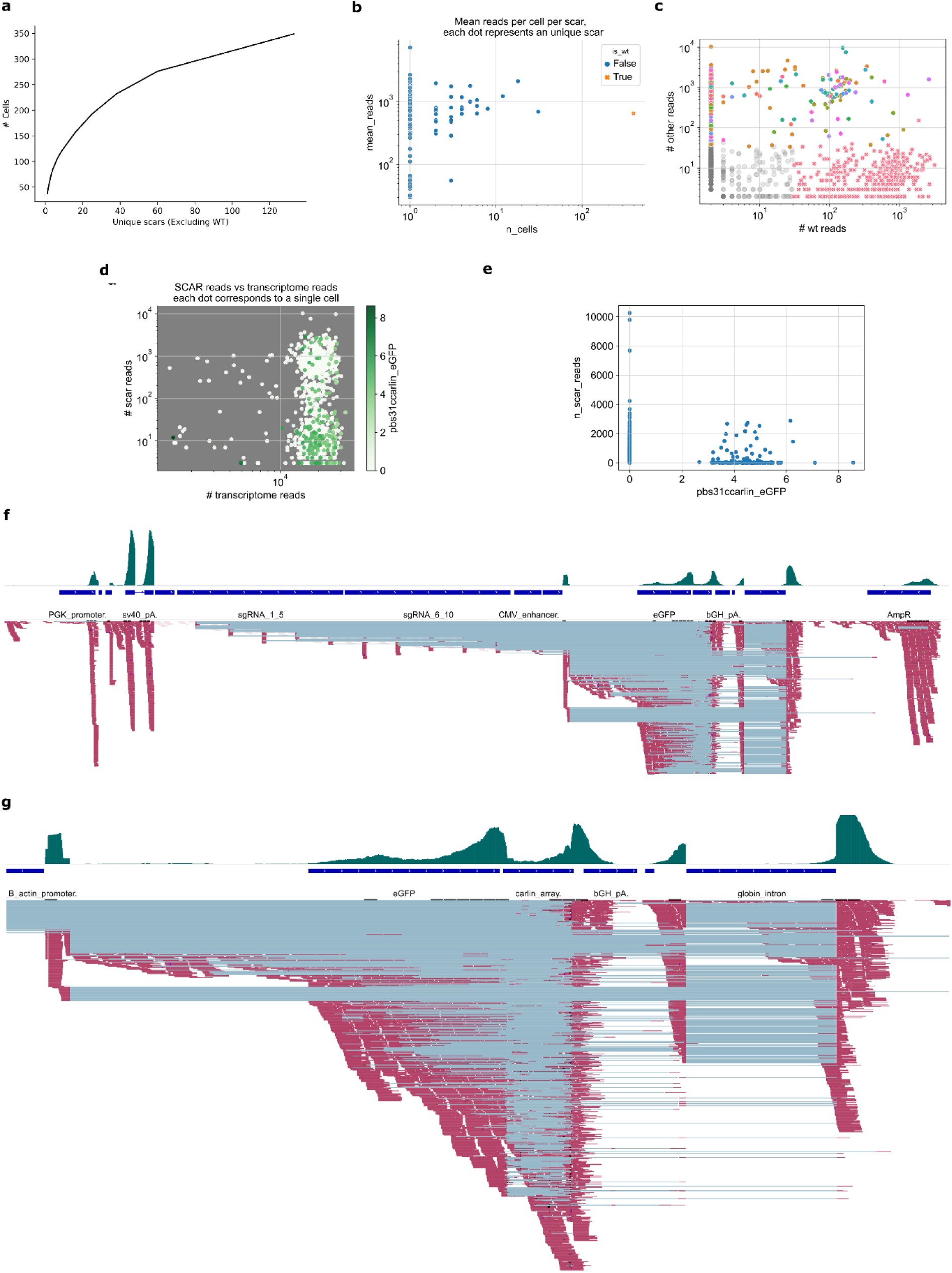
CARLIN test run, demonstrating suboptimal lineage barcode capture efficiencies. **a**, Cumulative distribution of unique lineage barcodes as a line plot from the lineage library of CARLIN-Day4. The plot collectively shows 133 total unique scars identified across all cells from the lineage library using the CARLIN method. **b**, Scatter plot comparing mean reads per scar (y-axis) versus the number of cells harboring each scar (x-axis), both axes on a log scale. Wild-type array (orange cross, True) shows amplification similar to scarred arrays (blue dots, False), indicating there is no substantial amplification bias between wild-type and edited sequences. **c**, Classification of cells based on wild-type versus edited array reads. Red crosses (mainly in the bottom right half of the plot) indicate cells with predominantly wild-type arrays; the colored dots represent cells with confidently assigned scars, while the grey dots (mainly in the bottom left half of the plot) show cells below detection thresholds. **d**, Correlation between lineage library reads (y-axis: number of scars) and transcriptome library reads (x-axis: number of transcriptome reads) for each cell on a log scale. The color gradient (from white to dark green) indicates the CARLIN-eGFP transcript abundance as detected in the transcriptome data, and demonstrates variable transgene expression across cells. **e**, Direct comparison of scar-enriched library reads (y-axis) versus CARLIN-eGFP transcriptome counts (x-axis), showing weak correlation between lineage barcode detection and transgene expression levels. **f**, Genome-wide view of transcriptome read alignment to the CARLIN-*Col1a1* integration site (≈12 kb long) visualized using Integrative Genomics Viewer. Some of the individual sequencing reads (shown as red horizontal bars) display alignment patterns with gaps (light blue-gray segments) representing insertions, deletions, or splice junctions within the transgene array. The bottom track shows annotated genomic features at the *Col1a1* locus (dark blue bars), while the top track displays cumulative read coverage depth (dark green peak areas). **g**, High-resolution view of the CARLIN TA architecture (3,595 bp) (zoomed-in version of panel f) showing detailed transcriptome alignment patterns. Individual reads (red) with alignment gaps (light blue-gray) map to the multi-component transgene cassette. The read coverage depth (green peaks, top) demonstrates a variable expression pattern across the transgene.

**Fig. S2.**
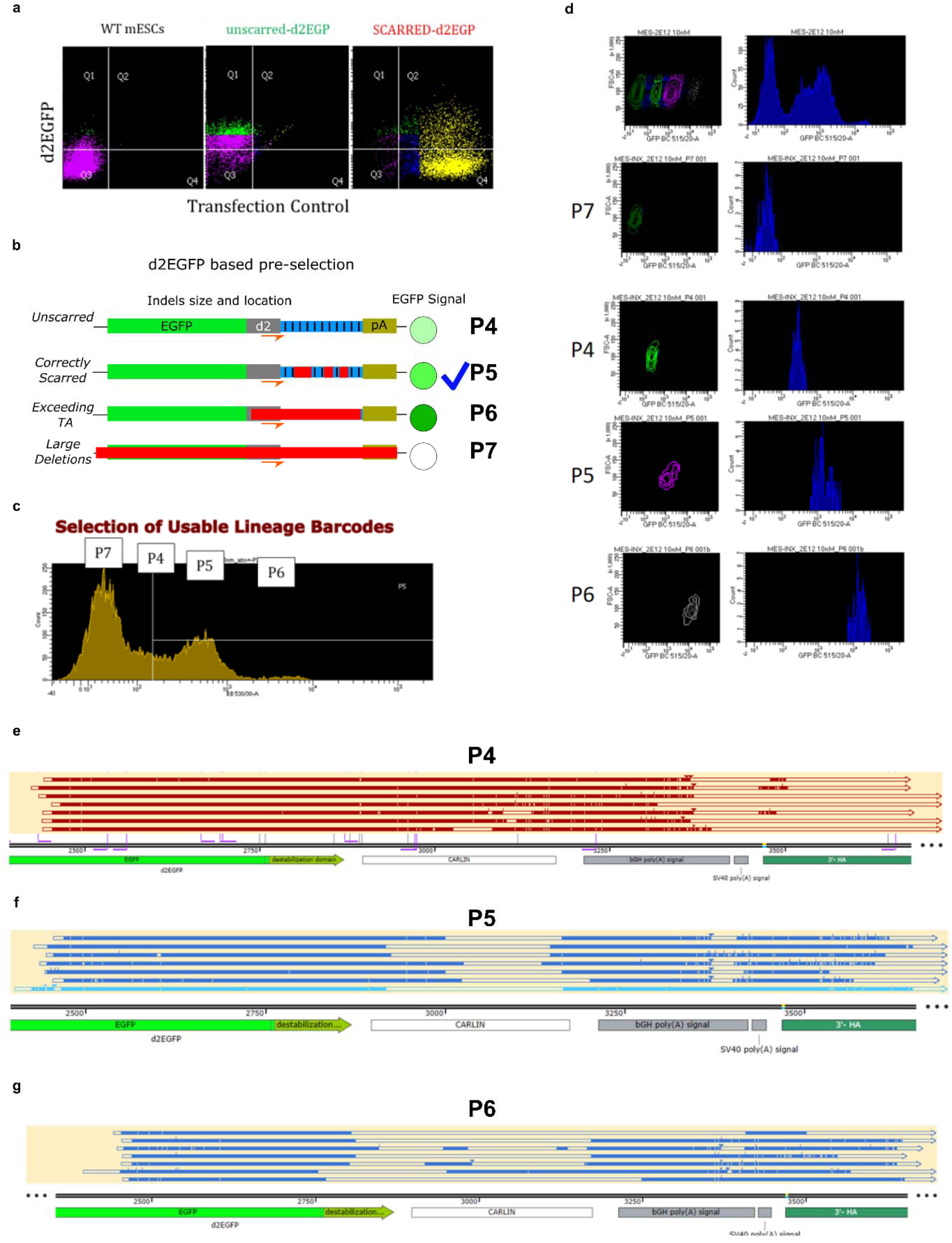
FACS-based enrichment for cells with traceable lineage barcodes (pre-selection). **a**, Representative flow cytometric analysis comparing d2eGFP fluorescence and transfection efficiency across three conditions. Left panel: wild-type (WT) mESCs showing background fluorescence only (purple in Q3). Middle panel: Unscarred scRIPT-mESCs displaying uniform d2eGFP levels (green) without fluorophore (ATTO550) signal. Right panel: scRIPT-mESCs 24-36 hours post-transfection with ATTO550-labeled Cas9-RNP complexes, showing distinct populations, as based on d2eGFP levels. Quadrants Q1-Q4 reveal Q2 (d2eGFP+/ATTO550+) with successfully transfected cells with intact or small indels that maintain d2eGFP signal, and Q4 (d2eGFP-/ATTO550+), indicating transfected cells with large deletions disrupting reporter production. Q3 still contains untransfected cells retaining baseline d2eGFP levels. **b**, Schematic illustration depicting possible outcomes of pre-selection by FACS, considering the relationship between indel patterns and resulting d2eGFP levels (from white to darker green). The thick sign (blue) indicates optimal scarring, preserving the binding sites for the primers. P4: the intact transgene with baseline eGFP signal; P5: partial deletions that increase d2eGFP stability; P6: extensive deletions exceeding the TA and affecting the destabilization domain; P7: example of complete transgene deletion, which eliminates fluorescence (white). **c**, Distribution of d2eGFP fluorescence intensity about 30 hours post-RNP transfection, showing four distinct populations: P7 (d2eGFP-negative, large deletions; see also panel b), P4 (baseline d2eGFP, mostly-WT array), P5 (moderate d2eGFP increase, optimal partial indels), and P6 (high d2eGFP, destabilization domain disrupted, i.e., the lineage primer binding site had been deleted). **d**, Flow cytometry plots showing the isolation strategy for the entire population and the four subpopulations of cells (P7, P4, P5, P6). The left panels display FSC-A versus d2eGFP intensity; the right panels show d2eGFP peaks with population-specific gates. **e-g**, SnapGene visualization of sequenced lineage barcodes from representative monoclonal colonies. P4 colonies retain wild-type array structure (shown as uniform red bars).

**Fig. S3.**
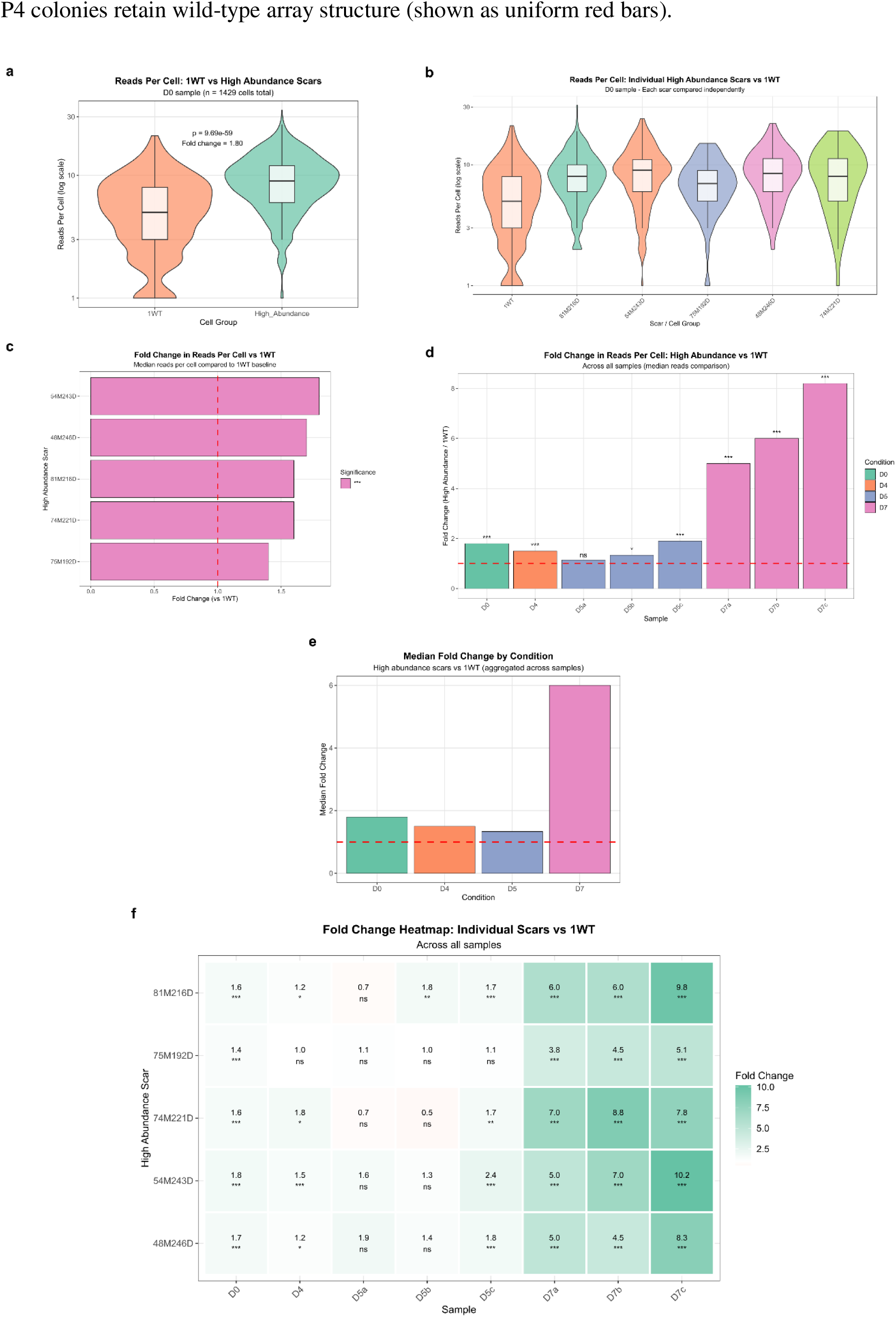
Indels in the transgene increased mRNA abundance at the cellular level. **a**, Violin plots comparing lineage library reads per cell between wild-type (1WT) and high-abundance scar populations from the D0 sample (n=1,429 cells). Wild-type cells (n=719, orange) show baseline expression, while cells with high-abundance scars (n=710, green) demonstrate 1.8-fold higher median reads per cell (p=9.69e-59, Wilcoxon rank-sum test). Box plots show median, quartiles, and 1.5× IQR. **b**, Individual comparisons of reads per cell for each high-abundance scar (for 81M216D, 54M243D, 75M192D, 48M246D, 74M221D, as indicated on the x-axis) versus 1WT baseline. Each scar independently shows variably increased transgene expression relative to WT. **c**, Fold-change in median reads per cell for five (see panel b) individual high-abundance scars compared to WT (1WT). Red dashed line marks fold-change=1.0 (i.e., no difference). **d**, Sample-level analysis showing fold-change between high-abundance and 1WT cells across all experimental time points. Bars colored by condition (D0=green, D4=orange, D5=blue, D7=pink). Statistical significance is indicated above the bars (***p<0.001, **p<0.01, *p<0.05, ns=not significant). **e**, Condition-level median fold-changes aggregated across replicate samples at each time point. The red dashed line marks the baseline. **f**, Heatmap displaying scar-specific fold-changes across all samples. Color intensity indicates fold-change magnitude (orange <1.0, white =1.0, green >1.0); numerical values and significance levels (including ns, non-significant) are shown.

**Fig. S4.**
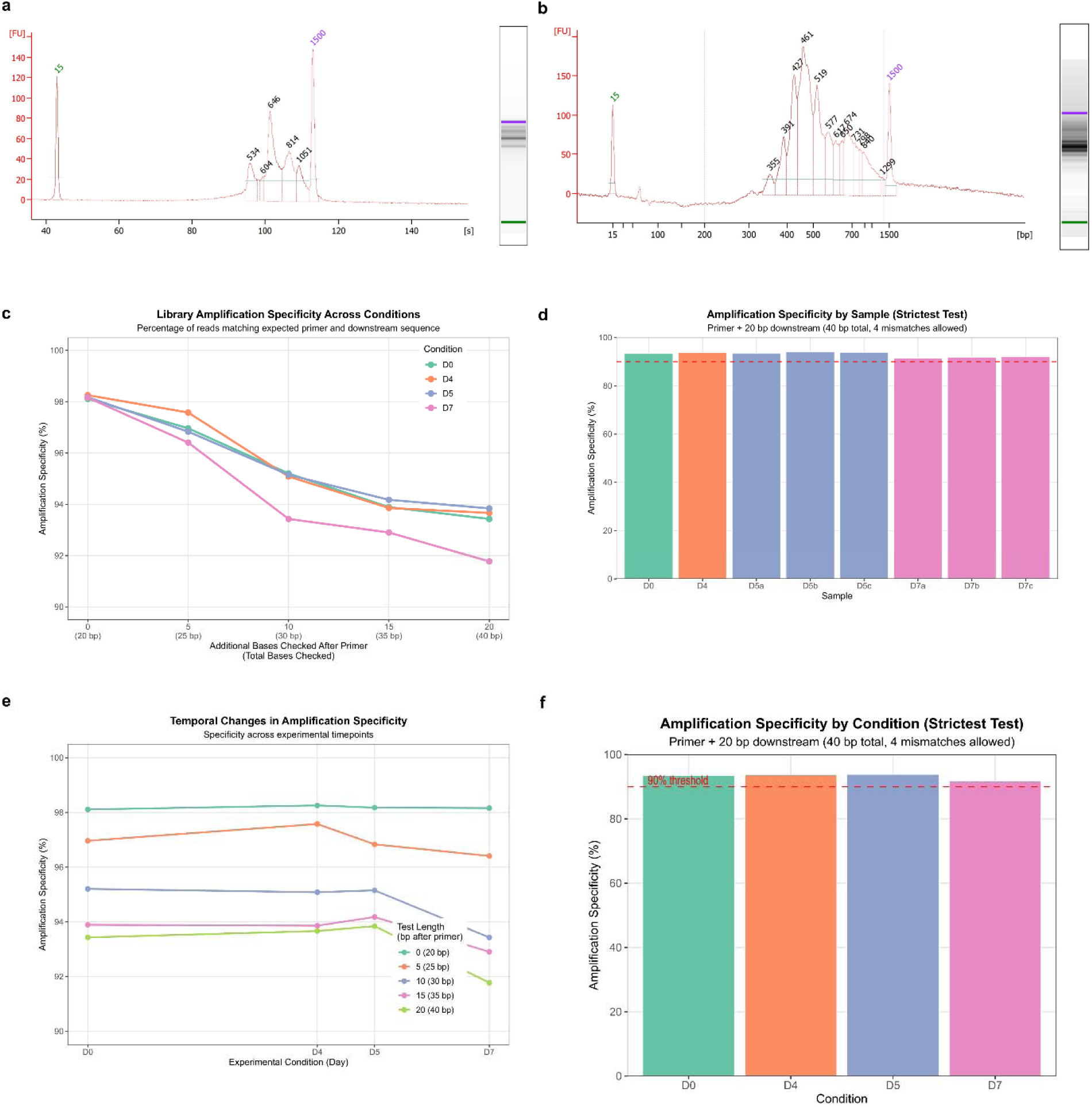
Validation of the applied simplified lineage library preparation and amplification specificity across conditions. **a**, Representative Bioanalyzer electropherogram trace displaying the size distribution of the PCR-amplified lineage library. The major peak at 646 bp represents the primary amplicon containing scarred and wild-type target arrays. Other peaks, given here at 534, 604, 814, and 1051, represent transcripts with different lengths of their 3’UTR. The peaks at 15 bp and 1500 bp represent internal standards. **b**, Representative electropherogram of a final indexed library ready for Illumina sequencing. The dominant peak area between 400 and 600 bp confirms successful library preparation. **c**, Line plot showing the percentage (between 90 and 100) of reads matching the expected primer and downstream sequences across five different nucleotide distances that are present in TA. Each line represents one experimental time condition (D0, D4, D5, D7), with points indicating specificity at each test length (from 20 to 40 bp). The x-axis shows progressive additional bases (0, 5, 10, 15, 20 bp) checked beyond the 20-bp primer, with the total checked bases in parentheses. Mismatch tolerance set at 1 per 10 bp (2-4 mismatches allowed). Data derived from 50.8 million (M) PE150 reverse reads across 8 samples (D0=5.8M, D4=6.1M, D5=19.6M, D7=19.3M). All conditions maintain >90% specificity, with D5 showing the highest (93.84%) and D7 the lowest (91.78%) specificity at the strictest stringency. **d**, Bar chart displaying amplification specificity for individual samples using the strictest test (primer + 20 bp downstream, i.e., 40 bp in total, and 4 mismatches allowed). Bars are colored by experimental condition. The red dashed line marks the 90% specificity threshold. All samples exceed 90% specificity, ranging from 91.53% (D7a) to 94.10% (D5b). **e**, Temporal changes in amplification specificity across experimental time points for all five applied test stringencies. Each line represents a different downstream sequence length (0-20 bp after the primer). **f**, Bar chart displaying amplification specificity for individual conditions (similar to panel d).

**Fig. S5.**
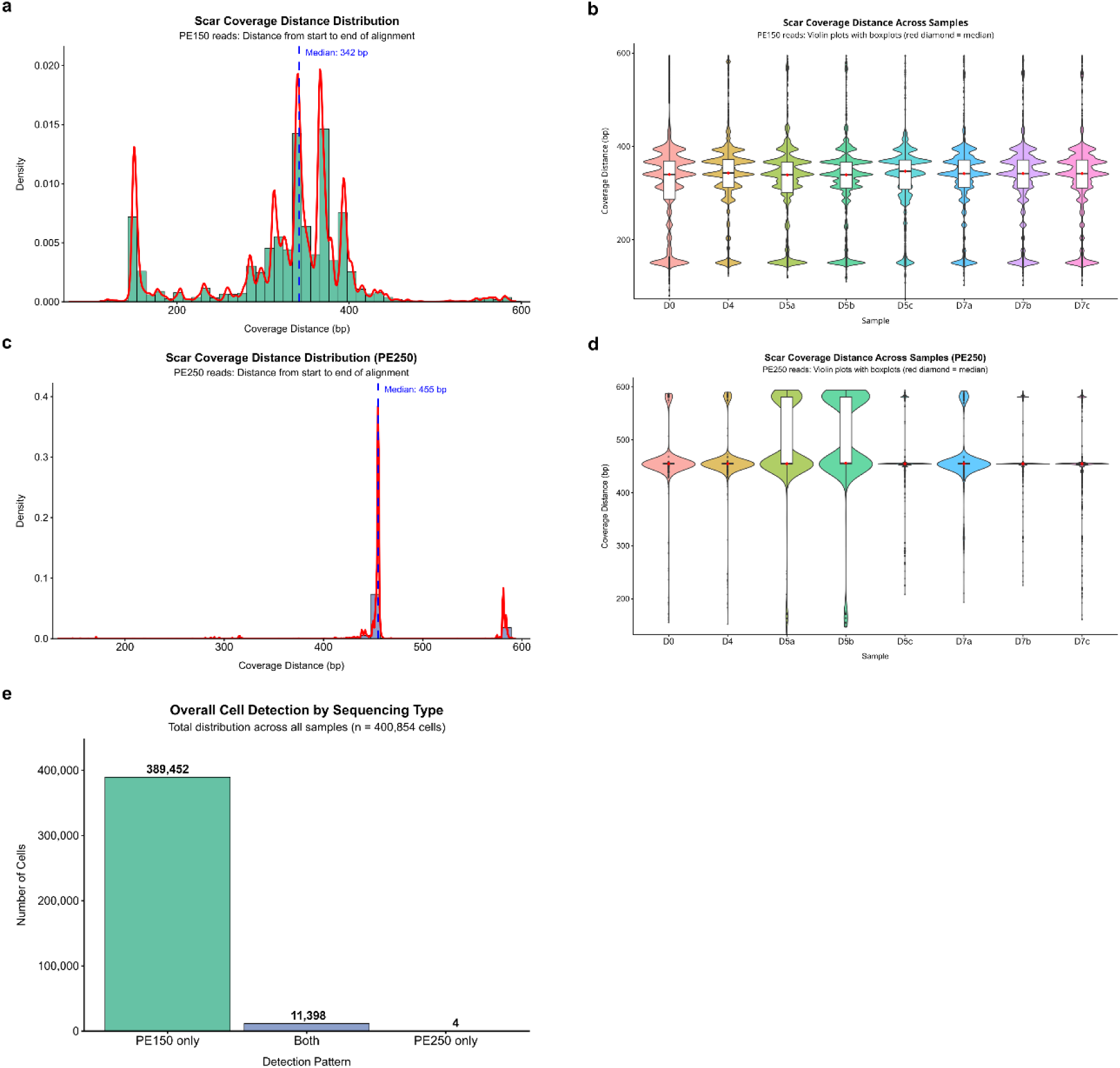
Comparison of PE150 and PE250 sequencing coverage for the detection of lineage barcodes. **a**, Distribution of scar coverage distances for PE150 reads displayed as a histogram (green) with kernel density overlay (red curve). Coverage distance represents the nucleotide span from primer start to alignment end. The blue dashed line indicates the median (342 bp). Data derived from all PE150 reads with valid cell barcodes and scar assignments. **b**, Sample-level scar coverage distributions for PE150 reads are shown as violin plots with boxplot overlays (the red diamond/point marks the median of each sample). Samples show consistent distributions across samples. **c**, Distribution of scar coverage distances for PE250 reads (similar to panel a). The median coverage (455 bp, blue dashed line) extends about 113 bp beyond that of PE150, confirming the advantage of longer reads. Note that PE250 sequencing comprises only about 1.5% of the total sequencing data. **d**, Condition-level aggregation of PE250 scar coverage distances (similar to panel b). **e,** Total distribution of cells by detection pattern across all samples (n=400,854 cells), with absolute counts indicated. PE150 only: 389,452 cells (97.2%); both PE150 and PE250: 11,398 cells (2.8%); PE250 only: 4 cells (<0.001%). This quantitative summary demonstrates that PE150 alone captures nearly all cells with lineage information, with minimal overlap between sequencing runs (2.8%) and essentially no contribution from PE250-only sequencing.

**Fig. S6.**
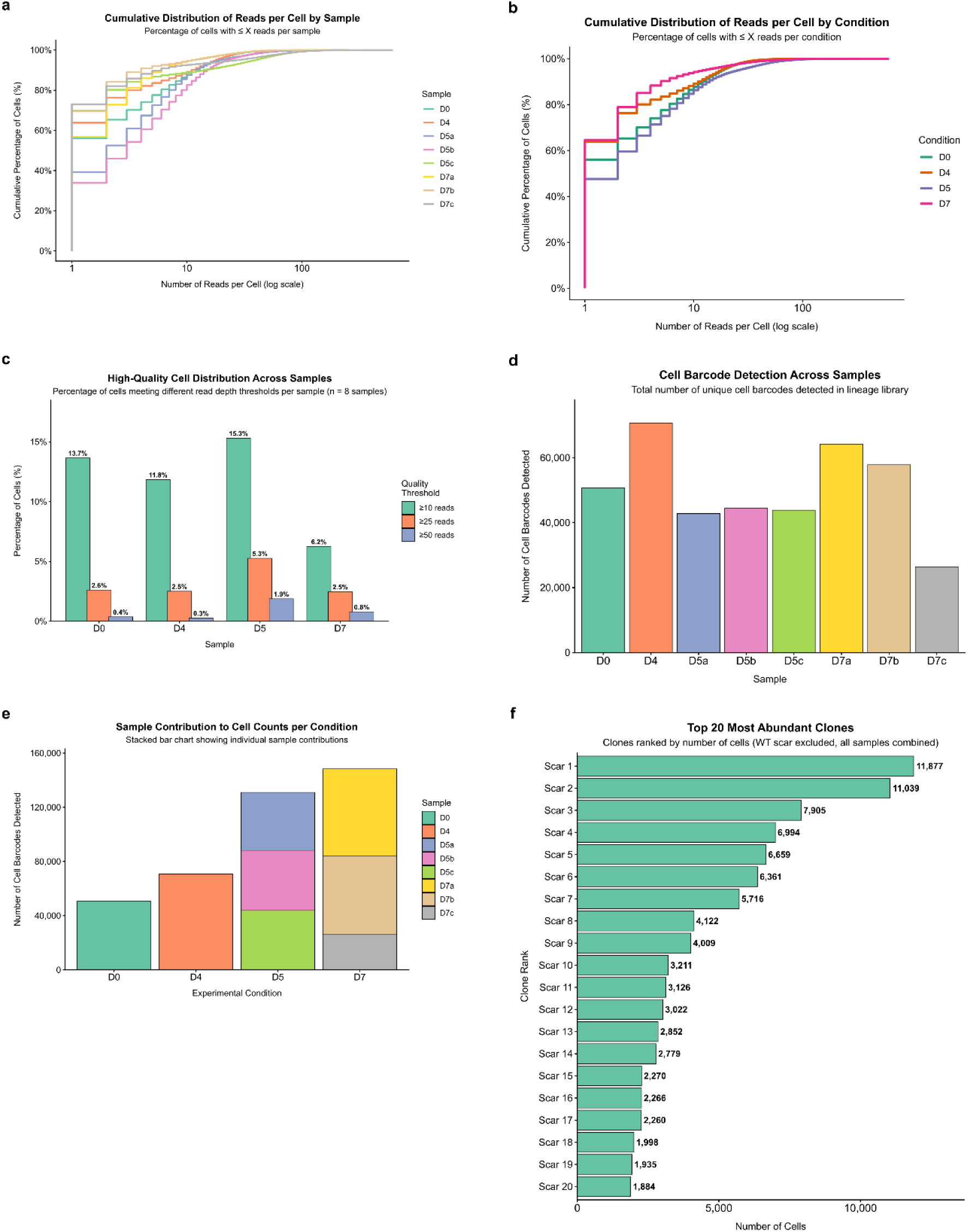
Comprehensive analysis of lineage library sequencing depth per cell, cell barcode detection across samples/conditions, and top-20 most abundant clones from the lineage library. **a**, Cumulative distribution (0-100 %) curves per sample (different time points; different samples at a given time point) on a log scale. **b**, Cumulative distributions (0-100 %) of reads per cell by condition at the four indicated time points on the log scale. **c**, Percentage of cells meeting pre-set quality thresholds per sample. At ≥10 reads threshold (green), 5.9-19.9% pass was obtained; at ≥25 reads (orange), 2.2-5.3% pass; at ≥50 reads (blue-gray), 0.5-2.3% pass. D5 (i.e., D5b) showed the highest retention at all thresholds. These metrics establish that, while most cells have low coverage, sufficient high-quality cells remain for robust lineage analysis after appropriate filtering. **d**, Total number of unique cell barcodes detected in the lineage library across eight experimental samples spread over different time points. Cell barcodes were identified from combined PE150 and PE250 sequencing reads, requiring at least one read with valid scar information. Raw detection counts are shown without quality filtering. Samples represent biological time points (D0, D4, D5, D7), with letters denoting technical partitions for D5 and D7. Total cells detected: n=400,854. **e**, A stacked bar chart showing individual sample contributions to total cell counts per experimental condition. D0 and D4 represent single samples, while D5 and D7 comprise three technical partitions each (a-c). D5 samples show relatively balanced contributions (D5a: 44,080; D5b: 45,556; D5c: 40,401 cells), while D7 samples exhibit more variation (D7a: 63,934; D7b: 58,790; D7c: 25,693 cells). **f**, Top-20 most abundant scarred clones across all samples (WT excluded).

**Fig. S7.**
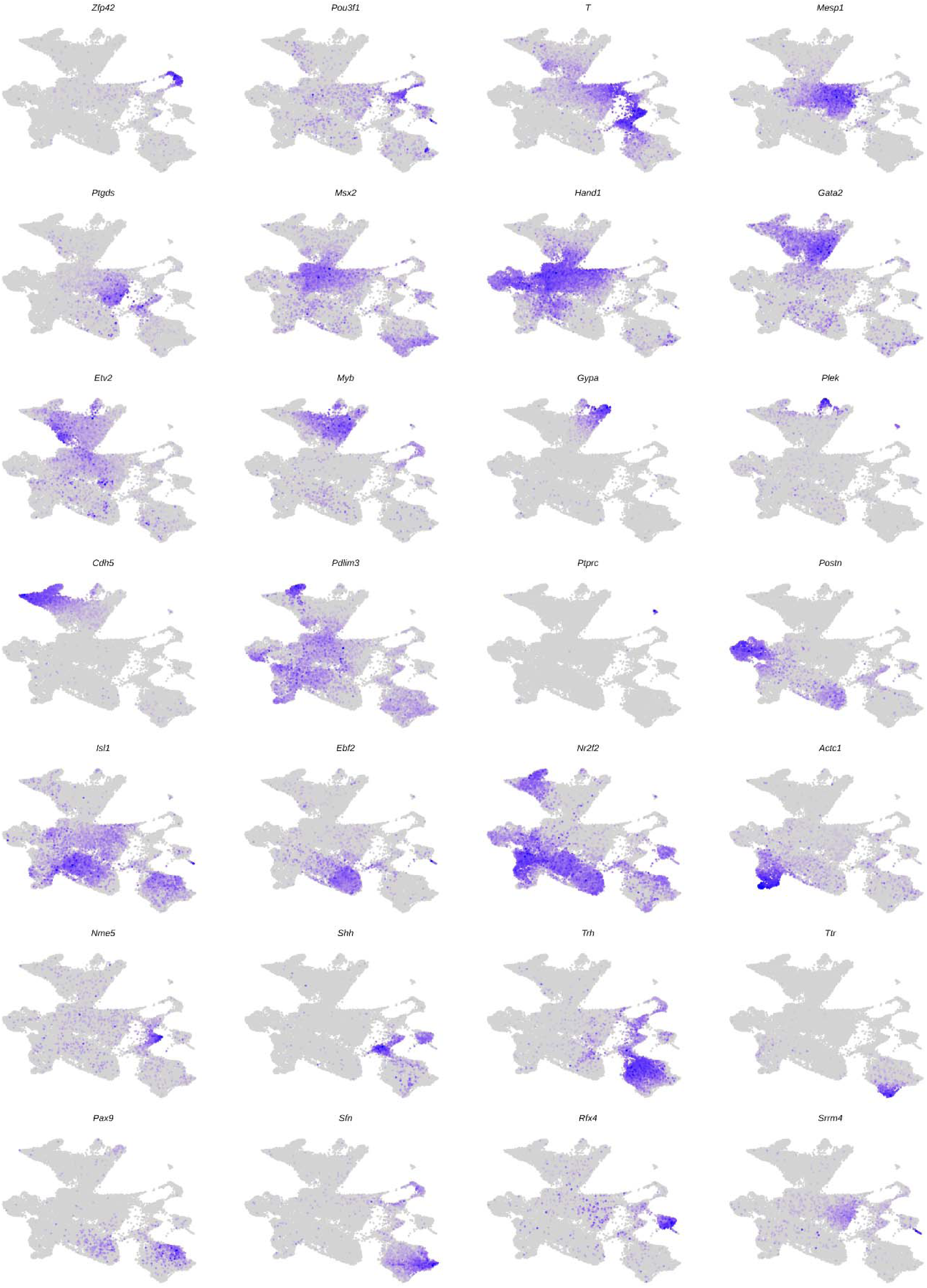
Marker gene expression across the whole embryoid-body (D4-D7) single-cell landscape. Harmony-integrated UMAP projection of 25,562 cells pooled from differentiation days D4, D5, and D7, with each panel colored by the log-normalized expression of one of the 28 marker genes selected (27 of which are listed in Fig. 2b) to span the major cell types present in the merged embryoid body dataset. Cells are plotted in order of increasing expression so that positive cells are drawn on top (order = TRUE); point size 0.5, expression shown on the default low-to-high color scale (grey to blue). Axes and per-panel color bars are omitted for clarity.

**Fig. S8.**
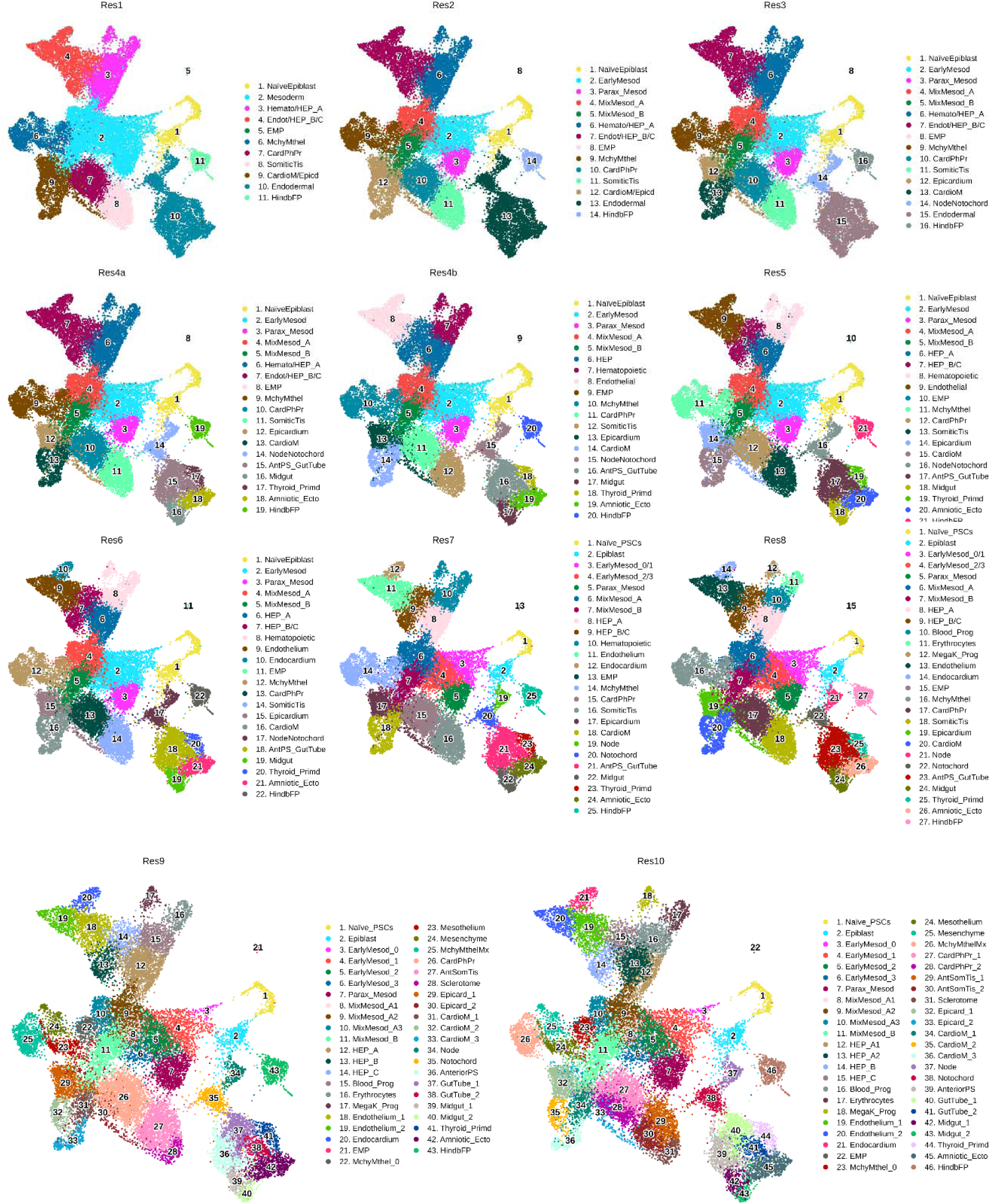
Hierarchical cell-type annotation of the embryoid-body dataset across successive resolutions. Harmony-integrated UMAP projection of 25,562 cells pooled from differentiation days D4, D5 and D7, shown at eleven successive levels of annotation resolution (labelled res1, res2, res3, res4a, res4b, res5, res6, res7, res8, res9 and res10). In each panel, cells are colored by their cell type assignment at that resolution, and each cell type is marked at its median UMAP position with a number; the accompanying legend lists the corresponding number-to-cell-type mapping. Numbers are assigned independently within each panel and therefore do not denote the same cell type across resolutions. Axes are omitted for clarity.

**Fig. S9.**
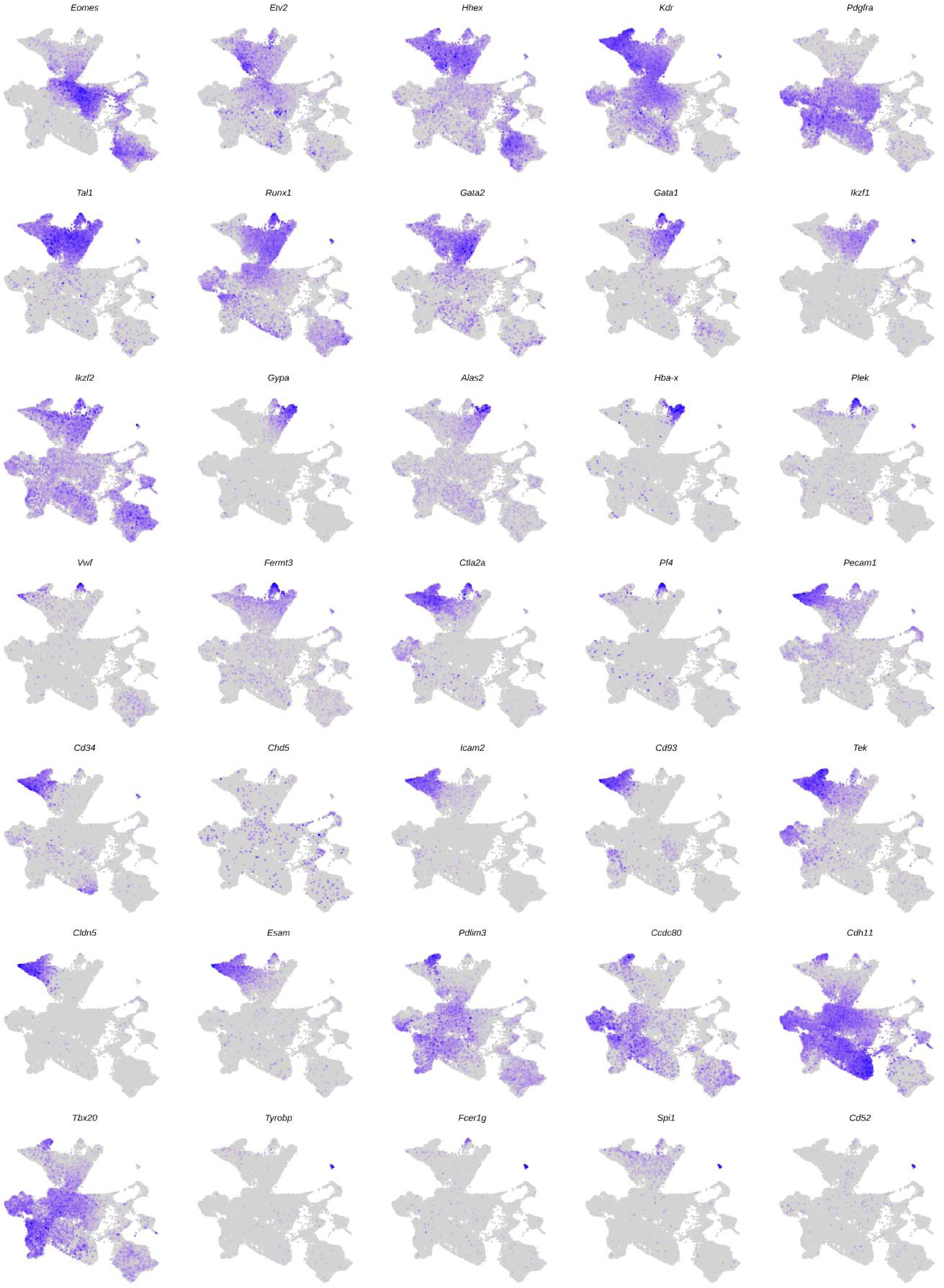
Expression of hematoendothelial lineage markers across the integrated single-cell landscape. Harmony-integrated UMAP projection of 25,562 cells pooled from differentiation days D4, D5, and D7, with each panel colored by the log-normalized expression of one of 35 marker genes. Cells are plotted in order of increasing expression so that positive cells are drawn on top (order = TRUE); point size is 0.5, with expression shown on the default low-to-high color scale (grey to blue). Axes and per-panel color bars are omitted for clarity. Genes are arranged to follow successive hematoendothelial programs.

**Fig. S10.**
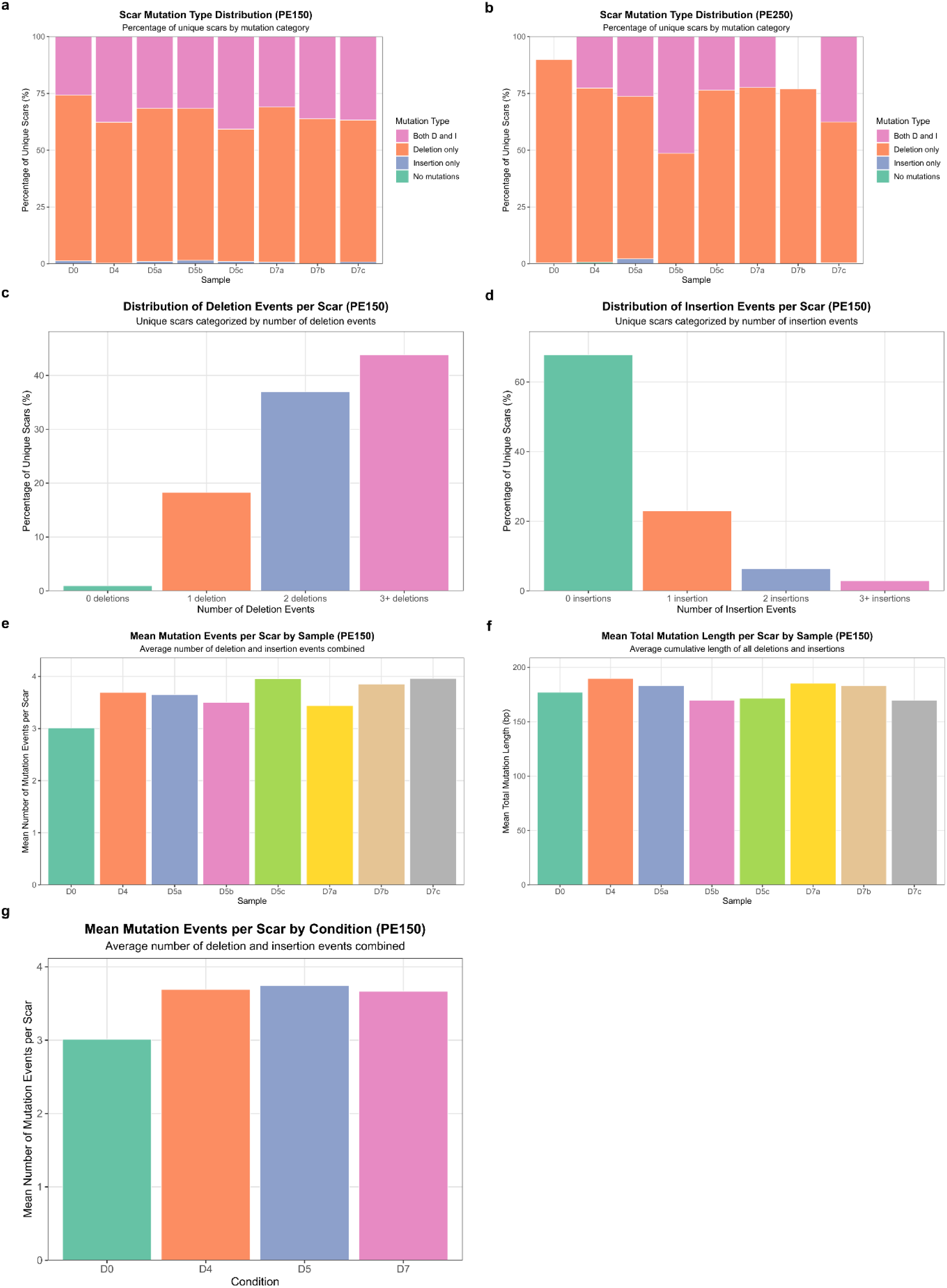
Characterization of lineage barcode mutation patterns. **a**, Percentage distribution of unique scars by mutation type across samples using PE150 data (n=27,962 unique scars). Categories: no mutations (teal, <1%), deletion-only (orange, 60-73%), insertion-only (blue, <1%), both deletions and insertions (pink, 27-40%). Deletion-only scars predominate across all samples. **b**, Mutation type distribution for PE250 data (n=1,727 unique scars) showing similar patterns to PE150, with deletion-only scars comprising 62-90% and combined deletion/insertion scars 10-38%. **c**, Distribution of deletion event frequencies per scar (PE150). Multiple deletion events indicate complex repair outcomes from simultaneous Cas9 cutting at multiple target sites. **d**, Distribution of insertion event frequencies per scar (PE150). **e**, Mean mutation events (deletions + insertions) per scar across samples. **f**, Average cumulative mutation length per scar across samples (170-190 bp). **g**, Condition-level mean mutation events.

**Fig. S11.**
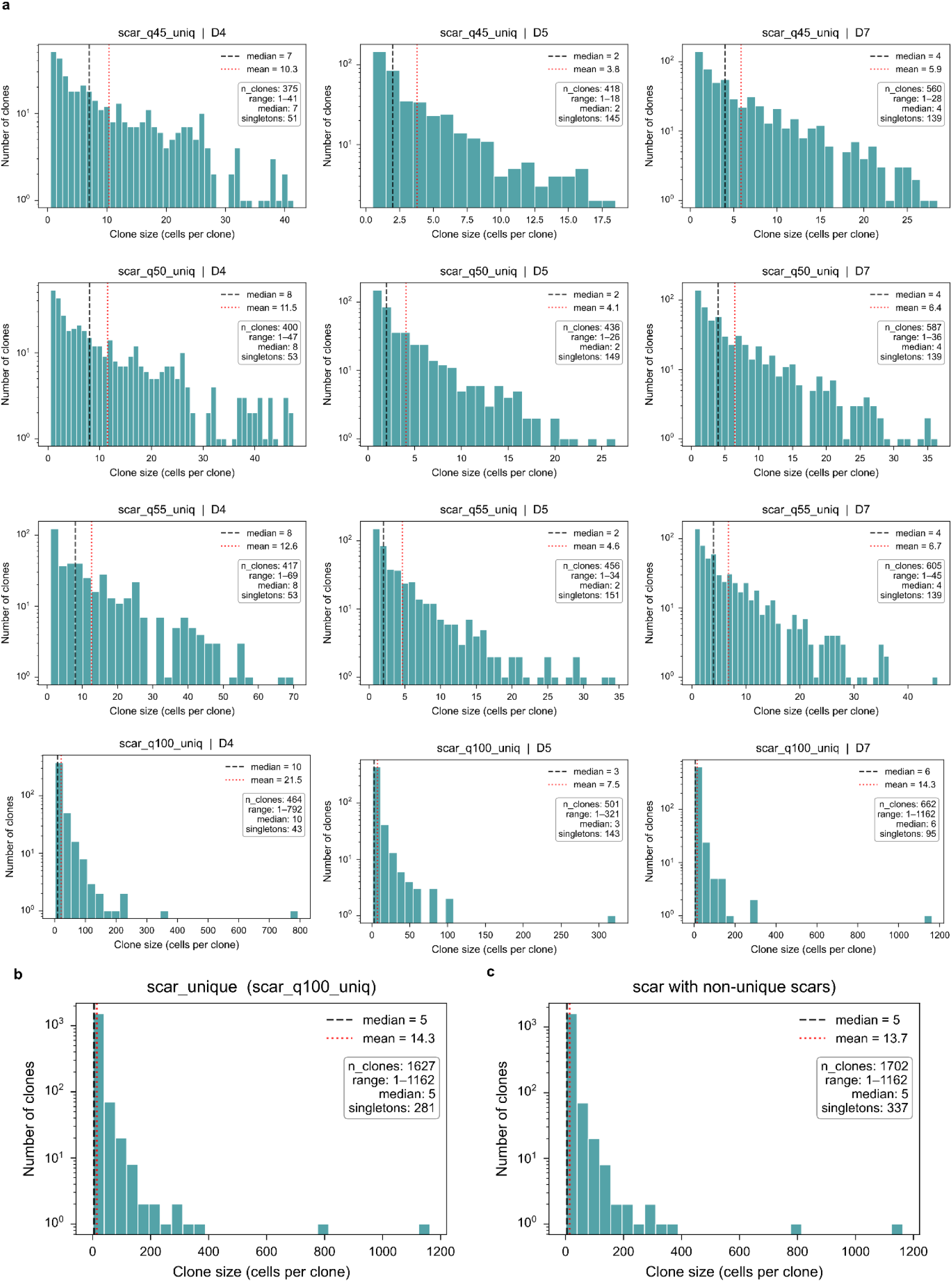
Clone-size distributions across scar stringencies and differentiation days, plus pooled unique-scar and all-scar baselines. **a**, Per-partition clone-size histograms (a 4×3 grid = 12 panels). Rows = scar stringency (from top to bottom: Q45, Q50, Q55, Q100; scar_q{45,50,55,100}_uniq metadata column). Columns = differentiation day (D4, D5, D7). Each panel shows the distribution of clone sizes for the cells assigned a clone under that (stringency, day) partition: x-axis (linear) = clone size in cells; y-axis (log) = number of clones. Teal bars are histograms on integer bins for sizes ≤ 50 (bin width = 1 cell) and on 30 linear bins otherwise. Black dashed line = median; red dotted line = mean. Inset text box: n_clones, size range (min - max), median, and singleton count. **b**, Pooled clone-size histogram for the unique-scar cell set, cells whose scar-barcode uniquely identifies a clone under Q100 (the least-restrictive filter), all days pooled. Same axes, bins, marker lines, and annotation box as in panel a. **c**, Pooled clone-size histogram for the all-scar cell set, cells with any scar-barcode assignment, including barcodes that fail the uniqueness filter (shared across clones), all days pooled. Same conventions as in panel a.

**Fig. S12.**
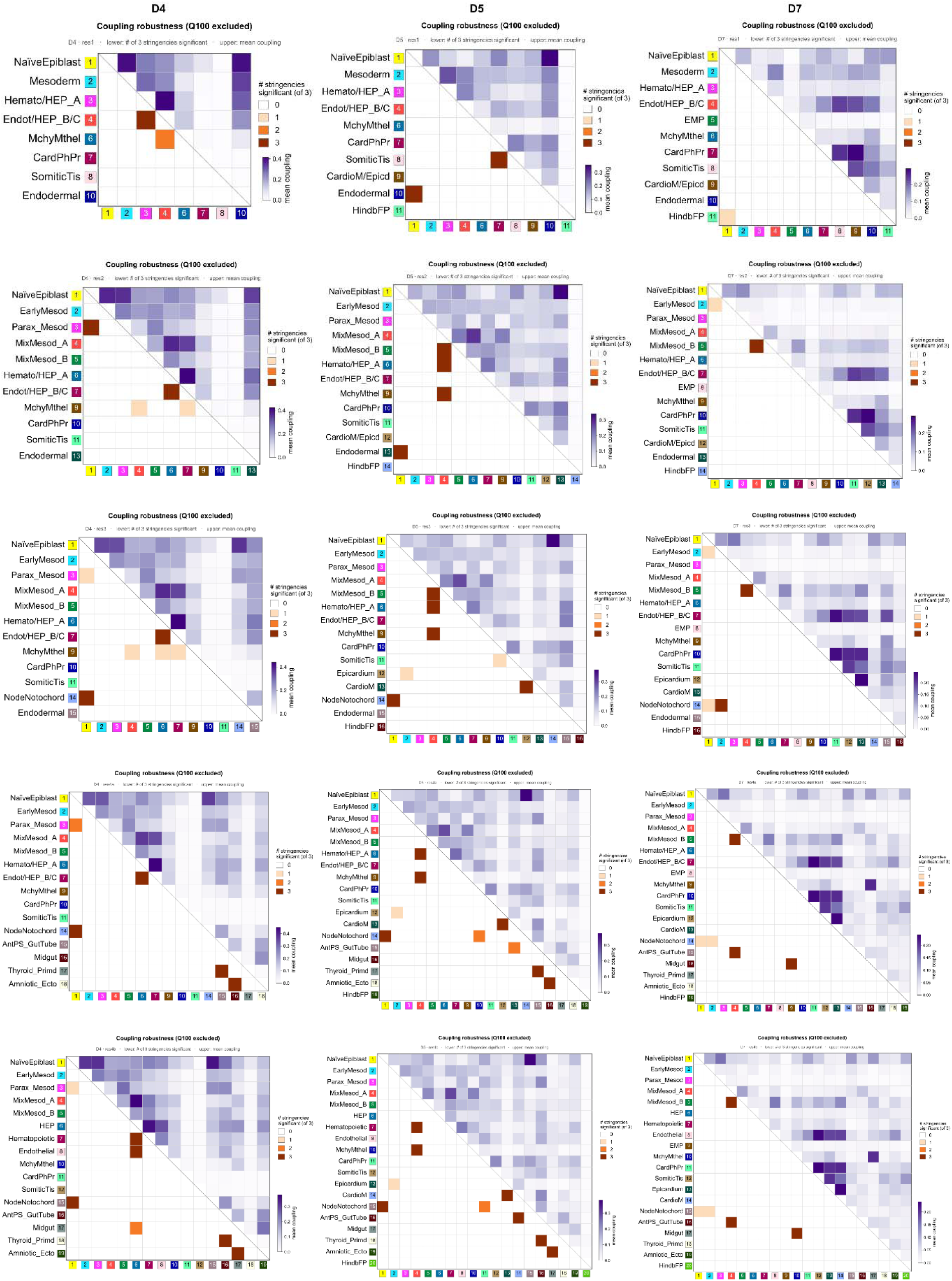
Coupling-robustness heatmaps across differentiation days D4 / D5 / D7 at low-to-medium cell-type annotation resolutions (res1 - res4b). Grid of 5 rows × 3 columns = 15 coupling-robustness heatmaps computed under three project-grade scar stringencies Q45 / Q50 / Q55 (Q100 was excluded). Each orange panel shows how many times a cell-type coupling was significant across three fate-coupling analyses with different stringencies. Rows = cell-type annotation resolution (from top to bottom: res1, res2, res3, res4a, res4b). Columns = differentiation day (D4, D5, D7). In each panel: lower-left triangle = number of the 3 stringencies at which each cell-type pair is significantly coupled (white to dark orange, 0 → 3); upper-right triangle = mean coupling score across the 3 stringencies (white to dark purple, 0 → 0.4).

**Fig. S13.**
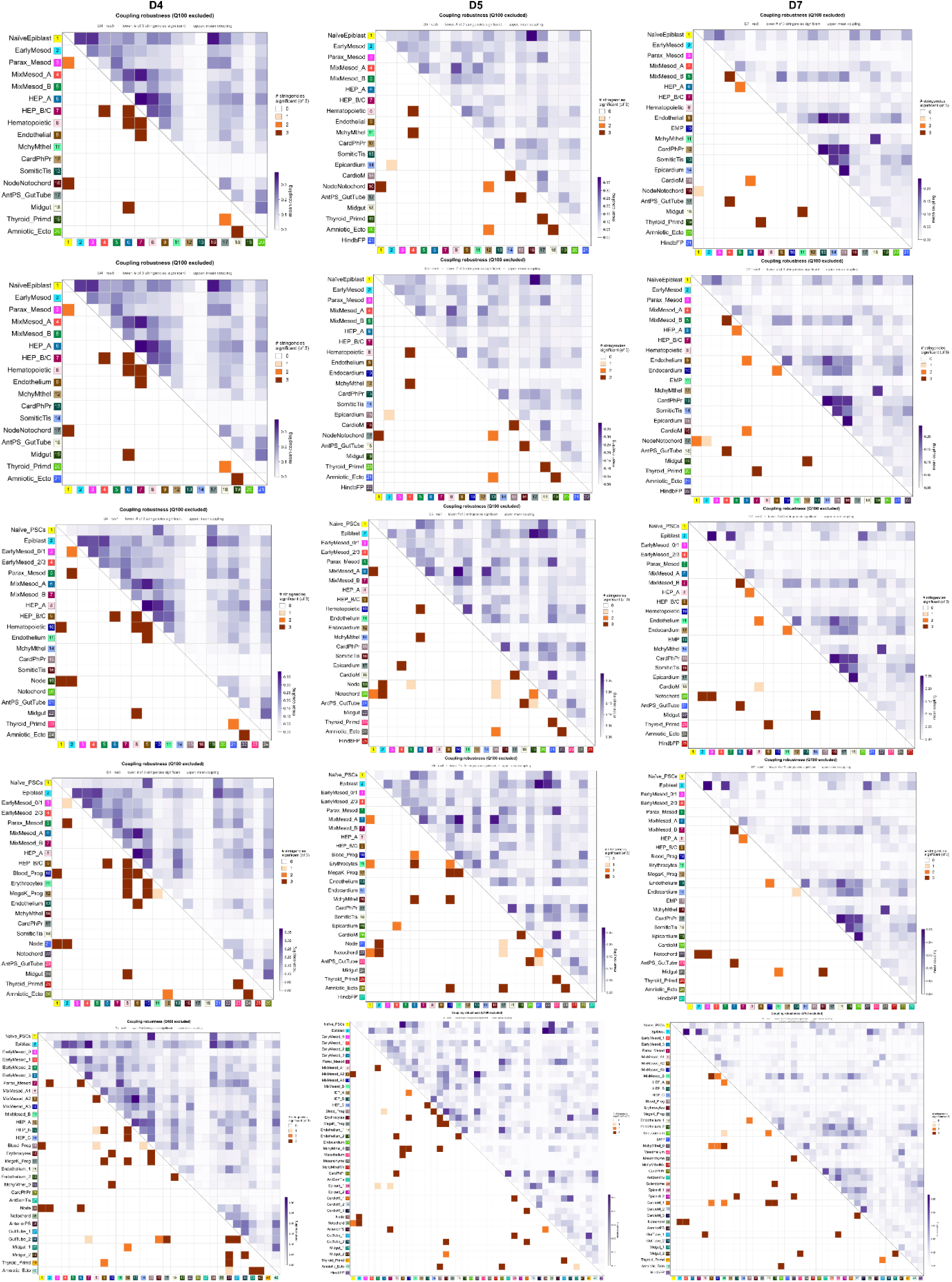
Coupling-robustness heatmaps across differentiation days D4 / D5 / D7 at high cell-type annotation resolutions (res5 - res9). As in Fig. S12, but with rows = res5, res6, res7, res8, res9 (from top to bottom). Same 5 × 3 grid, columns, triangle conventions, and color maps.

**Fig. S14.**
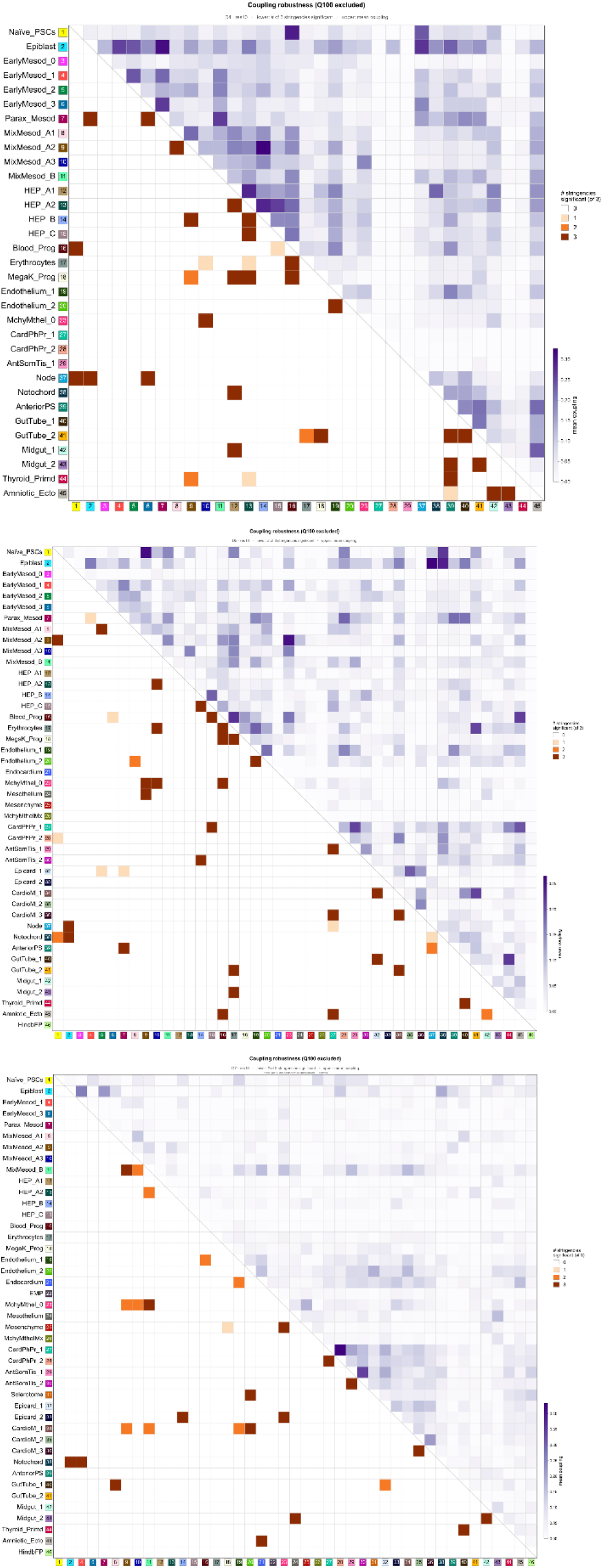
Coupling-robustness heatmaps at the finest cell-type annotation resolution (res10), stacked vertically across days D4 / D5 / D7. Three coupling-robustness heatmaps at res10, arranged vertically (from top to bottom: D4, D5, D7). Same triangle conventions and color maps as in Fig. S12.

**Fig. S15.**
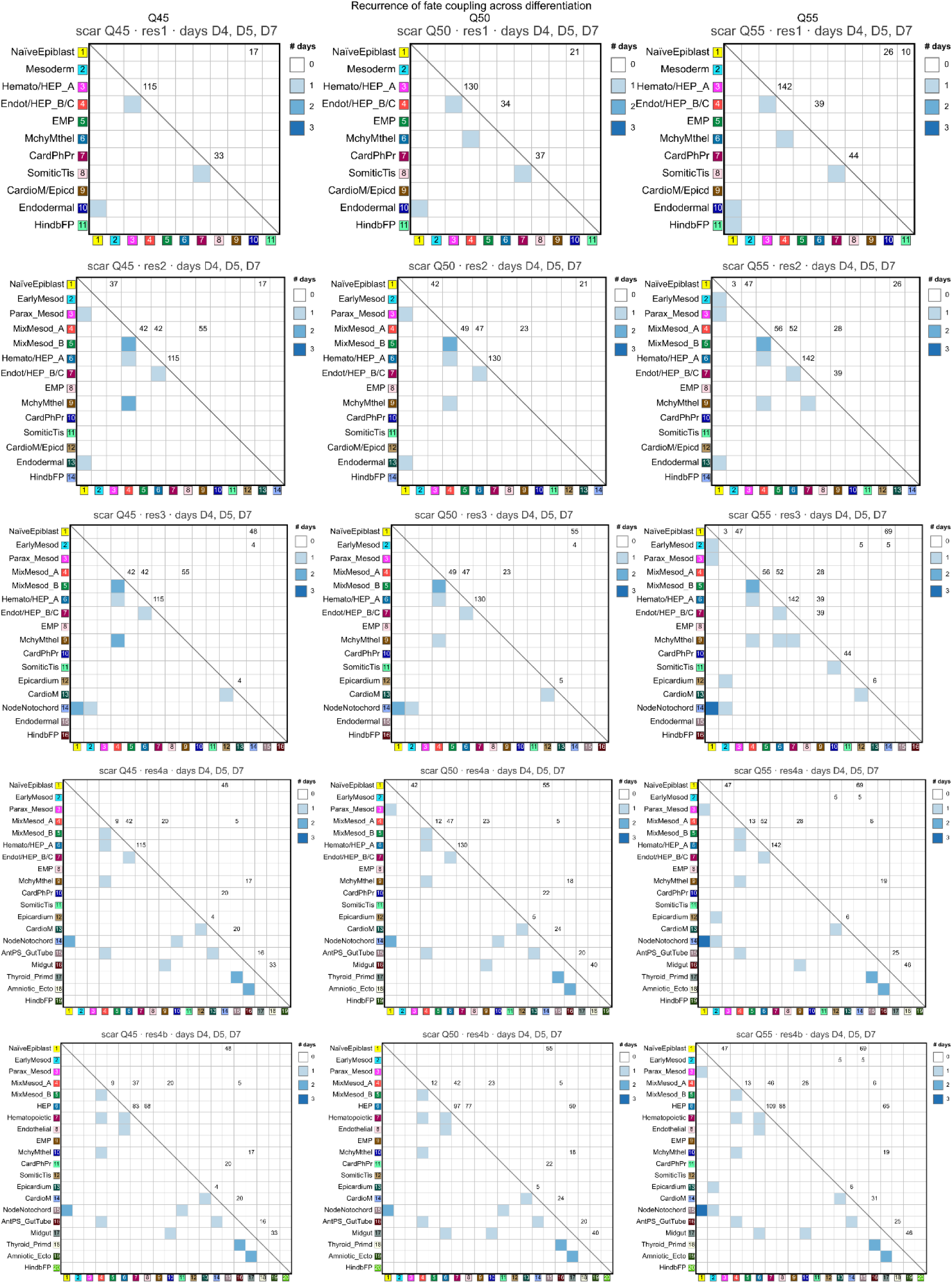
Cross-day recurrence of fate couplings across scar stringencies at low-to-medium cell-type annotation resolutions (res1 - res4b). Grid of 5 rows × 3 columns = 15 heatmaps summarizing the recurrence of significant cell-type couplings across differentiation days D4, D5, D7 within each (stringency, resolution) partition. Rows = cell-type annotation resolution (from top to bottom: res1, res2, res3, res4a, res4b). Columns = scar stringency (Q45, Q50, Q55). In each panel, each cell shows the number of days (0 - 3) at which the corresponding cell-type pair is significantly coupled at that stringency (white → dark blue). When a coupling is present at ≥ 1 day, the maximum supporting clone count across those days is annotated in the cell. Cell types are indexed by numbered, color-tagged labels shared with the corresponding day-resolved heatmaps in Fig. 4.

**Fig. S16.**
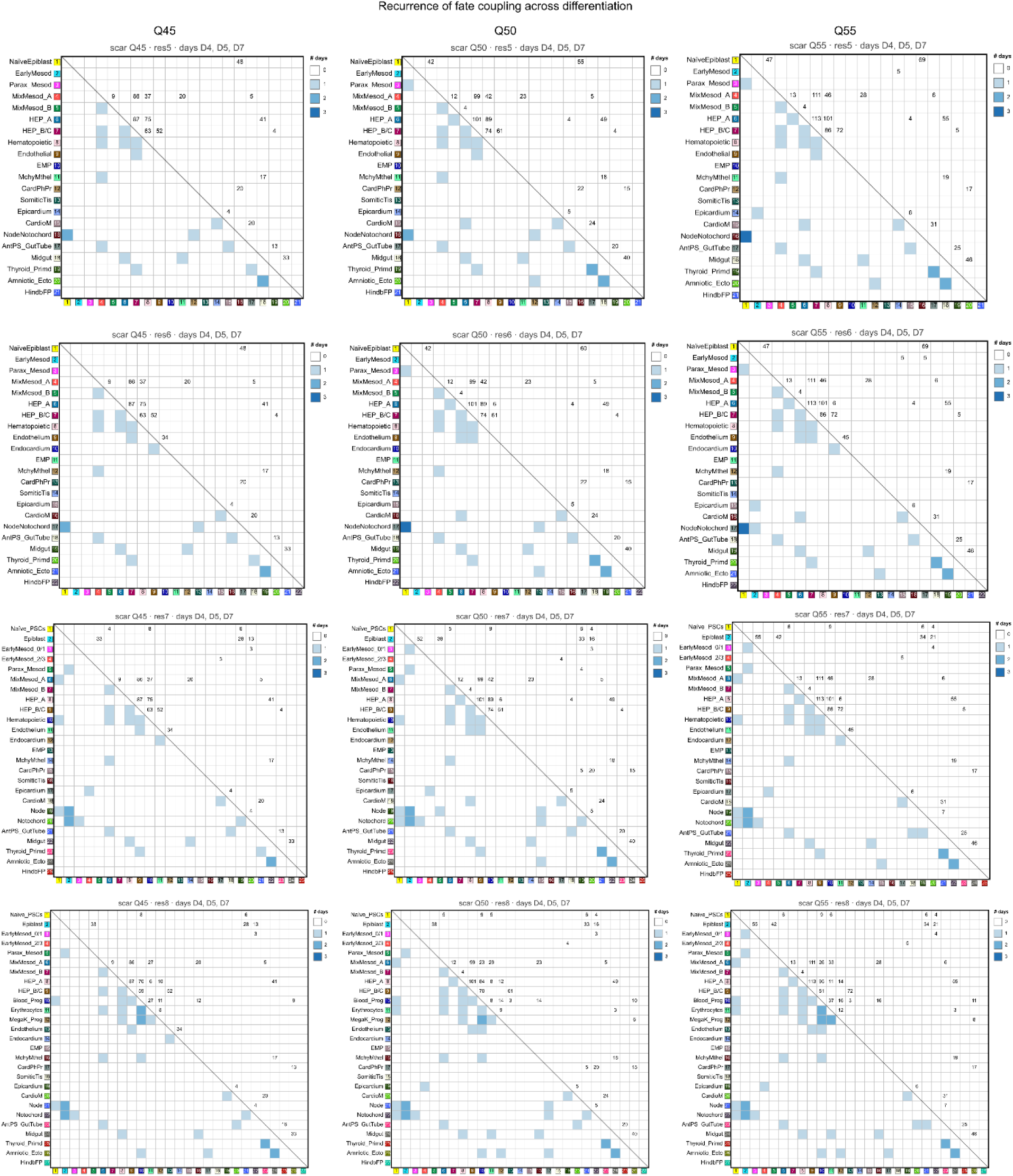
Cross-day recurrence of fate couplings across scar stringencies at high cell-type annotation resolutions (res5 - res8). As in Fig. S15, but with rows = res5, res6, res7, res8 (from top to bottom). Grid of 4 rows × 3 columns = 12 heatmaps. Columns, cell color, and annotation conventions match those of Fig. S15.

**Fig. S17.**
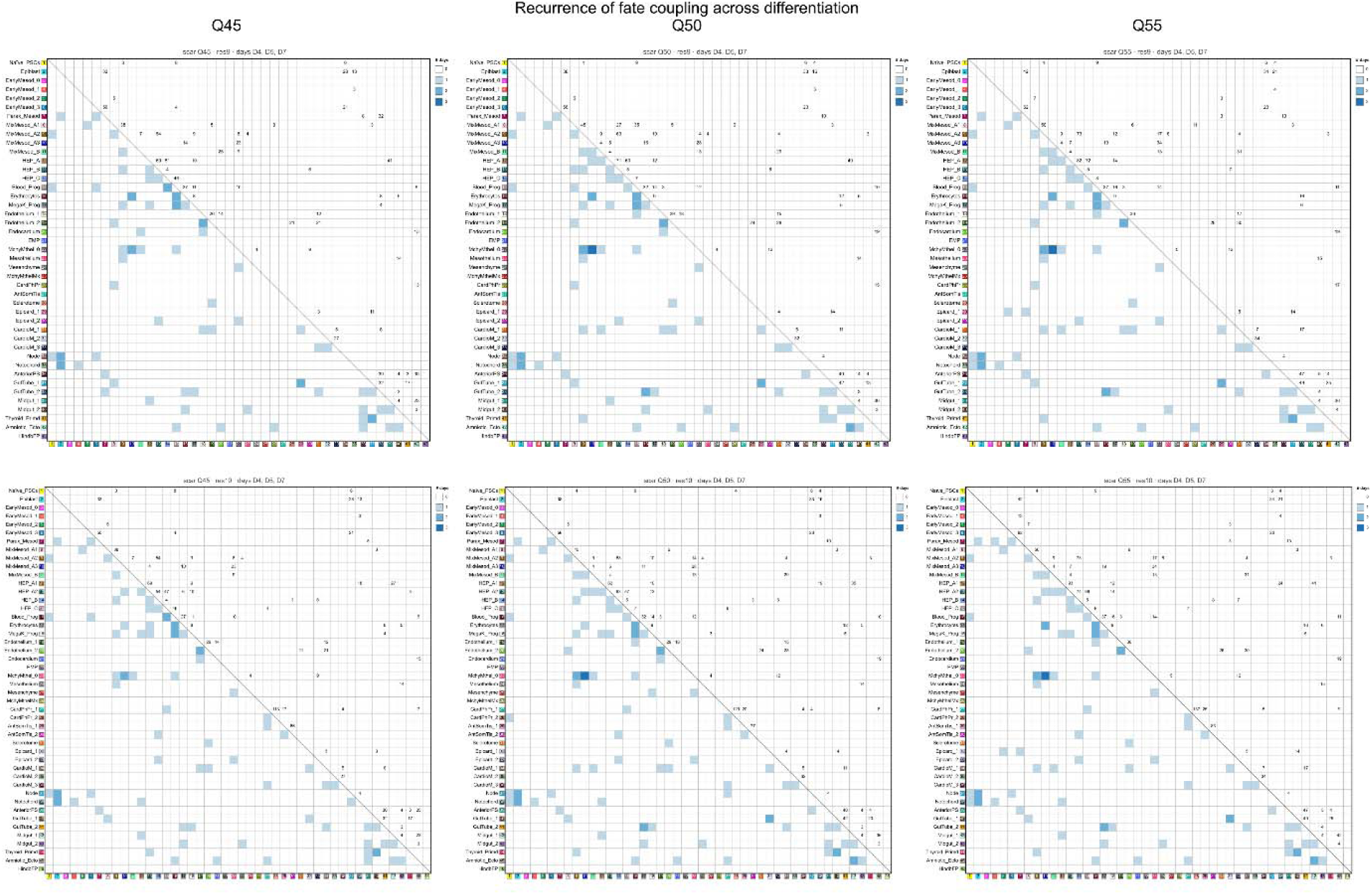
Cross-day recurrence of fate couplings across scar stringencies at the two finest cell-type annotation resolutions (res9 - res10). As in Fig. S15, but with rows = res9, res10 (from top to bottom). Grid of 2 rows × 3 columns = 6 heatmaps. Columns, cell color, and annotation conventions match those of Fig. S15.

**Fig. S18.**
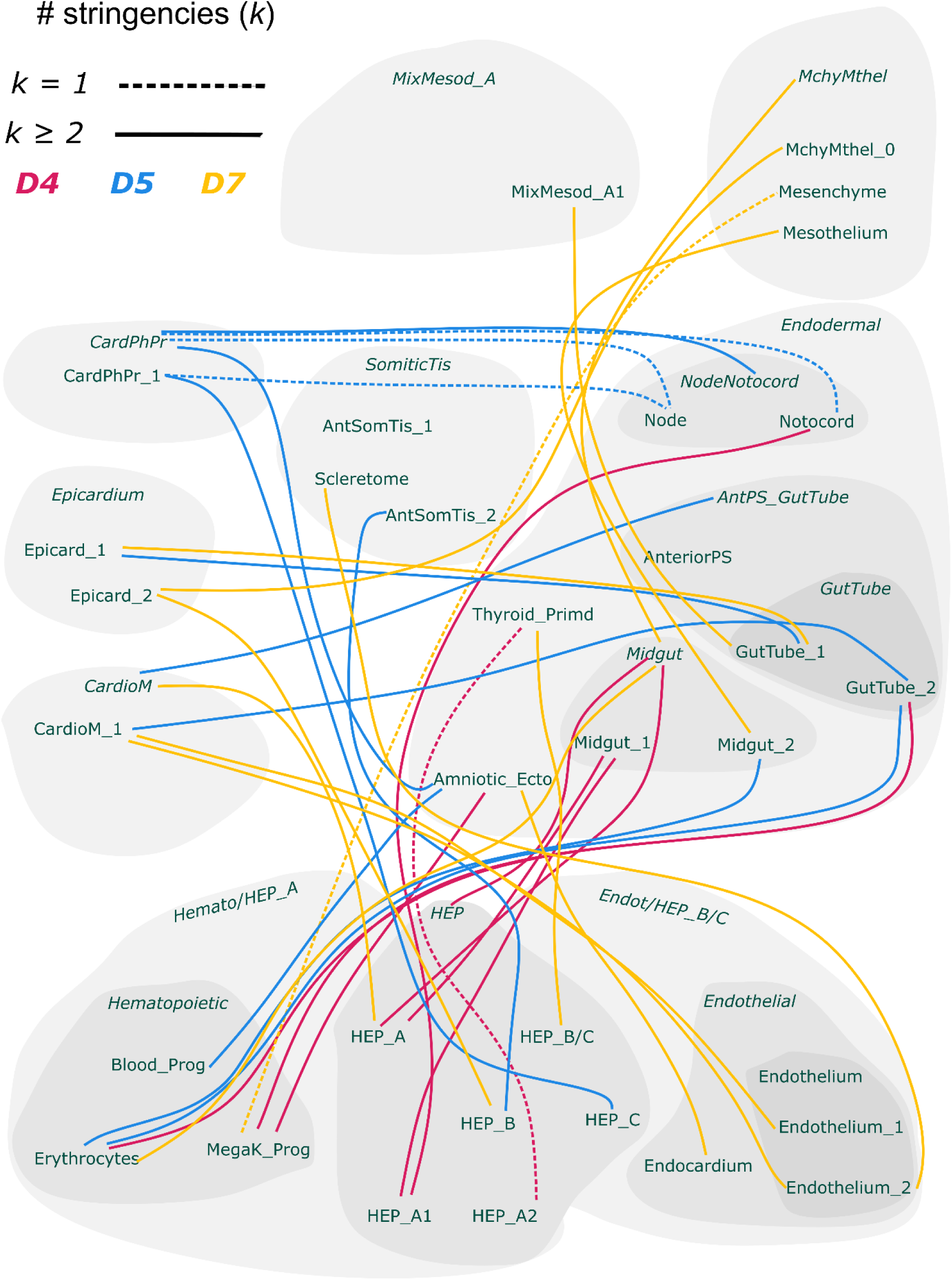
Schematic of robust lineage couplings in cross-trajectories. Schematic of robust lineage couplings. Edges = pairs significant at ≥ 1 of Q45 / Q50 / Q55, pooled across all 11 resolutions (res1 - res10). Edge color = day (D4 / D5 / D7). Edge style: solid = k > 1 (≥ 2 stringencies); dashed = k = 1.

## Supplementary Tables

**Table S1. CARLIN single-cell lineage-tracing results and comparison with transcriptome read counts.**

Lineage-tracing output from the CARLIN lineage library and its relationship to matched single-cell transcriptome data. (a) Cell-to-lineage assignments, mapping each detected cell to its lineage barcode recovered from the CARLIN library. (b) Per-cell scar counts derived from the CARLIN library. (c) Comparison of transcriptome counts with lineage counts for all cells detected in the CARLIN lineage library, including cells carrying wild-type (unedited) barcodes. (d) The same comparison restricted to cells whose barcodes contain indels (edited/scarred barcodes), excluding the wild-type barcodes.

**Table S2. Genetic elements and their sequences, as used for developing scRIPT.**

Sequences of the genetic components and oligonucleotides used to construct and validate the scRIPT lineage-tracing system. (a) Genetic elements of the transgene cassette targeted to the mouse *Tigre* locus, listed in 5′→3′ order with feature name, length and nucleotide sequence: the 5′ and 3′ homology arms (Tigre_5′HA, 3′HA; 1,000 bp-long each), the *EF1*α core promoter, eGFP, the modified destabilization domain d2 (destabilized by residues 422-461 of mouse ornithine decarboxylase), the CARLIN target array (TA), the *bGH* (bovine growth hormone) gene poly(A) signal-containing fragment, and the intervening linker/gap sequences (fully assembled insert; 3,974 bp-long); the capture-primer sequence and the gRNA-target (insertion) sequence are also given. (b) Primers used for cell-line generation and testing, each with its name, sequence, and use or binding location, grouped as used for genotyping/screening, 5′-homology-arm, transgene-body, and 3′-homology-arm primers. (c) Primers used to amplify the lineage (scar) library for sequencing, each with its name, sequence, and binding site/function.

**Table S3. Per-cell transcriptome QC, scar assignment, and doublet scores prior to filtering.**

Cell-level table for the 39,381 cells of the assignment-stage transcriptome that received a lineage-barcode (scar) assignment, before doublet removal and downstream filtering; each row is one cell (condition encoded in the Cell_ID suffix). The table integrates scar/lineage-barcode assignment, standard single-cell transcriptome quality metrics, and doublet scores, and it provides the inputs used for scDblFinder doublet calling and scar-entropy calculation.

Columns comprise: Cell_ID, cell barcode with sample/condition suffix; the assigned lineage scar and its supporting counts (scar, the scar/CIGAR string; most_common_scar_n, matching_scar_n, distinct_scar_n; total, long and short read counts n_total/n_long/n_short; fraction_common; cardinality_distinct, cardinality_matching; uniquely- and ambiguously-aligned read counts n_unique_alignments/n_ambiguous_alignments; the per-alignment-tier summaries X0/X1/X2_total_n, X0/X1/X2_cardinality and X1_total_frac); the scar-based doublet posterior (scar_posterior_doublet); standard transcriptome QC metrics (nCount_RNA, total UMIs; nFeature_RNA, detected genes; percent.mt, mitochondrial read percentage; percent.ribo, ribosomal read percentage); scDblFinder doublet scores and singlet/doublet classifications from three runs (scDblFinder_score/_class, scDblFinder_score_new/_class_new, scDblFinder_score_lv2/_class_lv2); and scar_entropy, the Shannon-entropy-based diversity score of the cell’s scar.

**Table S4. Cluster marker genes across annotation resolutions.**

Upregulated marker genes identifying each cell cluster at eleven levels of annotation resolution (res1, res2, res3, res4a, res4b, res5, res6, res7, res8, res9 and res10) of the integrated embryoid-body dataset (25,562 cells pooled from differentiation days D4, D5 and D7). The workbook contains one sheet per resolution level, each listing the top 100 marker genes per cluster. Markers were identified with Seurat FindAllMarkers using a Wilcoxon rank-sum test on the log-normalized RNA assay, testing only upregulated genes (only.pos = TRUE) detected in at least 10% of cells in either the cluster or the remaining cells (min.pct = 0.1), with a minimum average log2 fold-change of 0.25 (logfc.threshold = 0.25). Genes were retained at adjusted P < 0.01 (Bonferroni correction across all genes in the dataset), and for each cluster the 100 genes with the largest average log2 fold-change are reported. Each row is one gene in one cluster (a gene that marks several clusters appears once per cluster). Columns: cluster, cluster identity at that resolution; gene, gene symbol; avg_log2FC, average log2 fold-change of expression in the cluster relative to all other cells; pct.1, fraction of cells within the cluster expressing the gene; pct.2, fraction of cells outside the cluster expressing the gene; p_val, unadjusted P value; p_val_adj, Bonferroni-adjusted P value. Clusters with fewer than 100 genes meeting the significance threshold are reported in full.

**Table S5. Per-cell metadata for the integrated D4/D5/D7 embryoid-body Seurat object.**

Cell-level metadata for all 25,562 cells of the Harmony-integrated embryoid-body Seurat object, spanning differentiation days D4, D5 and D7 (n = 11,016, 4,123 and 10,423 cells, respectively); each row is one cell. This is the QC-filtered, doublet-removed dataset used for the analyses in the manuscript. Lineage-scar fields are populated only for cells with a recovered scar; cells without a scar are left blank, and cells carrying an unedited (wild-type, “1WT”) barcode are labelled as such.

Columns cover cell and sample identity; the assigned lineage scar and its derived attributes (raw scar/CIGAR string, the per-stringency and uniquely-aligned scar calls, scar entropy and complexity, and indel-fragment counts); clone-size measures (cells sharing each scar overall, per condition, and per day: day0, day4, day5, day7); reference-based cell-type predictions at three levels of granularity (Seurat label transfer); and the dataset’s own cell-type annotations at eleven resolution levels (res1-res10, including res4a and res4b).

**Table S6. Pairwise clonal fate-coupling results across all scar-stringency × day × annotation-resolution partitions of the scRIPT dataset.**

Pairwise clonal fate-coupling scores, right- and left-tail permutation p-values, permutation z-scores, null-distribution summaries (mean and standard deviation), and clone-support counts for every (scar stringency × differentiation day × cell-type annotation resolution) partition. This covers 132 partitions: four scar stringencies (Q45, Q50, Q55, Q100) × three differentiation days (D4, D5, D7) × eleven annotation resolutions (res1-res10, including res4a and res4b). A cell-type pair is called significant when pvalue_greater < 0.05 and clone_support ≥ 3. Coupling scores and permutation p-values (n_perm = 10,000) were computed with CoSpar v0.5.0 (SW method).

Q100, the least-restrictive stringency, is included for transparency; as shown in Fig. 5 it inflates the coupling network with stringency-fragile pairs (∼40-65% more significant pairs than the project-grade stringencies Q45/Q50/Q55) and should be interpreted with caution.

The workbook has six sheets: a README, per-partition metadata (132 rows), pair-level results at Q50 (the primary stringency; 8,814 rows), pair-level results across all four stringencies (35,494 rows), the significant subset (1,603 rows), and the cell types present per partition (2,856 rows). Full column definitions and the row-sorting scheme are in the README, and the full pipeline is in the manuscript’s Methods.

**Table S7. Predicted fate-bias differential gene-expression results for the HEP bifurcation into Hematopoietic and Endothelial fates (Q50, D4+D5, res4b).**

Differential gene-expression results for the bifurcation of Hematoendothelial progenitors (HEP) into Hematopoietic and Endothelial fates. Expression was tested on the committed t0 progenitors only (n = 945 of 2,404 t0 HEP cells; 1,459 ambiguous cells excluded; 435 committed to Endothelial, 510 to Hematopoietic) with a two-sided Wilcoxon rank-sum test and Benjamini-Hochberg correction across 27,998 genes. A gene is significant when |log2FC| > 0.25 and pval_adj < 0.05; log2FC is signed so that positive values mark higher expression in Endothelial and negative values in Hematopoietic (fate_A = Endothelial, fate_B = Hematopoietic, alphabetical). Of the 27,998 genes, 2,258 are significant (1,167 up in Endothelial, 1,091 up in Hematopoietic); among the 1,797 mouse transcription factors tested (from a 1,860-symbol source list), 218 are significant (101 up in Endothelial, 117 up in Hematopoietic).

The workbook has six sheets: a README, single-row analysis metadata, all 27,998 tested genes, the significant genes, all tested transcription factors, and the significant transcription factors. Full column definitions are in the README, and full methods (committed-progenitor classification via the CoSpar transition map, the Wilcoxon/BH differential-expression pipeline, and the transcription-factor source list) are in the manuscript’s Methods.

**Table S8. Predicted fate-bias differential gene-expression results for the CardPhPr bifurcation into Cardiac/Epicardial and Somitic-tissue fates (Q50, D7, res1).**

Differential gene-expression results for the bifurcation of Cardiopharyngeal progenitors (CardPhPr) into Cardiac/Epicardial (CardioM/Epicd) and Somitic-tissue (SomiticTis) fates. Expression was tested on the committed t0 progenitors only (n = 636 of 1,962 t0 CardPhPr cells; 1,326 ambiguous cells excluded; 325 committed to CardioM/Epicd, 311 to SomiticTis) with a two-sided Wilcoxon rank-sum test and Benjamini-Hochberg correction across 27,998 genes. A gene is significant when |log2FC| > 0.25 and pval_adj < 0.05; log2FC is signed so that positive values mark higher expression in CardioM/Epicd and negative values in SomiticTis (fate_A = CardioM/Epicd, fate_B = SomiticTis, alphabetical). Of the 27,998 genes, 1,459 are significant (734 up in CardioM/Epicd, 725 up in SomiticTis); among the 1,797 mouse transcription factors tested (from a 1,860-symbol source list), 161 are significant (45 up in CardioM/Epicd, 116 up in SomiticTis).

The workbook has six sheets: a README, single-row analysis metadata, all 27,998 tested genes, the significant genes, all tested transcription factors, and the significant transcription factors. Full column definitions are in the README, and full methods (committed-progenitor classification via the CoSpar transition map, the Wilcoxon/BH differential-expression pipeline, and the transcription-factor source list) are in the manuscript’s Methods.

**Table S9. GEMLI memory-gene catalogue across scar-stringency, day, and cell-type annotation resolution.**

Full output of the GEMLI (Gene Expression Memory-based Lineage Inference; Eisele et al., 2024) memory-gene sweep on the scRIPT dataset. GEMLI was run on 1,128 partitions (scar stringency × differentiation day × cell-type annotation resolution × cell type; minimum lineage size 3 cells; Q45/Q50/Q55 = 333/374/421 partitions), with Q50 as the primary stringency. Genes were graded into confidence tiers by resolution-robustness (a gene is res-robust on a day if called at all eleven resolutions that day): Tier-0, called in ≥ 1 Q50 partition but not res-robust on any day (background; n = 6,051); Tier-1, res-robust on 1 of 3 days (n = 4,692); Tier-2, on 2 of 3 days (n = 1,590); Tier-3, on all 3 days (n = 222, inclusive of Tier-4); and Tier-4, the Tier-3 genes additionally res-robust across all three stringencies (n = 35). Per-tier imprinted-gene enrichment was tested by one-sided Fisher’s exact test (upper tail) against the Q50-tested universe (|U| = 14,838 genes; 84 of 107 cataloged mouse-imprinted genes present; background 0.566%), Bonferroni-corrected across the five tests (α = 0.010).

The workbook has six sheets: a README, a partition manifest (1,128 rows), a per-tier summary, the full Q50 memory-gene cataloged (12,555 genes), the 35-gene Tier-4 core, and the mouse-imprinted reference list (107 genes). Full column definitions are in the README, and full methods (sweep design, tier construction, and the enrichment test) are in the manuscript’s Methods.

**Table S10. Naïve_PSC clone-persistence DGE results: 28 significant genes (Q50, celltype_res8).**

Differential gene-expression (DGE) results for the Naïve_PSC clone-persistence contrast: 2 persistent clones (one D4, one D7; 19 cells) versus 13 non-persistent reference clones (7 D4, 6 D7; 21 cells) in the Naïve_PSC compartment at Q50 scar-stringency and celltype_res8 annotation (40 cells in total). Significance was assessed with a two-sided, cell-level Cohen’s *d* permutation test on log-normalized expression (10,000 label permutations) and Benjamini-Hochberg correction across the 10,884 detected genes. The 28 genes shown all pass BH q < 0.05 and a clone-level direction-consistency filter; 25 are up-regulated in persistent clones and 3 in non-persistent clones (*N4bp2*, *Ttc28*, *Mbp*). GEMLI cross-validation annotates each gene’s Q50 memory tier and recurrence: 26 of 28 are called memory in at least one Q50 partition, none reach Tier-3 or Tier-4, and none overlap the 107-gene mouse-imprinted reference list. The workbook has five sheets: a README, single-row analysis metadata, per-clone metadata (15 clones), the 28 significant genes with their statistics and annotations, and per-clone mean expression. Full column definitions are in the README, and full methods (clone selection, the permutation test and direction-consistency filter, robustness diagnostics, and the GEMLI cross-validation) are presented in the Methods section.

